# PSI photoinhibition and protection at low temperatures and in fluctuating light: An ecophysiological vista

**DOI:** 10.64898/2026.11.02.763831

**Authors:** Ko Takeuchi, Riichi Oguchi, Kazuhiko Sugimoto, Ichiro Terashima, Masaru Kono

## Abstract

In green leaves *in vivo*, photosystem I (PSI) was thought to be resistant to photoinhibition. In 1994, it was found that, in chilling-sensitive plants such as cucumber (*Cucumis sativus* L.), exposing leaves to moderate constant light at temperatures below 10 °C induces PSI photoinhibition with little effect on PSII. Since the beginning of this century, PSI photoinhibition induced by fluctuating light (alternating high and low light, FL) or repetitive saturating pulses (rSPs) has been reported in many species. In this review, we compare the causes and consequences of PSI photoinhibition in cucumber at chilling temperatures with those of PSI photoinhibition induced by FL or rSPs. Changes in chloroplasts during these photoinhibitory treatments are followed, focusing on where and how reactive oxygen species are produced and which parts of PSI are damaged. We also discuss the roles of two pathways of cyclic electron flow around PSI, the NDH and PGR5/PGRL1 pathways, particularly in suppression of PSI photoinhibition. Lastly, we discuss problems with methods for measuring PSI parameters and future research directions from an ecophysiological viewpoint.

## Introduction

Photosystem I (PSI) is a plastocyanin/ferredoxin photooxidoreductase. The core complex is formed as a pseudo-homodimer (PsaA and PsaB) having a reaction center chlorophyll dimer (Chl *a* and Chl *a*’). Each of PsaB and PsaA has cofactors, namely Trp, three Chl *a* molecules including (A_0_), and phylloquinone (A_1_). Unlike photosystem II (PSII), charge separation occurs on both sides of the pseudo-homodimer. The two routes merge at the iron sulfur center F_X_ sandwiched between the PsaA and PsaB proteins, while F_A_ and F_B_ are cofactors in the subunit PsaC protein (Rutherford et al. 2012). PSI reduces ferredoxin (Fd) and thereby provides reducing power for carbon assimilation and other metabolic processes.

Although there were pioneering studies showing photoinhibition of PSI *in vitro* (Inoue et al. 1986, 1989; Satoh 1970a, b, c), there were few studies showing PSI photoinhibition *in vivo* (Öquist et al. 1987; Powles 1984) until 1994. After the discovery that PSI in a chilling-sensitive plant, *Cucumis sativus* L. (cucumber), is readily damaged by moderate constant light when the leaf temperature is kept below 10 °C (Terashima et al. 1994), chilling-induced PSI photoinhibition was reported in a wide range of species originating from warm areas.

PSI photoinhibition can also be induced by fluctuating light (FL, alternating high light (HL) and low light (LL)) (for reviews, see Kono and Terashima 2014; Yamori 2016). Tikkanen et al. (2010) grew several mutant lines of *Arabidopsis thaliana* (L.) Heynh. in the FL alternating 500/50 µmol photons m^−2^ s^−1^ for 1 min/5 min for 12 h daytime (500/50 | 1 min/5 min, 12-h photoperiod) and found that the *pgr5* mutant (the mutant of *PROTON GRADIENT REGULATION 5*, a component of the PGR5 cyclic electron flow pathway around PSI, CEF-PSI) died. The lethal phenotype was due to PSI photoinhibition (Suorsa et al. 2012). PSI photoinhibition induced by FL occurs within much shorter periods, not only in the *pgr5* mutant but also in the wild type. Kono et al. (2014) treated leaves of *A. thaliana* with the FL from red LEDs (240/30 | 2 min/2 min | 42 min). It has been also known that treatment of the leaves with repetitive saturating pulses (rSPs) induces remarkable PSI photoinhibition in *A. thaliana* (Tsuyama and Kobayashi 2009), *Helianthus annuus* L. (sunflower, Sejima et al. 2014), and *Triticum aestivum* L. (wheat, Takagi et al. 2019; Zivcak et al. 2015). For example, Sejima et al. (2014) used the regime of (20,000/0 | 300 ms/9.7 s | 240 min). Distinction between the FL and rSPs is not clear. In this article, the FL using very strong short pulses (less than 1 s duration) are called rSPs according to Sonoike (2025). Because it matters whether the LL phase of FL or rSPs consists of darkness or nonzero low irradiance, we treat these as distinct lighting regimes in § 3.

Photodamage to PSI imposes high cost on plants because repair of photodamaged PSI or PSI *de novo* synthesis is slow and thereby PSI photoinhibition adversely affects plant growth (Amin et al. 2024; Kudoh and Sonoike 2002). To avoid photoinhibition, plants must tightly control the redox state of PSI by balancing electron flows into and out of PSI, and by dissipating excitation energy through safe charge recombination. This protection depends on several coordinated processes, including photosynthetic control and non-photochemical quenching (NPQ), both induced by acidification of the thylakoid lumen, regulation of photosystem II (PSII) activity, alternative electron flow pathways including the Asada (water to water) pathway and two CEF-PSI pathways, the state transitions, spillover of excitations from PSII to PSI, etc.

We should also pay attention to light spectra used in the photoinhibition studies. If the FL in natural environments exerted the same effect as the FL from red LEDs, many plants in the field would suffer from photoinhibition. However, in natural environments, it does not appear that many plants suffer from the PSI photoinhibition. In the presence of sufficient far-red light, PSI photoinhibition by the red FL is largely suppressed (Kono et al. 2017, 2020). From an ecophysiological viewpoint as well, the role of far-red light in PSI photoinhibition should be properly recognized.

In this article, we review recent progress in the PSI photoinhibition studies in the model chilling-sensitive plant, cucumber, and those using the FL or rSPs. Then, the mechanisms underlying PSI photoinhibition and its photoprotection are argued. In particular, the roles of two CEF-PSI pathways are discussed in detail. Angiosperms have two pathways of cyclic electron flow around PSI that transfer electrons from Fd to the PQ pool: the PGR5/PGRL1-dependent pathway (hereafter PGR5 pathway), which is sensitive to antimycin A, and the NDH-dependent pathway (hereafter NDH pathway), which is catalyzed by the chloroplast NADH dehydrogenase-like complex (Munekage et al. 2002, 2004; Shikanai 2007; Yamori and Shikanai 2016).

We also consider technical problems in measurements of PSI parameters. In stress physiology studies, researchers often expose plants to drastic and unrealistic stress conditions. Although plants respond to such treatments and we are fully aware that such studies often deepen our knowledge of plant biological machinery, the plant responses in such studies should be interpreted with caution. We also have to seek ecological realism, because plant biological machinery has been mostly shaped by natural selection occurring in natural environments. The contents of this review are as follows:

1. PSI photoinhibition in cucumber, the chilling-sensitive model plant.
2. PSI photoinhibition in the fluctuating light and repetitive saturating pulses.
3. Meta-analyses of PSI photoinhibition studies
4. How do plants protect PSI?
5. Roles of NDH and PGR5/PGRL1 in protection from PSI photoinhibition
6. ROS: their generation and scavenging
7. Roles of far-red light and allocation of excitation between PSI and PSII
8. Technical problems in quantification of PSI parameters.
9. Future directions in PSI photoinhibition research

## 1. PSI photoinhibition in cucumber, the chilling-sensitive model plant

### 1.1 Chilling stress inhibits photosystem I

In 1994, it was found that illuminating cucumber leaves with weak light at temperatures below 10 °C decreased the quantum yield of PSI with little effect on PSII (Sonoike and Terashima 1994; Terashima et al. 1994). This phenomenon was not observed under anaerobic conditions or in darkness, indicating that it is oxidative damage requiring both light and oxygen (Bukhov et al. 2004; Sonoike 1996b; Terashima et al. 1998). By 2002, studies using cucumber as a model had shown that chilling weakens association of CF_1_ of the thylakoid ATP synthase with the thylakoid membrane, thereby impairing ΔpH formation and photosynthetic control (Peeler and Naylor 1988; Terashima et al. 1989, 1991a, b); degrades the PSI core protein PsaB (Kudoh and Sonoike 2002; Sonoike 1996a; Sonoike and Terashima 1994; Sonoike et al. 1997); and damages the F_A_/F_B_ and F_X_ Fe–S clusters (Sonoike et al. 1995). More recently, Takeuchi, Ifuku, and their colleagues demonstrated that destabilization of the PSI–NDH supercomplex and PSI over-reduction occur at an early stage of light treatment at chilling temperatures in cucumber (Takeuchi et al. 2022, 2025a, 2026).

After its discovery in cucumber, chilling-induced PSI photoinhibition was reported in a wide range of species, including *Solanum lycopersicum* L. (tomato), *Oryza sativa* L. (rice), *Zea mays* L. (maize), *Gossypium hirsutum* L. (cotton), *Saccharum officinarum* L. (sugarcane), *Secale cereale* L. (rye) etc. (Chen et al. 2024; Govindachary et al. 2004; Ivanov et al. 1998; Kornyeyev et al. 2003; Takeuchi et al. 2026; Tjus et al. 1999; Zhang Y et al. 2025). Except for rye, these plants all originate from warm-climate regions.

### 1.2 How is PSI photoinhibited?

PSI photoinhibition is not an immediate consequence of exposing leaves to low temperatures. Instead, it develops gradually as chilling weakens the array of mechanisms that protect PSI. The key to understanding this process is to determine the sequence of events leading to PSI photoinhibition. The time course outlined below is based primarily on studies of cucumber leaves chilled at 4 °C under irradiances of 100–250 µmol photons m⁻^2^ s⁻^1^ (400-700 nm). The timing of individual events varies with plant species or varieties, leaf age, temperature, and the irradiance level.

#### Phase 1 (<30 min)

Immediately upon exposure to chilling treatment, the Calvin-Benson-Bassham (CBB) cycle, photorespiration, and other metabolic processes that consume reducing power downstream of PSI slow down (Zhou et al. 2007). As NADP⁺ regeneration slows, oxidation of Fd by ferredoxin–NADP⁺ reductase (FNR) becomes restricted. Irreversible PSI photoinhibition is not yet detectable, but transient over-reduction of the electron transfer chain can promote reactive oxygen species (ROS) formation. ROS generated within PSI are normally scavenged by the Asada cycle (water-water cycle) (Asada 1999; Miyake 2010; Miyake and Asada 1992; Miyake et al. 1998). At low temperatures, however, the activity of thylakoid-bound ascorbate peroxidase (APX) decreases markedly (Terashima et al. 1998), limiting the capacity for ROS scavenging. Insufficient scavenging during this early phase may contribute to the destabilization of the chloroplast H⁺-ATP synthase and PSI–NDH supercomplex observed in Phase 2.

In parallel with these early metabolic and redox responses, the physical state of the thylakoid membrane may also change. The phase transition temperature of polar lipids extracted from cucumber thylakoids is approximately 10 °C (Hodgson et al. 1987; Orr and Raison 1987, 1990). Notably, this transition temperature is strikingly close to the approximately 10 °C threshold below which photoinhibition occurs in intact cucumber leaves (Hodgson et al. 1987; Sonoike 2011; Terashima et al. 1994). Cucumber thylakoids also show lower lipid fluidity and a larger fraction of ‘motion-restricted lipids associated with proteins’ than thylakoids from *Pisum sativum* L., a chilling tolerant species (Li et al. 1990). Exposure to temperatures below the transition may therefore rapidly reduce local membrane fluidity and affect lipid–protein interactions involved in the regulation of photosynthetic electron transport and P700 oxidation. However, no study has directly examined both the membrane phase transition and PSI photoinhibition under the same experimental conditions. Thus, it remains unclear whether thylakoid lipid properties contribute to PSI photoinhibition in cucumber. Despite this uncertainty, changes in thylakoid membrane fluidity would contribute, at least in part, to the destabilization of the thylakoid ATP synthase and PSI–NDH supercomplex observed in Phase 2.

#### Phase 2 (30 min to 2 h)

Between 30 min and 2 h of chilling in the light, the CF_1_ sector of the thylakoid ATP synthase becomes more loosely associated with the thylakoid membrane, potentially weakening the coupling between proton translocation and ATP synthesis (see § 1.3). This impairs the formation and maintenance of ΔpH across the thylakoid membrane, thereby weakening both NPQ and photosynthetic control. Electron flow from PSII to PSI consequently continues even though the CBB cycle is suppressed. During the same period, NDH activity also declines (see § 1.3). By accepting electrons from reduced Fd and returning them to the PQ pool, NDH provides a route for Fd oxidation and can relieve acceptor-side over-reduction when photosynthetic control is weakened. Destabilization of the PSI–NDH supercomplex within the first hour of chilling, however, abolishes NDH activity and eliminates this route. Taken together, loose association of thylakoid ATP synthase and NDH inactivation allow the transient PSI over-reduction observed in Phase 1 to become enhanced.

#### Phase 3 (2–5 h)

At this stage, the P700 oxidation capacity substantially decreases, as indicated by decreased Y(ND), and reduced Fd accumulates. This persistent PSI acceptor side over-reduction accelerates ROS production within PSI and promotes irreversible PSI photoinhibition. The maximum amount of photooxidizable P700 (P_m_) decreases markedly while the maximum quantum yield of PSII photochemistry (F_v_/F_m_) is relatively well maintained, indicating preferential photoinhibition of PSI. Pronounced damage to the Fe–S clusters and degradation of core proteins such as PsaB are also observed. Once this stage is reached, subsequent plant growth is severely inhibited. With further chilling, the loss of PSI activity restricts electron transport downstream of PSII, eventually causing F_v_/F_m_ to decline markedly.

Taken together, chilling-induced PSI photoinhibition proceeds in the following sequence (Fig. 1): (1) reduced electron consumption by enzymatic reactions and a decrease in thylakoid membrane fluidity; (2) transient over-reduction of the PSI acceptor side and ROS production; (3) weakened association of CF_1_ of the ATP synthase with the thylakoid membrane and destabilization of the PSI–NDH supercomplex; (4) impairment of photosynthetic control and Fd oxidation capacity; and (5) persistent PSI over-reduction and damage to the Fe–S clusters and PSI reaction center.

**Fig. 1.**
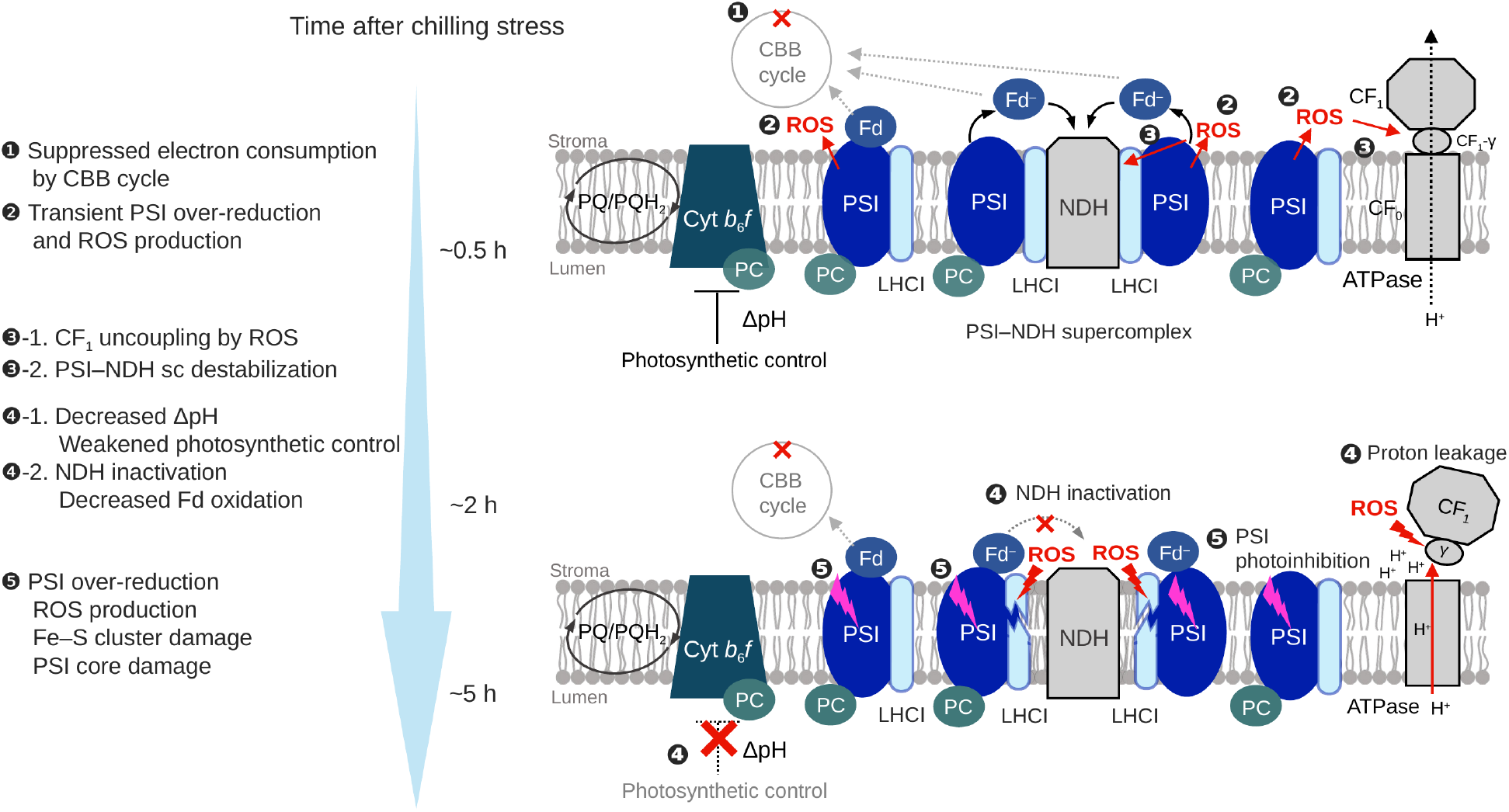
Temporal progression of chilling-induced PSI photoinhibition. Within the first 0.5 h of the onset of chilling stress, suppression of the CBB cycle and possible decrease in thylakoid membrane fluidity cause transient over-reduction of the photosynthetic electron transport chain and ROS production (1, 2) (Phase 1). These early redox changes and changes in membrane physics would weaken the association of CF1 with the thylakoid membrane, resulting in uncoupling of the chloroplast ATP synthase (3-1), as well as destabilization of the PSI–NDH supercomplex (3-2). Both changes occur within the first hour of chilling (upper panel). The weakened association of CF₁ allows H⁺ to leak from the lumen into the stroma, thereby dissipating ΔpH and weakening photosynthetic control at the Cyt b6fcomplex (4-1). At the same time, destabilization of the PSI–NDH supercomplex leaves NDH in an inactive monomeric form, reducing NDH-mediated Fd oxidation (4-2) (Phase 2). The combined impairment of photosynthetic control and Fd oxidation leads to a marked decline in P700 oxidation capacity between 2 and 5 h after the onset of chilling, accompanied by further ROS production in PSI and damage to the Fe–S clusters and PSI core proteins (5; lower panel) (Phase 3).

### 1.3 Dysfunction of protection mechanisms against chilling-induced PSI photoinhibition

#### Donor-side

Damage to the thylakoid ATP synthase is a key event in the early progression of chilling-induced PSI photoinhibition. When chilling-sensitive cucumber leaves were illuminated at low temperatures, the association of the catalytic portion (CF_1_) of ATP synthase with the membrane loosened, resulting in uncoupling of ATP synthesis (Terashima et al. 1989, 1991a, b, 1998). Accordingly, when thylakoids were isolated immediately after the treatment of the leaves at chilling temperatures in the light, substantial amounts of CF_1_ polypeptides were recovered in the supernatant. Such loose association would leave the CF_0_ proton channel open to the stroma, allowing H⁺ to leak from the lumen. This proton leakage impairs formation of the trans-thylakoid pH gradient (ΔpH), thereby weakening ΔpH-dependent NPQ and photosynthetic control at Cyt *b*_6_*f* complex (Kono et al. 2022; Takagi et al. 2017a). Indeed, when cucumber leaves were exposed to 4 °C and 100 µmol photons m⁻² s⁻¹, NPQ formation was markedly suppressed within 60 min after the onset of treatment (Fig. 2). Using the electrochromic shift (ECS) at 515 nm, Kono et al. (2022) showed that proton leakage associated with CF_1_ destabilization or dissociation is slower than protonophore-mediated proton transport but sufficiently rapid to impede ΔpH formation. Proton leakage was detectable after only 30 min of chilling and can therefore precede PSI photoinhibition.

**Fig. 2.**
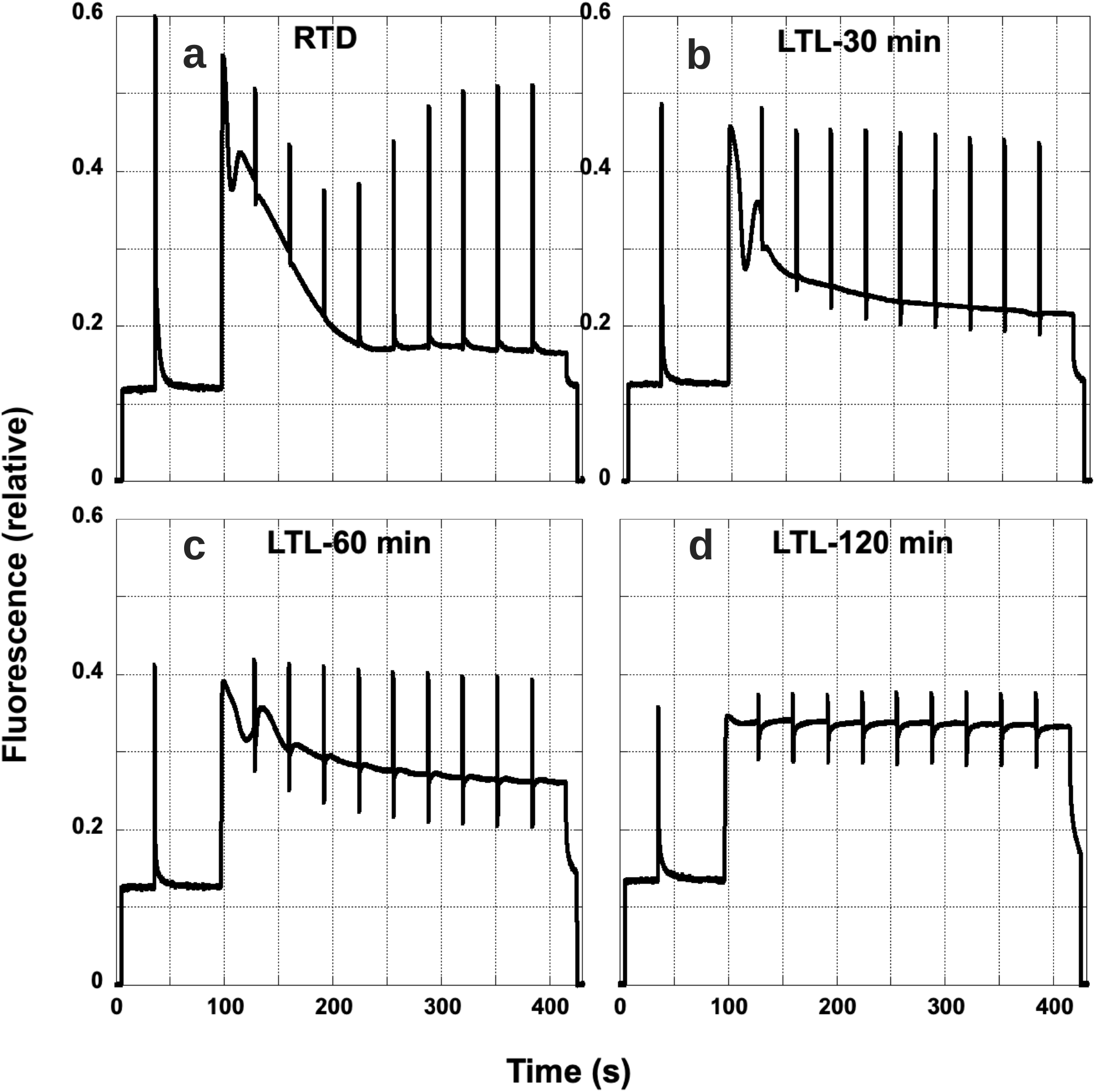
Quenching analysis curves in the cucumber leaf segments after the light-chilling treatment. Cucumber ‘Nanshin’ leaf segments were treated at 4 °C and an irradiance of 100 µmol photons m^-2^ s^-1^. a, control leaf segment kept in the dark at 23 °C (RTD); b, leaf segment treated at 4 °C in the light for 30 min (LTL-30 min); c, LTL-60 min; and d, LTL-120 min. Fluorescence was measured with a Dual-PAM (Walz, Germany) at 23 °C. The irradiance level of the actinic light (red LEDs at 635 nm) was 100 µmol photons m^-2^ s^-1^. As *F_m_*’ levels were nearly constant in c and d, development of NPQ was suppressed by the chilling treatment in the light for 60 to 120 min. This may be explained by the loose association of CF_1_ of ATP synthase with the thylakoid membrane. The quantum yield of PSII photochemistry, *Y(II)* = (*F*_m_’– *F*_s_’)/*F*_m_’, measured at 23 °C decreased from 0.68 (control) to 0.1 at 2 h with the increasing damage to PSI. Reproduced from Kono et al. (2022) with permission from the publisher.

#### Acceptor-side

A comparative study among cucumber cultivars revealed substantial variation in susceptibility to chilling-induced PSI photoinhibition (Takeuchi et al. 2022). Notably, a chilling-tolerant cucumber cultivar was relatively resistant to PSI photoinhibition despite a decrease in ΔpH (Takeuchi et al. 2025a). This observation points to an important role for acceptor-side pathways in protecting PSI when ΔpH-dependent control is weakened. Takeuchi et al. (2025a) identified NDH activity as the major difference on the PSI acceptor side. NDH is a large thylakoid membrane complex that accepts electrons from reduced Fd and reduces PQ (Ifuku et al. 2011; Shikanai 2016; Shikanai et al. 2025). In land plants, NDH associates with two PSI units through Lhca5 and Lhca6 to form the PSI–NDH supercomplex (Kato et al. 2018b, 2021; Otani et al. 2017, 2018). Fd binds to the subcomplex E (SubE) of the PSI–NDH supercomplex, and electrons pass through three internal Fe–S clusters before reaching PQ (Introini et al. 2025; Schuller et al. 2019; Shen et al. 2022; Su et al. 2022). The close association between PSI and NDH may facilitate electron transfer from Fd to NDH. Despite this structural arrangement, the physiological role of NDH in C_3_ plants remained unclear for many years because NDH-deficient mutants generally show no obvious phenotype under stable growth conditions. We discuss the functions of NDH in C_3_ plants in more detail in §§ 2 and 5.

In a chilling-sensitive cucumber cultivar, NDH activity was lost within 1 h of chilling treatment (Takeuchi et al. 2025a, 2026). This loss was not observed in leaves chilled in the dark, consistent with photooxidative inactivation of PSI–NDH supercomplex (Takeuchi et al. 2026). During chilling in the light, the PSI–NDH supercomplex was destabilized and NDH became monomeric. Because monomeric NDH is inactive (Kato et al. 2018a; Otani et al. 2018), maintenance of the PSI–NDH supercomplex during chilling is important for limiting PSI over-reduction and subsequent photoinhibition. It has been further shown that an NDH-deficient rice mutant was more susceptible to chilling-induced PSI photoinhibition than the wild type (Takeuchi et al. 2026). Taken together, these results show that NDH protects PSI in C_3_ plants at low temperatures. It is noteworthy that Terashima, Sonoike, and colleagues luckily chose a very chilling-sensitive cultivar ‘Nanshin’ for their studies started in 1990s.

How does NDH prevent over-reduction of the PSI acceptor side? NDH accepts and transfers electrons to the PQ pool. At the same time, NDH pumps protons into the lumen (Burnap 2026; Miller et al. 2021; Richardson et al. 2021; Strand et al. 2017). ΔpH thus formed would strengthen photosynthetic control. A highly reduced PQ pool would restrict electron transfer by the reduction-induced suppression of electron flow (RISE) (Shaku et al. 2016; Shimakawa et al. 2018; Tikhonov 2024). NDH contains three internal Fe–S clusters that may act as transient electron sinks and thereby shorten the lifetime of reduced states on the PSI acceptor side (Richardson et al. 2021). The physiological importance of NDH during chilling is supported by the reports that NDH abundance increases under chilling in the light (Teicher et al. 2000) and that reduced NDH activity in maize is associated with lower chilling tolerance (Urzinger *et al*. 2025). However, large ΔpH and highly reduced PQ pool could suppress NDH activity (Strand et al. 2017). We will discuss regulation of NDH activity in §§ 5.1 and 5.3.

The importance of relieving acceptor-side reduction is further illustrated by flavodiiron proteins (FLVs). FLV provides a terminal electron sink by transferring electrons to O_2_ and reducing it to water. Rice plants expressing FLV are highly resistant to chilling-induced PSI photoinhibition (Takeuchi et al. 2026). Conifers such as *Pinus sylvestris* L. (Scots pine) and *Picea abies* (L.) Karst. (Norway spruce) have lost chloroplast NDH during evolution and instead rely on FLV to mitigate early-spring chilling damage (Bag et al. 2023; Wakasugi et al. 1994). These observations suggest that NDH and FLV provide biochemically distinct but functionally overlapping routes for protecting PSI during chilling stress. Concerning another CEF-PSI pathway involving PGR5/PGRL1, which is widely regarded as more active than NDH in C₃ plants (Yamori and Shikanai 2016), information on its role in PSI photoinhibition at low temperatures is scarce.

## 2. PSI photoinhibition in the fluctuating light and repetitive saturating pulses

Fluctuating light (FL) and repetitive saturating pulses (rSPs) have been used to clarify the mechanisms underlying PSI photoinhibition and the mechanisms that protect PSI against photoinhibition. PGR5- and NDH-pathways both generate proton motive force (pmf) without net production of NADPH, and both have been linked to the protection of PSI from photoinhibition. However, there are contrasting views on which pathway is more important.

In the paper that first reported the *A. thaliana pgr5* mutant, the plants grown at 50 µmol photons m⁻² s⁻¹ were used. When the plants were exposed to a constant HL of 1,500 µmol photons m⁻² s⁻¹ for 10 min, up to 60% of PSI was damaged in *pgr5*, whereas less than 10% was damaged in WT (Munekage et al. 2002). This pioneering study showed that PSI in the *pgr5* mutant is susceptible to photoinhibition, although the *pgr5* mutant does not show a marked phenotype when grown in constant light.

When *A. thaliana pgr5* seedlings were grown in white fluctuating light (FL) alternating between 500 µmol photons m⁻² s⁻¹ for 1 min and 50 µmol photons m⁻² s⁻¹ for 5 min (500/50 | 1 min/5 min), they died due to PSI photoinhibition, whereas the WT and *ndho*, an NDH mutant, showed no visible phenotype (Suorsa et al. 2012). PSI photoinhibition occurs in milder FL treatments for much shorter periods. In the red FL for only 42 min (240/30 | 2 min/2 min | 42 min), P_m_ decreased to 89% in the wild type (WT) and 62% in *pgr5*, whereas in *crr2-2*, an NDH mutant, the decrease in P_m_ was comparable to that in WT (Kono et al. 2014). In the harsher red FL (1,200/30 | 2 min/2 min | 162 min), P_m_ decreased to 70% in WT, 20% in *pgr5* and 60% in *crr2-2* (Kono and Terashima 2016). These results clearly indicate that PSI is readily photoinhibited by FL and that PGR5 is the main actor in suppression of PSI photoinhibition. When *Oryza sativa* L. WT and mutants grown at 28 °C were treated with the FL (1,500/200 | 10 min/10 min | 5 h), P_m_ decreased to 81%, 61% and 33% of the initial levels in WT, *crr6* (an NDH mutant) and PGR5-knockdown lines, respectively (Yamori et al. 2016), indicating that PGR5 contributes to protection much more than NDH.

When far-red light (FR, peaking at 740 nm, 25.6 W m⁻²) was added to both HL and LL phases of the red FL regime (1,200/30 | 800 ms/10 s | 120 min), the conclusion reversed. Without FR, P_m_ declined to 70% in all genotypes, whereas with FR in both phases, P_m_ in WT and *pgr5* remained at 97% and 92% of the initial level, and P_m_ in *crr2-2* and *crr4-3*, NDH mutants, declined to 80% (Kono et al. 2017). With FR, therefore, PSI protection depended on the NDH pathway rather than on the PGR5 pathway. Similar results were obtained in the tobacco (*Nicotiana tabacum* L.) *ndhB*-disrupted line and in the rice *crr6* line in the FL with a HL phase of 2,000 µmol photons m⁻² s⁻¹: FR protected PSI in WT, but the protection was weaker in the NDH-deficient lines (M. Kono, unpublished data).

In rice lines grown at a low temperature of 20 °C, the NDH-deficient mutant *crr6* showed lower ETR(I), CO₂ assimilation rate and relative growth rate than WT. When grown at 28 or 35 °C, the growth of *crr6* was similar to that of WT, but ETR(I) and CO₂ assimilation rate were still lower below the growth irradiance, 800 µmol photons m⁻² s⁻¹ (Yamori et al. 2011). A recent comprehensive study by Kodama and Yamori (2026) using rice WT and *crr6*, grown in the field, showed that NDH deficiency markedly decreases ETR(I) and CO₂ assimilation rate, at low temperatures and/or in low irradiances. In the liverwort *Marchantia polymorpha* L., disruption of *ndhB* increased the reduction state of the PQ pool at low irradiances (Ueda et al. 2012), probably due to PSI acceptor-side limitation.

As already discussed, NDH plays a crucial role in the protection of PSI at low temperatures (§ 1). Taken together, NDH is important at low temperatures, at low irradiances, and in FL in the presence of FR, whereas PGR5 functions in ordinary HL and in FL of high irradiances with no FR supplementary light.

## 3. Meta-analyses of PSI photoinhibition studies

The number of studies on PSI photoinhibition has been increasing since the 1990s. Recently, studies examining the effects of fluctuating-light or intermittent strong light, and those using the mutants related to CEF-PSI have been increasing. These studies have used various plant species. To conduct meta-analysis and discuss the effects of photoinhibitory conditions on the rate of PSI photoinhibition, we collected data from 107 experiments from 32 papers (Table S1). In photoinhibition studies, the extent of photoinhibition is related both to the photon-exposure (fluence) and to the irradiance level (fluence rate) (Kornyeyev et al. 2003; Oguchi et al. 2021), although this approach may not be simply applicable to the FL or rSPs regimes. Since the repair process of PSI is slow, we may neglect the repair process. Using the equation proposed by Kok (1956), the rate of PSI photoinhibition is described by a simple differential equation:

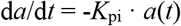

where *a*(*t*) is the active fraction of PSI reaction centers and *K*_pi_ is the rate constant for PSI photoinhibition. This is readily solved:

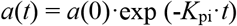

With *I* and X, denoting irradiance (fluence rate) in mol m^-2^ s^-1^ and photon-exposure (fluence) in mol m^-2^, respectively, the solution can be expressed as:

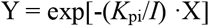

where Y is the ratio of PSI activity at the end of photoinhibitory treatment to that before the treatment. The ratio of P_m_ after the photoinhibition treatment to that before the treatment can also be used as Y. *K*_pi_ /*I* in mol^-1^ m^2^ is the coefficient of PSI photoinhibition, which is a useful parameter to consider the effect of irradiance and fluence (Table S1; Oguchi et al. 2009; Tyystjärvi and Aro 1996).

Light at very high irradiances would give damage to PSI in a manner different from light at moderate irradiances (see § 6; Shimakawa et al. 2024; Sonoike 2025), thus we divided the categories of the FL and intermittent pulses based on whether the irradiance was below or above 10,000 µmol photons m^-2^ s^-1^. It has also been suggested that the LL phase of the FL regime affects the extent of PSI photoinhibition, particularly depending on whether this phase is darkness or low light (Kono et al. 2017; Sejima et al. 2014). Therefore, we also divided the category of FL and flashes depending on whether the LL phase was actual LL or darkness. Intermittent HL/pulses denote that the HLs or saturating pulses were given with no background light. We group the experimental conditions into 8 categories.

1. isolated thylakoids, constant light
2. leaves, at chilling temperatures, constant light
3. leaves, at moderate temperatures, constant light
4. leaves of *A. thaliana pgr5*, at moderate temperatures, constant light
5. leaves, FL (< 10,000 µmol photons m^-2^ s^-1^)
6. leaves, intermittent HL/pulses (< 10,000 µmol photons m^-2^ s^-1^)
7. leaves, FL (≥ 10,000 µmol photons m^-2^ s^-1^)
8. leaves, intermittent HL/pulses (≥ 10,000 µmol photons m^-2^ s^-1^)

In early studies of PSI photoinhibition, isolated thylakoids were usually treated with constant light. Although *K*_pi_/*I* varied widely among these studies, its mean value was higher than that obtained for leaves exposed to constant light at chilling temperatures (1 *vs*. 2 in Fig. 3a). The *K*_pi_/*I* of the leaves photoinhibited at room temperature in constant light was very low, although not zero, and was significantly lower than that of leaves photoinhibited at chilling temperatures (3 *vs*. 2). Even in the constant light, *K*_pi_/*I* in the leaves of *pgr5* (the mutant of PGR5/PGRL1 CEF-PSI) were higher than WT plants (3 *vs*. 4, because the number of reports with *pgr5* mutant is small, the difference is not statistically significant). In these experiments, *pgr5-1* was used. It should be noted that *pgr5-1* also carries another mutation, which enhances the phenotype (Wada et al. 2021).

**Fig. 3.**
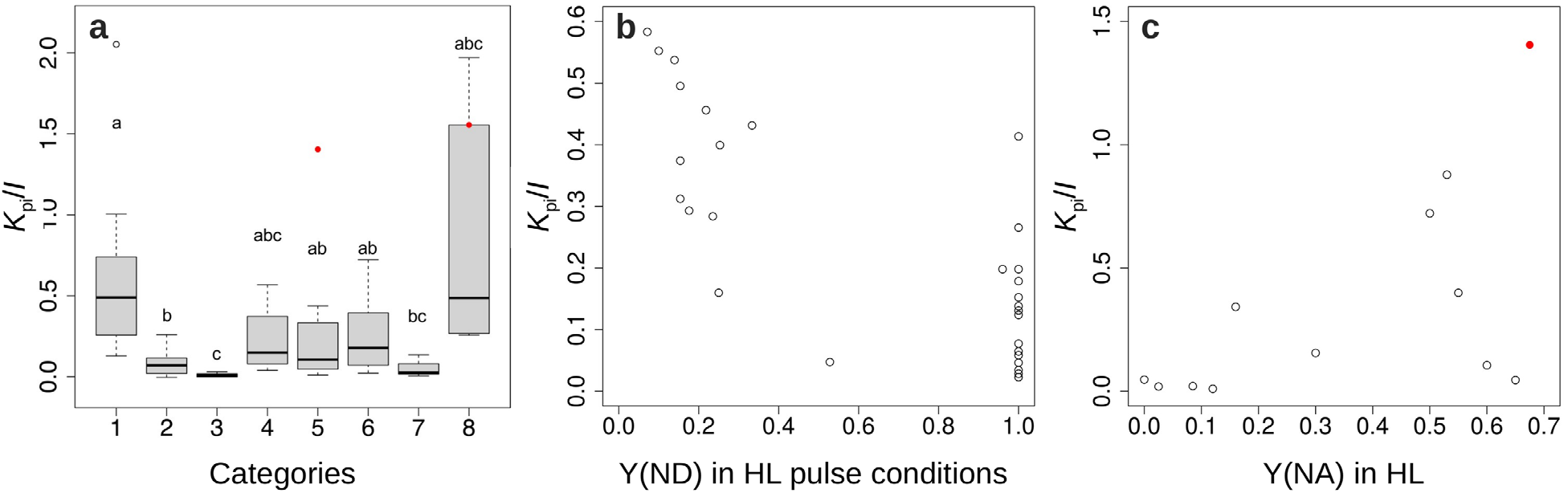
Boxplot of *K*_pi_/*I* (a) and scatter plots between Y(ND) and *K*_pi_/*I* (b) and between Y(NA) and *K*_pi_/*I* (c) produced from meta-analysis of recent PSI photoinhibition studies. Categories from 1 to 8 in panel (a) indicate different types of photoinhibition environments as follows: 1, 2, 3 and 4 indicate the data gained under steady-light conditions using isolated thylakoids, leaves in chilling condition, leaves in moderate temperature, and leaves of *A*. *thaliana pgr5* in moderate temperature, respectively. 5, 6, 7 and 8 indicate the data with leaves in fluctuating light condition, leaves in intermittent light condition, leaves in fluctuating light condition with very strong light and leaves in intermittent light condition with very strong light, respectively. We classified treatments with an irradiance of 10,000 µmol photons m⁻² s⁻¹ or higher during the high-light phase as very-strong-light treatments. The data of *A*. *thaliana pgr5* obtained under FL or intermittent light are not included in the boxes of categories 5 and 8 but are shown as red points. Different letters indicate statistically significant differences among the categories according to the Games-Howell test. In panel (b), Y(ND) was calculated as proportion of P700^+^ at 1 s from the start of high-light phase in the fluctuating-light conditions to the maximum P700^+^. In the case of Takagi et al. 2017b, the maximum P700^+^ was measured under pulse light (5,000 µmol photons m^-2^ s^-1^) condition without far-red light. In the case of Kono et al. 2017, Y(ND) was calculated with the data measured at 100th high-light phase, in which we considered the PSI photoinhibition was still small. In panel (c), Y(NA) was calculated as (P_m_ – P_m_’)/P_m_, where P_m_ means total P700 measured under saturating pulse with far-red light and P_m_’ means the P700^+^ measured under saturating pulse at a steady state of high-light phase in the fluctuating-light conditions. We should note that the timing of the measurement was different in panel (b): at 1 s and in panel (c): at a steady state. In panel (b) and (c), the data of references in categories from 1 to 4 and references in which Y(ND) or Y(NA) was measured at a light intensity different from that of the high-light phase were not included.

When the irradiance in the HL was lower than 10,000 µmol photons m^-2^ s^-1^, the *K*_pi_/*I* of leaf samples was significantly higher than that of leaf samples photoinhibited in constant light, both in the FL and intermittent HL/pulse regimes (3 vs. 5, and 3 vs. 6). The difference between the FL and intermittent HL/pulse regimes was not clear (5 *vs*. 6). When the irradiance of HL/pulse was 10,000 µmol photons m^-2^ s^-1^ or higher, the *K*_pi_/*I* of leaves in the FL regime with actual LL was not higher than that of leaves in constant light, whereas leaves in the intermittent HL/pulse showed higher but highly variable *K*_pi_/*I* values (category 3 *vs*. 7, and category 3 *vs*. 8). The large variation among the reported values and the limited number of data points in very HL/pulse may have prevented the detection of a statistically significant difference. To see the effect of very high HL/pulse on *K*_pi_/*I* of PSI, increment of the number of future studies under the same conditions would be necessary. We should note that the *K*_pi_/*I* and the extent of PSI photoinhibition can be also affected by differences in leaf traits, including leaf thickness and chlorophyll content per leaf area, among material species and growth environments, as the extent of PSII photoinhibition shows a gradient in a leaf and is influenced by leaf thickness and Chl content per leaf area (Oguchi et al. 2011; Pätsikkä et al. 2002).

As shown in the boxplot, *K*_pi_/*I* of PSI varies considerably among studies conducted in the FL or intermittent HL/pulse regime, which may be attributable to differences in the redox state around PSI during the exposure. The variation of redox state around PSI can be caused by different conditions among the photoinhibition experiments and by different materials. As already mentioned, suppression of the electron flow from PSII to PSI and the increment of the proportion of oxidized P700 (P700^+^) are important to reduce the extent of PSI photoinhibition (Shimakawa and Miyake 2018; Sonoike 2011; Sonoike and Terashima 1994; Terashima et al. 1994; Trissl 1997), therefore we plotted the relationship between the proportion of P700^+^ to total P700 (Y(ND)) and the *K*_pi_/*I* of PSI, using data from the meta-analysis (Fig. 3b). We only used the data from reports, which measured P700^+^ during the HL phase of fluctuating or intermittent-light conditions. The negative correlation between Y(ND) and the *K*_pi_/*I* of PSI would support the suggestion that keeping higher Y(ND) in fluctuating-light conditions is important to reduce the extent of PSI photoinhibition.

In his recent review, Sonoike (2025) claims that, although an increase in [P700^+^] or Y(ND) is a necessary condition for protection of PSI, oxidation of the acceptor side, or the decrease in Y(NA), is essential for PSI protection (see §§ 4 and 5). Although the studies in which Y(NA) was measured during the HL phase are few, we plotted such data (Fig. 3c). Although the data points are not many, we may see the trend indicating that *K*_pi_/*I* is inversely related to Y(NA).

A recent paper by Tanigawa et al. (2026) is worth mentioning here. They used *A. thaliana* overexpression lines of PGR5. When they grew these lines in a moderate FL regime, growth of the 2X over-expressor outcompeted WT and the over-expressors with more PGR5, indicating that maintaining an appropriate level of PGR5 is important for efficient photosynthetic regulation in white FL.

## 4. How do plants protect PSI?

Studies in cucumber have revealed how chilling-induced PSI photoinhibition develops. Chilling-tolerant plants, by contrast, retain mechanisms that limit electron flow into PSI and relieve over-reduction of its acceptor side (Kono et al. 2022; Miyake 2020). How plants protect PSI from photoinhibition induced by chilling temperatures, FL and rSPs is discussed below.

### 4.1 Photosynthetic control

The decreased capacity to regenerate NADP⁺ and the decline in ATP consumption under chilling stress lead to acidification of the thylakoid lumen. This restricts PQH_2_ oxidation at the Cyt *b*_6_*f* complex and thereby suppresses electron flow into PSI. This ΔpH-dependent restriction of electron flow at the Cyt *b*_6_*f* complex is known as the photosynthetic control (see Degen and Johnson 2024 for a recent review). Lumen acidification also promotes NPQ, further decreasing electron flow into PSI. In cucumber, however, uncoupling of the thylakoid ATP synthase dissipates ΔpH and weakens both photosynthetic control and NPQ.

### 4.2 Suppression of the electron flow to PSI

PSII has a greater repair capacity than PSI. Under various environmental stresses, PSII photoinhibition occurs first, limiting electron flow from PSII to PSI and protecting PSI (Ivanov et al. 1998; Kim et al. 2005; Lima-Melo et al. 2019; Messant et al. 2024; Napaumpaiporn et al. 2024; Obara et al. 2023; Ozawa et al. 2024; Sonoike 1996b; Takeuchi et al. 2025b; Tikkanen et al. 2014; Zhang Z et al. 2016). As has been discussed in § 1.2, this is not the case in cucumber. PSII photoinhibition appears to be a sacrificial defense, although it is difficult to prove. In the chilling-tolerant plant *A. thaliana,* exposure to chilling in the light induced PSII photoinhibition, as indicated by a decrease in F_v_/F_m_, whereas P_m_ was maintained. By varying the duration of chilling treatment, Takeuchi et al. (2025b) obtained *Arabidopsis* plants with F_v_/F_m_ values ranging from approximately 0.2 to 0.8 and subsequently exposed them to rSPs that induce PSI photoinhibition. Plants with lower initial F_v_/F_m_ maintained P700 in a more oxidized state during the SPs and showed a slower decline in PSI activity. F_v_/F_m_ before the pulse treatment was strongly correlated with susceptibility to PSI photoinhibition (*r* = 0.97) (Takeuchi et al. 2025b). Thus, PSII photoinhibition protects PSI. Even in the plants in which PSII photoinhibition maintained P700 in an oxidized state, however, Fd became reduced during the SPs. Steady-state oxidation of P700 alone therefore cannot completely prevent acceptor-side reduction during intense excitation. Nevertheless, reoxidation of Fd after each short pulse was faster in plants that had undergone PSII photoinhibition. Faster Fd reoxidation indicates that electrons were removed more efficiently from the PSI acceptor side in plants in which PSII was photoinhibited. By keeping the PQ pool relatively oxidized, PSII photoinhibition may also favor CEF-PSI and thereby contribute to rapid Fd reoxidation (Allen 2003; Satoh et al. 2025).

We have dealt with several examples in § 2. However, is susceptibility of PSI to red FL a general phenomenon? PSI photoinhibition was indeed readily induced by FL from red LEDs in most of the plants we tested (Kono et al. 2017), whereas *Alocasia odora* (Lodd.) Spach, a shade-tolerant plant, showed considerable resistance. Among the *A. odora* plants grown at several different irradiances, the plants grown at the lowest irradiance, 10 µmol photons m⁻² s⁻¹, were most resistant to fluctuating-light-induced PSI photoinhibition. Their leaves showed the highest Chl content per leaf area (Chl_area_) and P700_area_ and the lowest Cyt *f*/Chl and Cyt *f*/P700. The electron flow rate per P700 was lowest in such leaves (Terashima et al. 2021). On the other hand, concerning PSII photoinhibition per se, it is established that high light-grown leaves are more resistant (Anderson and Aro 1993; Miyata et al. 2012, 2015; Öquist et al. 1987), although there are some exceptions (Öquist et al. 1992).

Kono et al. (2025) investigated PSI tolerance in the leaves from transgenic tobacco (*Nicotiana tabacum* L.) lines with varying Cyt *b*_6_*f* levels and found that a marked reduction in Cyt *b*_6_*f* levels conferred substantial PSI photoprotection against the FL-induced PSI damage, even under extremely high light pulses. In such leaves of low levels of Cyt *b*_6_*f*, however, the PSII quantum yield, Y(II), was lowest, indicating a trade-off between PSI photoprotection and overall photosynthetic productivity.

Taken together, suppression of the electron transport flow to PSI is effective in alleviating PSI photoinhibition. The decrease in the content of Cyt *b*_6_*f* per Chl with the decrease in the growth irradiance has been long known (Anderson 1986; Boardman 1977; Hikosaka and Terashima 1995; Schöttler and Tóth 2014). P700/Chl in these reports were rather stable in most of the experiments. In natural environment, the decrease in irradiance in most cases occurs due to vegetation cover. In such cases, the fraction of far-red light increases with decreasing irradiance. The effects of an increased fraction of far-red light on thylakoid composition have been examined in a few studies (Chow et al. 1990; Ibrahim et al. 2022; Tullberg et al. 2000). In these studies, however, P700/Chl decreased with far-red light enrichment. Since Cyt *f*/P700 increased in such plants, the electron flow to PSI would not be suppressed sufficiently. Alternatively, abundant far-red light in natural environment could still protect PSI. More ecophysiology-oriented studies are needed in this research field.

### 4.3 Flavodiiron proteins (FLVs)

Using electrons from Fd, FLV reduces O_2_ to water. Conifers use FLV to prevent photoinhibition at low temperatures in early-spring (Bag et al. 2023), whereas angiosperms have lost FLVs. For FLV-expressing rice plants at low temperature, see § 1.3. Yamamoto et al. (2016) expressed FLV in *A. thaliana* WT and *pgr5* mutant. FLV expression significantly increased rates of pseudo-CEF in the mutant plants, particularly at high light intensities, and partially restored the photosynthetic phenotype. In WT plants, FLV did not compete with PGR5-dependent CEF. Wada et al. (2018) expressed FLV in rice single mutant of PGR5, that of NDH, and a double mutant, and exposed these lines to the FL and found that PSI photoinhibition was not detected in any of these lines.

## 5. Roles of NDH and PGR5/PGRL1 in protection from PSI photoinhibition

As already detailed in § 1, NDH transfers electrons from reduced Fd to the PQ pool rather than passing them to O_2_ as FLVs do (Richardson et al. 2021; Takeuchi et al. 2025a, 2026). NDH also limits acceptor-side over-reduction. Although PGR5 contributes to proton motive force formation and photosynthetic control, direct evidence that PGR5 or PGRL1 prevents chilling-induced PSI photoinhibition is lacking. On the other hand, in the PSI photoinhibition studies using FL and rSPs, the roles of these two pathways have been compared.

### 5.1 Differences between the two CEF-PSI pathways

Although both NDH and PGR5 transfer electrons from Fd to the PQ pool, they play different roles. Table 1 summarizes their properties.

**Table 1.**
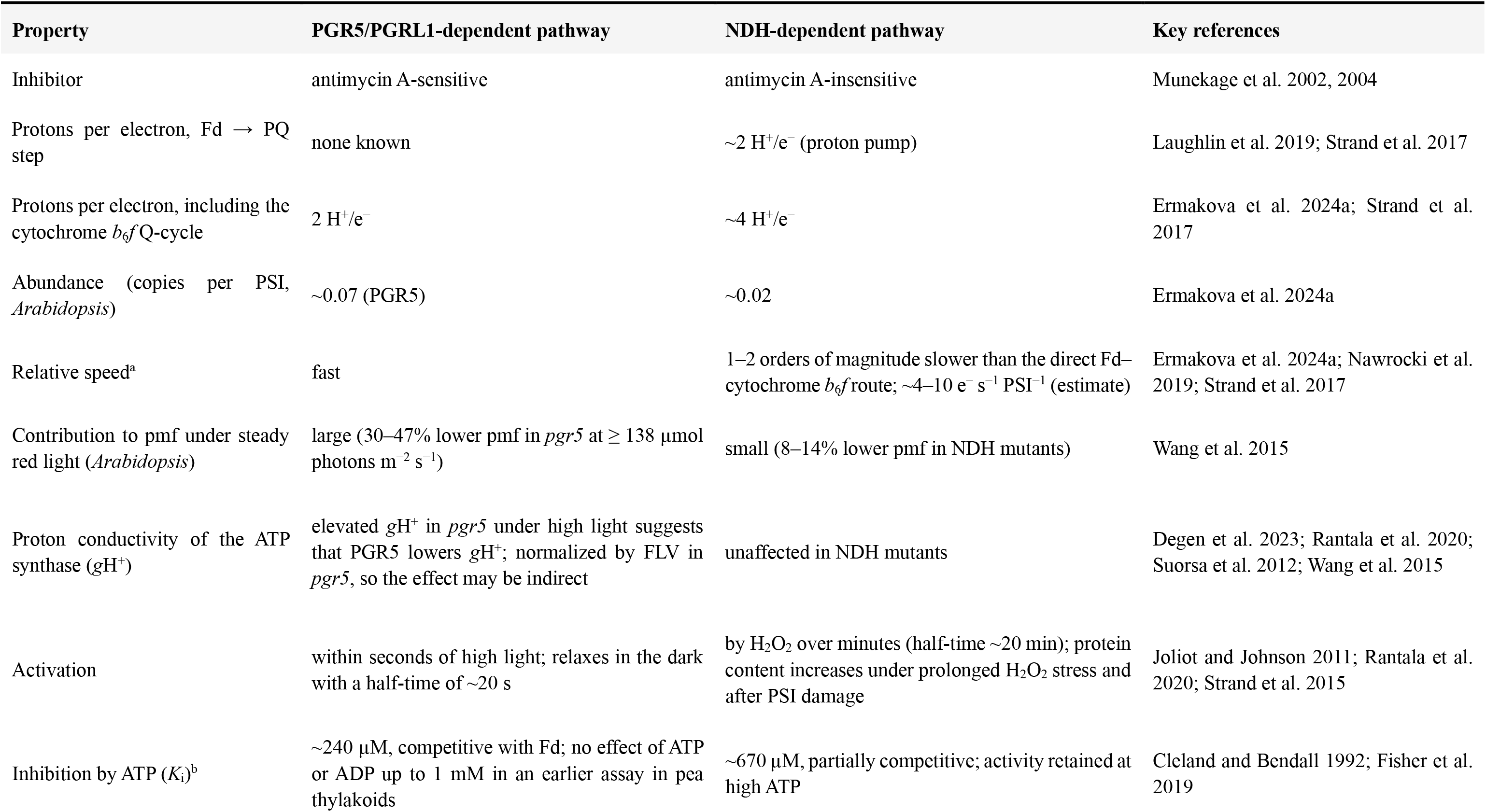
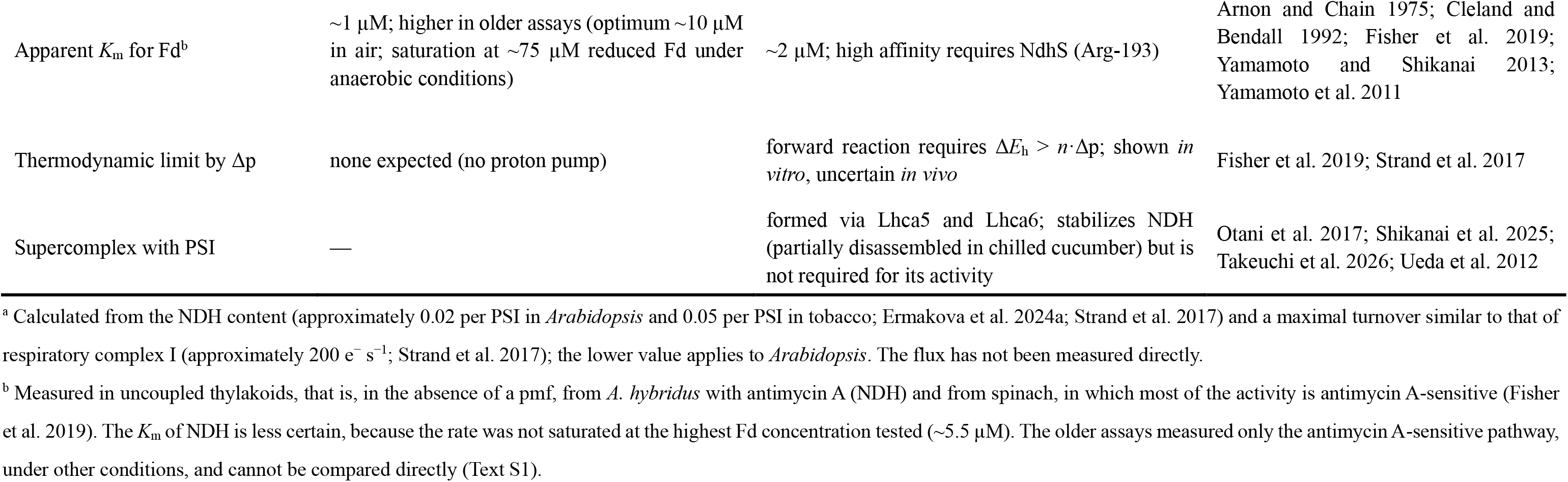
Properties of the two CEF-PSI pathways relevant to their division of roles.

#### Abundance and velocity

In *A. thaliana*, the NDH complex is present at 0.02 copies per PSI, compared with 0.07 for PGR5 (Ermakova et al. 2024a). Assuming a turnover similar to that of respiratory complex I (approximately 200 electrons s⁻¹; Strand et al. 2017), an NDH content of 0.05 per PSI, as reported for tobacco (Strand et al. 2017), would support a cyclic flux of approximately 10 electrons s⁻¹ per PSI, and the lower content of 0.02 per PSI in *A. thaliana* approximately 4 electrons s⁻¹ per PSI. The direct Fd–Cyt *b*₆*f* route, reported for *Chlamydomonas reinhardtii* P.A.Dang., is the fastest way to recycle electrons, and the NDH-dependent pathway is slower by one to two orders of magnitude (Ermakova et al. 2024a; Nawrocki et al. 2019). In *A. thaliana* in constant light, the mutants with reduced NDH activity showed only an 8–14% reduction in the light-induced pmf, whereas *pgr5* showed a 30–47% reduction at irradiances ≥ 138 µmol photons m⁻² s⁻¹ (Wang et al. 2015).

#### Proton stoichiometry

The chloroplast NDH complex is a homologue of respiratory complex I. Strand et al. (2017) provided evidence for its proton pumping: coupled thylakoids from *Spinacia oleracea* L. (spinach) synthesized ATP in the dark when Fd, NADPH and decyl-plastoquinone were supplied, and the ATP synthesis was abolished by uncouplers but insensitive to antimycin A. They estimated a stoichiometry of 2 H⁺ per electron transferred from Fd to PQ (2 H⁺/e⁻). When the Q-cycle at the Cyt *b*_6_*f* is included, each electron passing through the NDH pathway deposits 4 H⁺ in the lumen (4 H⁺/e⁻), whereas each electron passing through the PGR5 pathway deposits 2 H⁺ (2 H⁺/e⁻). In terms of ATP production, the NDH pathway would therefore produce twice as much ATP per electron (Ermakova et al. 2024a; Strand et al. 2017). PGR5/PGRL1 has no known proton-pumping module, and whether these proteins act as electron carriers is still debated (Ermakova et al. 2024a; see Table S2, row 8).

#### Regulation by ATP

Both pathways are inhibited by stromal ATP in a rapid and reversible manner (Fisher et al. 2019). The inhibition constants differ: 670 µM for NDH and 240 µM for the antimycin A-sensitive PGR5 pathway, both within the physiological range. Because the inhibition of NDH is only partially competitive, NDH retains some activity even when ATP is high, whereas the PGR5 pathway is switched off more completely. These measurements were made in uncoupled thylakoids from spinach, *Amaranthus hybridus* L. and *A. thaliana*. The inhibition by ATP is not, however, settled: in an earlier assay of the antimycin A-sensitive Fd–PQ reductase in washed pea thylakoids, ATP and ADP had no effect up to 1 mM, whereas NADPH was inhibitory (Cleland and Bendall 1992). The reason for this difference is not known. This weakens the ATP feedback hypothesis as the explanation for division of their roles (see Table S2, row 3).

#### Sensitivity to the proton motive force

Because NDH pumps protons, it must work against the proton gradient across the thylakoid membrane. The proton motive force, pmf or Δp (in mV), has an electrical component, the membrane potential ΔΨ, and a chemical component, the ΔpH across the thylakoid membrane:

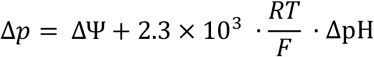

where *R*, *F* and *T* are the gas constant, the Faraday constant and the absolute temperature. At 25 °C (298 K), 2.3 × 10³ · (*RT*/*F*) equals 59 mV. We should also consider Δ*E*_h_, the difference in redox potential between the electron acceptor and the electron donor of NDH, that is, between PQ/PQH_2_ and Fd_ox_ /Fd_red_ (Δ*E*_h_ = *E*_h_ (PQ) − *E*_h_ (Fd)). The *E*_h_ of component A is expressed as

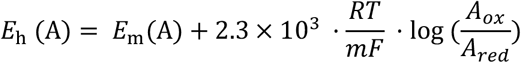

where *E*_m_(A) is the midpoint potential of A, at which [A_ox_] = [A_red_], and *m* is the number of electrons involved in the redox reaction of A. Δ*E*_h_ is, therefore, large when Fd is highly reduced and/or the PQ pool is oxidized, and small when Fd is oxidized and/or the PQ pool is reduced. From Δ*E*_h_ in mV, the energy available per mol of electrons is 10⁻³· Δ*E*_h_ · *F*. A pump that moves *n* protons per electron must spend energy corresponding to *n*·Δp against the gradient. We therefore call Δ*E*_h_ − *n*·Δp the net driving force of NDH. The forward reaction (Fd → PQ) can occur only when the net driving force is positive; when it is zero, the net rate is zero, and when it is negative, the reverse reaction is favored. Strand et al. (2017) used midpoint potentials of −430 mV for Fd and +80 mV for PQ at pH 7.5. With these values and treating PQ as a two-electron couple (*m* = 2), Δ*E*_h_ for 90% reduced Fd and PQ pools is ∼ 540 mV (520 mV in Strand et al. 2017), so NDH can operate as a 2 H⁺/e⁻ pump against a Δp of 180 mV. However, if Fd equilibrates with NADPH (midpoint −335 mV), or if the PQ pool becomes highly reduced while Δp stays high, the net driving force shrinks and the reaction approaches a standstill (Strand et al. 2017). In thylakoids from spinach and *A. hybridus*, collapsing the pmf with uncouplers doubled both the amplitude and the rate of the NDH-dependent post-illumination fluorescence rise, which is the expected behavior of a pump (Strand et al. 2017). The PGR5 pathway, which pumps no protons of its own, should not be limited in this way. Whether the back pressure of the pmf actually limits NDH *in vivo*, however, is still elusive (see § 9.1 and Table S2, row 1).

#### Thioredoxins (Trxs)

Trxs receive electrons from reduced Fd through Fd–Trx reductase (FTR) and regulate many chloroplast proteins (for a review, see Yoshida and Hisabori 2023). The m-type Trxs appear to restrain both CEF-PSI pathways. In *A. thaliana*, loss of Trx m4 increased NDH activity, overexpression of Trx m in tobacco suppressed it, and Trx m4 partially inhibited the PGR5-dependent pathway in vitro (Courteille et al. 2013). Okegawa and Motohashi (2020) further showed that Trx m4 forms a disulfide-linked complex with PGRL1 and suppresses the PGR5-dependent pathway; during photosynthetic induction, the complex temporarily dissociates, and CEF-PSI is activated. By contrast, the x- and y-type Trxs work together with CEF-PSI. In the *trx x* and *trx x trx y1 trx y2* mutants of *A. thaliana*, the PSI acceptor side was more reduced than in WT, particularly under FL, and PSI photoinhibition was more severe, although NDH activity was unchanged (Okegawa et al. 2023). In the *pgr5 trx x trx y1 trx y2* quadruple mutant, the acceptor side was even more reduced, and the leaves bleached and died under FL (Okegawa and Sakamoto 2026). Other Trxs also affect photosynthesis under FL: Trx m1 and m2 are needed for full photosynthetic activity in the HL phases, and NADPH-dependent Trx reductase C (NTRC) for the dynamic responses of NPQ and of the PQ redox state (Thormählen et al. 2017); the combined loss of Trx f1 and NTRC also makes PSI core proteins more vulnerable to light (Thormählen et al. 2015). Thus, Trxs can both restrain and support CEF-PSI, and how they regulate the two pathways under red and FR light deserves closer examination.

#### Interaction with Fd

Both pathways accept electrons from reduced Fd, so the way each binds Fd could, in principle, decide which pathway is used. For NDH, the binding site is known. The subunit NdhS has an SH3-like fold, and its conserved Arg-193 is required for the high-affinity, electrostatic binding of Fd: in ruptured *A. thaliana* chloroplasts, NDH-dependent PQ reduction was almost saturated at 3 µM Fd in WT, whereas in NdhS-deficient mutants it remained low even at 20 µM Fd (Yamamoto and Shikanai 2013; Yamamoto et al. 2011). For the PGR5-dependent pathway, the site is less clear. PGRL1 interacts with Fd, PGR5, PSI, the Cyt *b*_6_*f* complex and FNR (DalCorso et al. 2008), and recombinant PGRL1 reduces quinones with electrons from Fd in an antimycin A-sensitive manner (Hertle et al. 2013). Whether PGRL1 is the Fd–PQ reductase itself is still debated (Maekawa et al. 2024; see Table S2, row 8). Measured in the same system, the two pathways differ little in their affinity for Fd: the apparent *K*_m_ was approximately 2 µM for NDH, measured in *A. hybridus* with antimycin A, and 1 µM in spinach, in which most of the activity is antimycin A-sensitive (Fisher et al. 2019); the value for NDH is less certain, because the rate was not saturated at the highest Fd concentration tested (approximately 5.5 µM). Older assays of the antimycin A-sensitive pathway required more Fd, with an optimum of about 10 µM in air (Arnon and Chain 1975) and saturation at about 75 µM reduced Fd under anaerobic conditions (Cleland and Bendall 1992). These assays did not measure NDH and used other species and conditions, so their values cannot be compared directly with those of Fisher et al. (2019). All these values are similar to those of the other Fd-dependent enzymes of the stroma: the dissociation constant for Fd is about 1–10 µM for FNR at physiological ionic strength, about 1 µM for nitrite reductase and about 10 µM for Fd-dependent glutamate synthase (Knaff and Hirasawa 1991; see Text S1). The two pathways therefore compete with these enzymes for reduced Fd on roughly equal terms, and FNR, which is present at about half the amount of Fd (Böhme 1978; Terashima and Inoue 1985), is the largest competitor. Total Fd is unlikely to limit CEF-PSI, because chloroplasts contain roughly one to six Fd per PSI, or 3–6 mmol Fd per mol Chl (Antal et al. 2013; Böhme 1978; Terashima and Inoue 1985), which, with a chlorophyll concentration of 30–60 mM in the chloroplast (Antal et al. 2013), corresponds to about 0.1–0.4 mM (see Text S1), more than ten times these *K*_m_ values. What can limit CEF-PSI is the fraction of Fd that is reduced, and this is set by the balance between the supply of electrons from PSI and their use by FNR and the other enzymes. Tagawa et al. (1963) already observed in isolated chloroplasts that cyclic photophosphorylation began when NADP⁺ ran out, and in leaves, Fd-dependent CEF-PSI increases with the reduction level of Fd and appears when linear electron flow is saturated (Ohnishi et al. 2023; Satoh et al. 2025). Here, kinetics and thermodynamics should be kept apart: the rate of each pathway depends on the concentration of reduced Fd (Cleland and Bendall 1992), whereas Δ*E*_h_ in the condition Δ*E*_h_ > *n*·Δp depends on the ratio of reduced to oxidized Fd. Under pure FR, PSI receives few electrons from PSII: the oxidized PQ pool favors the forward reaction of NDH, but little Fd is reduced, which is why CEF-PSI under FR needs a small supply of electrons from PSII (Arnon and Chain 1975; Joët et al. 2002; see Table S2, row 2). Because both pathways draw on the same pool of reduced Fd with similar affinity, affinity alone cannot explain why one pathway is favored under one light quality and the other under another; the explanation must be sought in the pmf, ATP and the redox state of the chain (see §§ 5.2–5.4).

#### Physical organization

NDH forms a supercomplex with PSI through the minor antenna proteins Lhca5 and Lhca6; the stromal loop of Lhca6 is required for binding to NDH (Otani et al. 2017). The supercomplex stabilizes NDH. In chilling-sensitive cucumber, the PSI–NDH supercomplex is partially disassembled and NDH activity is lost within 1 h of chilling in the light, and this loss accompanies PSI photoinhibition (Takeuchi et al. 2026; see § 1.3). However, in the liverwort *M. polymorpha*, the supercomplex is not required for NDH activity: NDH functions without Lhca5 and Lhca6 (Ueda et al. 2012). Cyanobacterial NDH-1 has two Fd-binding site candidates (Laughlin et al. 2019), and the supercomplex may place NDH close to the Fd reduced by PSI; however, as noted above, both pathways draw on the same pool of reduced Fd, so physical proximity alone cannot explain the division of roles between light qualities.

#### Proton conductivity of the ATP synthase

It has been argued that PGR5 may lower the proton conductivity of the ATP synthase, *g*_H⁺_ (Rantala et al. 2020; Suorsa et al. 2012; Wang et al. 2015), whereas *g*_H⁺_ is unaffected in NDH mutants (Wang et al. 2015; see Table 1).

### 5.2 Red FL: How does the PGR5 pathway protect PSI?

In red FL, PSI is protected by a sequence of events (Kono and Terashima 2016). When the irradiance is increased from 30 to 1,200 µmol photons m⁻² s⁻¹, electrons gush into PSI at a rate faster than the rate of electron consumption by CBB cycle, photorespiration and other sinks. Thus, the acceptor side of PSI becomes reduced (high Y(NA)). CEF-PSI removes a part of the excess electrons and transfers them to the PQ pool. The protons deposited in the lumen by CEF-PSI build up ΔpH, which suppresses the oxidation of PQH_2_ at the Cyt *b*_6_*f* complex (photosynthetic control), and the flow of electrons into PSI is suppressed. As a result, the acceptor side is relieved and P700 is oxidized: Y(NA) decreases and Y(ND) increases. Kono and Terashima (2016) argued that the relief of the acceptor side must come first: because Y(I) + Y(ND) + Y(NA) = 1, Y(ND) can become large only when Y(NA) has become small. This depends on PGR5. When the PGR5 pathway was blocked, either genetically in *pgr5* or pharmacologically with antimycin A, Y(NA) rose step by step during the FL treatment, and the large Y(ND) of the HL phase never developed. P_m_ of *pgr5* and of antimycin A-treated WT leaves fell to 20% and 40% of the initial value, respectively, compared with 70% in untreated WT (Kono and Terashima 2016). The large Y(ND) in the HL phase was observed only when the PGR5 pathway was functional, either in WT or in *crr2-2*.

Two terms describe the restriction of electron flow at the Cyt *b*_6_*f* complex. "Photosynthetic control" is an old concept (Rumberg and Siggel 1969; Tikhonov et al. 1981; West and Wiskich 1968): when the lumen becomes acidic, protonation of the histidine ligand of the Rieske Fe–S protein, whose p*K*_a_ lies between about 6 and 6.5 in the oxidized state, slows the oxidation of PQH_2_ at the Q_o_ site (Tikhonov 2024; Vilyanen and Kozuleva 2025). "Reduction-induced suppression of electron flow" (RISE) describes the similar bottleneck from the PQ side: a highly reduced PQ pool limits the turnover of the Q-cycle even without a change in lumen pH (Miyake 2020; Shimakawa and Miyake 2018). Degen and Johnson (2024) treat the two as a pH input and a redox input on the same step. In leaves, the two inputs usually occur together, because the conditions that acidify the lumen also reduce the PQ pool. They differ, however, in what they require: the pH input requires a ΔpH, whereas the redox input requires a reduced PQ pool, a difference that becomes important under FR (see § 5.3). An increase in Y(ND) alone does not show that electron flow is restricted at the Cyt *b*_6_*f* complex by either input; the rate constant of P700⁺ re-reduction is a more direct indicator (Degen and Johnson 2024).

The *pgr5* phenotype also shows that the mutant fails to build and hold the pmf. In white FL, *pgr5* leaves generate a normal pmf during LL, but the pmf collapses almost completely at the transition to HL, whereas in WT, it increases sharply (Suorsa et al. 2012). The consequence is that P700 stays reduced throughout the HL phase in *pgr5* (Suorsa et al. 2012), and the acceptor side is also reduced. The fact that the damage originates on the acceptor side is shown by two observations: methyl viologen, an artificial electron acceptor, restores P700 oxidation in *pgr5* in high light (Munekage et al. 2002), and lowering the O₂ concentration to 2.7% or 0% during the red FL enhanced CEF-PSI in *pgr5*, released the acceptor-side limitation, and abolished PSI photoinhibition (Kono et al. 2014). In red FL, the protective effect of CEF-PSI comes from relieving the acceptor side and building the pmf, and the pathway that can do this fast enough is the PGR5 pathway.

Why is this role assigned to PGR5 instead of NDH? In the HL phase, a large pmf must be built quickly from a large electron flux. This exactly fits the characteristics of the PGR5 pathway (Table 1). It can also be understood why NDH is largely dispensable in red FL, at least in *A. thaliana*. NDH can recycle only a few percent of the maximum linear flux (Table 1), which is negligible in the HL phase. When the linear electron flow is fast, the PQ pool becomes strongly reduced and the pmf is high. Both changes diminish the net driving force of NDH (Δ*E*_h_ − *n*·Δp; Table 1) and push NDH toward a standstill.

However, this conclusion should not be generalized. Photosynthetic responses of the rice NDH mutant *crr6*, grown at 28 °C, to HL in FL (1,500/200 | 10 min/10 min) were almost identical to those of WT, whereas the mutant phenotype appeared in the LL phase, where NPQ was lower and the PQ pool was more reduced than in WT (Yamori et al. 2016). NDH plays a larger physiological role in rice (Yamori et al. 2011, 2016) than in *A. thaliana* or tobacco, whose NDH mutants grow normally and show little change in NPQ or electron transport, even at low irradiances (Munekage et al. 2004; Shikanai et al. 1998).

### 5.3 Far-red light: Why does the NDH pathway take over?

#### What does FR change?

Adding far-red light (FR) to the FL changed the state of PSI in both phases (Kono et al. 2017). In the LL phase, 10 s after the 100th HL pulse, Y(ND) rose from 0.006 without FR to 0.389 with FR, and Y(NA) fell from 0.432 to 0.224. In the HL phase, Y(ND) rose from 0.253 to 0.528 and Y(NA) fell from 0.428 to 0.232. FR thus both increased P700 oxidation and decreased acceptor-side limitation. The high Y(ND) of the LL phase cannot be explained by ΔpH-dependent photosynthetic control, because at the irradiance of 30 µmol photons m⁻² s⁻¹ the pmf is small, and in an FL regime with much longer phases the total pmf measured by the electrochromic shift was not clearly changed by FR (Kono et al. 2020). Instead, FR excites PSI selectively, so that P700 is oxidized faster than PSII can supply electrons to re-reduce it. This corresponds to the "linear mode" of Joliot and Johnson (2011): under FR, the electrons released by P700 are passed on to acceptors and few return, so that P700 is oxidized rapidly and the PQ pool stays largely oxidized. It follows that the redox input of photosynthetic control (see § 5.2) is not engaged in FR; only the pH input can act, but this is also weak in the LL phase. Consistent with this, when FR is given alone after darkness, the PGR5-dependent control of P700 remains in its inactive dark state (Rantala et al. 2020). In short, FR changes the main cause of P700 oxidation. In red FL, P700 is oxidized mainly by the pmf built by CEF-PSI; with FR, it is oxidized by the balance of excitation between the two photosystems. This is the basis of what may be called the masking hypothesis. In the sequence mentioned in § 5.2, the task of the PGR5 pathway was to relieve the acceptor side and build the pmf that oxidizes P700. Addition of FR reaches the same two endpoints, low Y(NA) and high Y(ND), without going through CEF-PSI. The specific contribution of PGR5, a fast, pmf-driven oxidation of P700, is therefore no longer required, and *pgr5* recovers to 92% of P_m_ when FR is present (Kono et al. 2017). This does not mean that FR acts "before" or "instead of" CEF-PSI. Every oxidation of P700 releases one electron to Fd, and these electrons must be recycled; P700 oxidation and CEF-PSI run at the same time and support each other. What FR removes is the need for the fast and large pmf-generating step. Whether P700⁺ is protective itself has been questioned (Rantala et al. 2020; Sonoike 2025); we regard high Y(ND) as protective only when it is accompanied by an oxidized acceptor side, a condition that FR meets (see Table S2, row 5).

#### Why NDH? A working hypothesis

If the pmf-generating step no longer must be fast, why does the protection by FR still require a CEF-PSI pathway, and why NDH in particular? Our working hypothesis, which may be called the redox–Δp gating, is that the redox state of the electron transport chain and the size of the pmf gate the two pathways differently. A proton pump such as NDH can run forward only while the redox driving force from Fd to PQ exceeds the back-pressure of the proton gradient, that is, while the net driving force (Δ*E*_h_ − *n*·Δp) is positive (Strand et al. 2017; see § 5.1). In FR, the PQ pool is oxidized, so Δ*E*_h_ is large, and the pmf is moderate, so *n*·Δp is small. Thus, the forward reaction of NDH is thermodynamically favored. Defining this hypothesis by the net driving force, rather than by Δp alone, makes it consistent with the observation that NDH-dependent CEF-PSI operates in the H_2_O_2_-accumulating line GO5 despite the pmf about twice that of WT, because in GO5 the stroma is strongly reduced and Δ*E*_h_ is correspondingly large (Strand et al. 2015). That NDH supplies electrons to the PQ pool under FR is indicated by leaves of *A. thaliana*: when actinic light was turned off with FR kept on, FR re-oxidized P700 more slowly in WT than in plants with impaired NDH activity, as in *ndhB*-disrupted tobacco (Shikanai et al. 1998, 2025). In tobacco leaves under FR, cyclic PSI activity is weak in air, because the intersystem chain is too oxidized; it becomes fast when the PQ and NADP pools are partly reduced under anaerobiosis, and the fast cycling requires NDH (Joët et al. 2002). This observation indicates two important points. First, in FR, NDH is the pathway that carries out electron cycling. Second, the rate of cycling depends on an adequate supply of reduced stromal donors, and not on the light. The redox–Δp gating determines whether the forward reaction of NDH is possible, but how fast it runs also depends on how much reduced Fd and NADPH is available. In our FR-supplemented FL (or rSPs), the red light of the LL phase continues to drive PSII (Kono et al. 2017). In an FL regime with much longer phases (800/30 | 10 min/15 min; Kono et al. 2020), adding FR to low red light oxidized the PQ pool while increasing the PSII yield. FR thus keeps the chain on the oxidized side without cutting off the supply of electrons from PSII, the condition NDH needs.

#### High proton efficiency at low flux

In the presence of FR, the role of CEF-PSI is not to drive a large electron flux but to provide a safe return route for the electrons released by P700 oxidation, together with just enough pmf to keep the Cyt *b*₆*f* complex under minimal photosynthetic control. NDH is suited to this role because its H⁺/e⁻ ratio, including the Q-cycle, is 4, whereas that of the PGR5 pathway is 2 (Table 1). Therefore, a small flux through NDH has a disproportionate effect on the pmf. A natural objection is that proton pumping activity should show up in NPQ. However, this is not observed in *A. thaliana* under steady red light: FR changed the PSI parameters below 500 µmol photons m⁻² s⁻¹, but left Y(II), Y(NO) and Y(NPQ) unchanged (Kono et al. 2017). The total proton input through NDH is small, which would be enough to support the small photosynthetic control needed for P700 oxidation but not enough to raise NPQ. In addition, FR increases the proton conductivity and proton flux through the ATP synthase (Kono et al. 2020), so that additional proton input can be used for ATP synthesis rather than stored as pmf. This applies to *A. thaliana* in steady light and to tobacco, in which the disruption of *ndhB* did not change NPQ at 20–2,000 µmol photons m⁻² s⁻¹ (Shikanai et al. 1998); in rice, whose NDH content is higher, *crr6* shows a clearly lower NPQ in low light (Yamori et al. 2011, 2016). The same reasoning explains the asymmetry seen with FR: *pgr5* still has the pathway that the FR regime needs and recovers to the WT level, whereas P_m_ in *crr2-2* and *crr4-3*, which lack it, remained at 80% of the initial level even with FR (Kono et al. 2017).

#### The operating window of NDH

The FL (or rSPs) regime of Kono et al. (2017) spends 92.6% of its time in the LL phase (1,200/30 | 800 ms/10 s). Low, steady electron cycling is the operating domain of NDH; Joliot and Johnson (2011) noted that, although the NDH content is far too low to sustain the cyclic rates required under strong light, NDH can carry a significant fraction of CEF-PSI under low light. Because NDH pumps protons, its own activity raises the pmf, until *n*·Δp approaches Δ*E*_h_ and the forward reaction stops (Table 1). The upper limit of NDH operation is therefore set by the pmf. In the LL phase of FR-supplemented FL, the pmf is moderate and stays below this limit, whereas in the HL phase of red FL it is high enough to stop NDH. The lower limit is set not by the pmf but by the supply of electrons: the PQ pool must be oxidized enough to keep Δ*E*_h_ large, but enough reduced Fd and NADPH must remain to supply the reaction, because cycling almost stops when the electron transport chain is too oxidized (Joët et al. 2002). Similarly, in isolated chloroplasts, cyclic photophosphorylation under 715-nm light required a small supply of electrons from PSII and stopped when PSII was blocked by DCMU (Arnon and Chain 1975). In addition, in the LL phase the stromal ATP level falls while the demand for ATP persists, and a moderate fall in ATP activates NDH first (Fisher et al. 2019); ATP feedback would thus favor NDH under the same conditions, although the ATP dynamics under FR have not been measured.

#### Maintaining redox poise across both phases

FR added to the FL protected PSI fully only when it was present in both phases of the FL; FR in the HL phase alone or in the LL phase alone gave only partial protection (Kono et al. 2017). This can be explained as follows. In the LL phase, FR keeps P700 and the donor-side carriers in an oxidized state. These carriers (plastocyanin, the Cyt *b*_6_*f* complex and the PQ pool) together hold a few tens of electron equivalents per P700. This oxidized pool is small, but it prevents P700 from becoming fully reduced at the onset of the next HL pulse. During the HL pulse, FR keeps P700 oxidized while CEF-PSI returns the released electrons. Therefore, the protective state is not a stock of P700⁺ built in advance, but a continuously maintained redox poise of the whole chain. If ΔpH, photosynthetic control and P700 oxidation are relaxed in the LL phase, the chain must rebuild them in the next HL pulse, and the acceptor side is transiently over-reduced during this "cold start". FR prevents this cold start. This also resolves an apparent paradox in the NDH literature: the loss of NDH makes the PQ pool more reduced under low light (Ueda et al. 2012; Yamori et al. 2011, 2016), although NDH reduces PQ. If the pmf and ATP generated by NDH sustain the CBB cycle and minimal photosynthetic control when the light is low, then NDH acts as an enzyme that keeps the intersystem carriers poised, not merely as a PQ reductase (Yamori et al. 2011). Under FR, this poising function is what the plant needs, and the NDH pathway provides it.

### 5.4 Summary and outlook: light quality sets the redox poise and the pmf, and these gate the two pathways

#### Two regimes, one rule

Table 2 places the two regimes side by side. In the high-light phase of red FL, a reduced PQ pool and a high pmf drive the net driving force of NDH (Δ*E*_h_ − *n*·Δp) close to zero, and P700 is oxidized by the pmf that the PGR5-dependent pathway builds within seconds (Fig. 4a, c). Under FR-supplemented FL, an oxidized PQ pool and a moderate pmf make the net driving force large, and P700 is oxidized by the excitation balance, so that NDH can run forward with high proton efficiency (Fig. 4b, c). One rule covers both regimes: light quality sets the redox state of the PQ pool and the pmf, and these two variables decide which pathway can run. It explains the result of Kono and Terashima (2016), its reversal in Kono et al. (2017), and why the reversal stayed hidden until FR was added: red LED light produces the one regime in which NDH has the least to contribute.

**Fig. 4.**
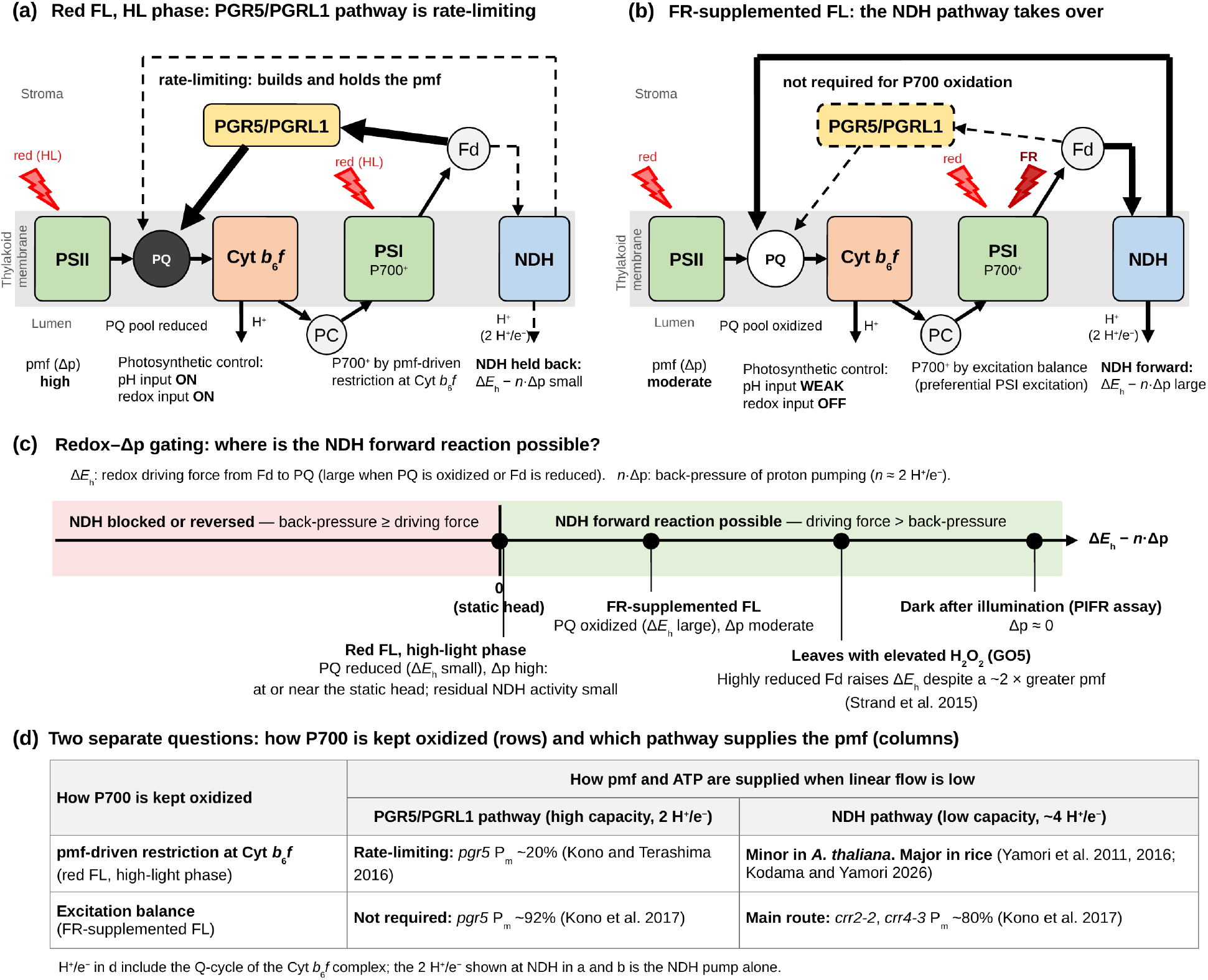
Light quality sets the redox poise and the pmf, and these gate the two CEF-PSI pathways. (**a**, **b**) Schemes of the electron transport chain in the high-light phase of red FL (**a**) and under FR-supplemented FL (**b**); only the light quality differs between the two panels. Filled PQ, reduced pool; open PQ, oxidized pool; arrow thickness indicates electron flux; dashed arrows and outlines indicate a pathway that is held back or not required. The 2 H⁺/e⁻ shown at NDH is the stoichiometry of the NDH pump alone. (**c**) Position of the two regimes on the axis Δ*E*ₕ − *n*·Δp that governs the NDH forward reaction (redox–Δp gating hypothesis). The high-light phase of red FL is placed at the static head (Δ*E*ₕ − *n*·Δp ≈ 0) rather than far below it, because the loss of NDH still lowers the light-induced pmf by ∼10% under steady red light (Wang et al. 2015). Two reference states are shown for comparison: leaves with elevated H₂O₂ (the GO5 line), in which NDH-dependent CEF-PSI operates despite a pmf ∼2 times that of the wild type, because the stroma is strongly reduced (Strand et al. 2015); and the dark period after illumination, in which the post-illumination fluorescence rise (PIFR) is measured at Δp ≈ 0. The axis is schematic; positions indicate only the sign of the term and relative distances. (**d**) Two-axis summary: the means by which P700 is kept oxidized (rows) and the pathway that supplies the pmf and ATP when linear flow is low (columns); the H⁺/e⁻ values include the Q-cycle of the cytochrome *b*₆*f* complex. Data from Kodama and Yamori (2026), Kono and Terashima (2016), Kono et al. (2017, 2025), Theeuwen et al. (2022), Wang et al. (2015) and Yamori et al. (2011, 2016). CEF-PSI, cyclic electron flow around PSI; Cyt, cytochrome; Fd, ferredoxin; FL, fluctuating light; FR, far-red light; NDH, NADH dehydrogenase-like complex; PC, plastocyanin; pmf, proton motive force; PQ, plastoquinone.

**Table 2.** Two light regimes and the resulting division of roles between the CEF-PSI pathways.

|  | **Red FL (high-light phase unless stated)** | **FR-supplemented FL (both phases)** |
|---|---|---|
| Main cause of P700 oxidation | pmf-driven restriction at cytochrome *b*<sub>6</sub>*f* (photosynthetic control) | excitation balance (selective PSI excitation) |
| PQ pool | reduced | oxidized |
| Inputs of photosynthetic control | pH input and redox input both engaged | pH input only, weak in the low-light phase |
| pmf | high in the high-light phase; collapses in *pgr5* | moderate; total pmf not clearly changed by FR |
| Δ*E*<sub>h</sub> − *n*·Δp for NDH<sup>a</sup> | near zero; NDH held back | large; NDH forward reaction favored |
| PGR5/PGRL1 pathway | rate-limiting: builds and holds the pmf | not required for P700 oxidation |
| NDH pathway | minor in *Arabidopsis*; larger in rice | main route for recycling electrons at low flux, with high proton efficiency |
| Y(ND) / Y(NA), wild type | low-light phase: 0.006 / 0.432<br>high-light phase: 0.253 / 0.428 | low-light phase: 0.389 / 0.224<br>high-light phase: 0.528 / 0.232 |
| Decisive observation | *pgr5* P<sub>m</sub> ~20%; *crr2-2* ~10% worse than the wild type | WT P<sub>m</sub> ~97%; *pgr5* ~92%; *crr2-2*, *crr4-3* ~80% |
| Main data sources | Kono and Terashima 2016; Kono et al. 2017 | Kono et al. 2017, 2020 |

Y(ND) and Y(NA) are values for wild-type *Arabidopsis* 10 s after the 100th high-light pulse (low-light phase) and during the pulse (high-light phase) in the FL regime of Kono et al. (2017), without FR (left) and with FR (right). P<sub>m</sub> values are those after the FL treatment, expressed relative to the value before the treatment. Other entries summarize §§ 5.2 and 5.3.
<sup>a</sup> Inferred from the redox–Δp gating hypothesis; not measured *in vivo* (§ 5.4; Table S2, row 1).

#### Two separate questions

Two questions have often been conflated in the literature (Fig. 4d). The first is by what means P700 is kept oxidized, and three means are now documented. One is the pmf-driven restriction at the Cyt *b*_6_*f* complex, built mainly by the PGR5-dependent pathway under red high light (see § 5.2). Another is a restriction of the electron flux into PSI from upstream, either by a low content of the Cyt *b*_6_*f* complex (Kono et al. 2025) or by PSII photoinhibition, which can itself protect PSI (Sonoike 2025): in *A. thaliana* leaves whose F_v_/F_m_ had been lowered by prior chilling, the residual PSI after rSPs increased as F_v_/F_m_ decreased, independently of ΔpH (Takeuchi et al. 2025b). Both protect PSI at the cost of photosynthetic capacity. The third is the excitation balance imposed by FR, which oxidizes P700 without a high pmf and, in the low-light phase of FL, raises, instead of lowers, CO_2_ assimilation (Kono et al. 2020). The second question is how the pmf and ATP are supplied when linear flow is low. Here, the two pathways share the task according to their capacity and efficiency: a large flux at two protons per electron through the PGR5-dependent pathway, or a small flux at four through NDH (Table 1). Seen this way, FR and a low Cyt *b*_6_*f* content act at the same point: both remove the need for a fast, pmf-driven oxidation of P700 and leave the plant with the second question, which the NDH pathway answers efficiently at low flux.

#### A general rule for low light

If the redox–Δp gating is correct, NDH should matter whenever the light is low enough to keep the PQ pool oxidized and the pmf moderate, whether or not FR is present. This is what the NDH mutants show: the Burren *ndhG* variant of *Arabidopsis* affects PSII efficiency only below approximately 350 µmol photons m⁻² s⁻¹ (Theeuwen et al. 2022, preprint), the loss of NDH in *M. polymorpha* increases PQ reduction only < 100 µmol photons m⁻² s⁻¹ (Ueda et al. 2012), and the rice *crr6* phenotype appears below the growth irradiance and in the low-light phase of FL (Yamori et al. 2011, 2016). In each case—paradoxically, since NDH reduces PQ—the intersystem chain is more reduced without NDH when the light is low, a paradox that the poising function of NDH resolves (§ 5.3). FR is the extreme case of this condition, a light that excites PSI while driving little electron flow from PSII. Seen in this way, the "low light versus high light" rule in § 2 and the "red versus far-red" reversal in § 5.3 are two expressions of the same gating. The rule is therefore about redox poise and pmf, not about irradiance as such, and C₄ plants illustrate the difference. In the bundle-sheath cells of NADP-malic enzyme species, which operate in full sunlight, the chloroplasts contain little PSII, the stroma receives NADPH from malate decarboxylation, and the CBB cycle needs ATP that linear flow cannot supply; it is in these cells that NDH accumulates to its highest levels (Takabayashi et al. 2005). What they share with an FR-illuminated leaf is not low light but a chain fed by little PSII and a reduced stroma, which should keep Δ*E*_h_ large enough for NDH to run even against a substantial pmf, as in GO5 (see Table S2, row 1), together with a demand for ATP that only a proton-efficient cycle can meet. The high NDH content serves the CO₂-concentrating mechanism rather than PSI protection (Takabayashi et al. 2005), but the condition that selects NDH is the same as under FR.

## 6. ROS: their generation and scavenging

### 6.1 Routes of ROS generation

Electrons are transferred within PSI from the P700 to Fd via A_0_, phylloquinone A_1_, and the Fe–S clusters F_X_, F_A_, and F_B_. Because these electron carriers have highly negative midpoint potentials (A_0A_/A_0B_, approximately −1200 mV; PhQ A_1A_/A_1B_, −800 to −600 mV; F_X_/F_A_/F_B_, −600 to −500 mV), their reduced forms can transfer electrons to O_2_ to generate the superoxide radical anion (O_2_^•^⁻). For comparison, the reduction potential of the O_2_/O_2_^•^⁻ couple is approximately −160 mV in water, while −600 to −500 mV in N,N-dimethylformamide (DMF) (Cherepanov et al. 2017; Degen and Johnson 2024; Kozuleva and Ivanov 2010; Ptushenko et al. 2008; Wardman 1990). O_2_^•^⁻ is converted to H_2_O_2_ through non-enzymatic dismutation or through dismutation catalyzed by chloroplast superoxide dismutase (SOD) localized near PSI on the stromal surface of the thylakoid membrane (Asada 2006; Ogawa et al. 1995). H_2_O_2_ can generate •OH through Fenton reactions involving the reduced PSI Fe–S clusters or free Fe^2+^, further amplifying PSI damage (Šnyrychová et al. 2006; Sonoike 2011). In addition, O_2_^•^⁻ is known to react directly with solvent-exposed [4Fe–4S] clusters, inducing their breakdown through cluster oxidation and release of Fe (Bulteau et al. 2003; Flint et al. 1993; Gardner 1997; Petronek et al. 2021). The Fe–S clusters of PSI may therefore be direct targets of O_2_^•^⁻.

Severe over-reduction of the PSI acceptor side provides an additional pathway for ROS production. Charge recombination of the P700⁺/A_0_⁻ or P700⁺/A_1_⁻ radical pair can form triplet P700* (^3^P700*), from which energy transfer to O_2_ produces singlet oxygen (^1^O_2_) (Kim et al. 2001; Rajagopal et al. 2005; Rutherford et al. 2012; Shimakawa et al. 2024; Shuvalov et al. 1986). Not all charge recombination leads to ^3^P700* formation, however; an alternative pathway returns the charge-separated state directly to the ground state without forming ^3^P700* (see § 6.3). ROS can therefore be generated in PSI by at least two pathways: O_2_^•^⁻ generation through electron transfer to O and ^1^O generation through charge recombination (Sonoike 2025). O_2_^•^⁻ is less reactive than •OH and ^1^O and therefore does not rapidly attack most proteins or lipids. By contrast, ^1^O_2_ and •OH are highly reactive but short-lived, so their effects are largely confined to pigments and proteins close to their sites of formation.

### 6.2 Sites of O_2_^•^⁻ generation within PSI

Which PSI electron carrier reduces O_2_ during chilling-induced PSI photoinhibition has long been debated. Chilling-induced PSI photoinhibition was barely detectable in darkness or under anoxic conditions, directly implying involvement of ROS in the damage. The early loss of Fe–S cluster activity in cucumber chilled in the light initially pointed to these clusters as the principal sites of both ROS production and damage (Shimakawa et al. 2024; Sonoike et al. 1995). However, the site where damage is first detected is not necessarily the site where ROS are generated. O_2_^•^⁻ reacts weakly with most biomolecules but selectively attacks Fe–S clusters (Flint et al. 1993; Furutani et al. 2023). O_2_^•^⁻ generated closer to the PSI core, for example at A_1_, could therefore damage the Fe–S clusters even if the clusters themselves are not its site of production. Because A_0_ and A_1_ have more negative redox potentials than F_A_/F_B_, electron transfer from their reduced forms to O_2_ is thermodynamically more favorable. Under normal electron transport conditions, however, the reduced states of A_0_ and A_1_ are short-lived, making electron accumulation at these cofactors unlikely. By contrast, the reduced states of F_A_/F_B_ are considerably longer-lived. The site of O_2_ reduction within PSI is therefore determined not only by the redox potential of each carrier but also by the lifetime of its reduced state and its accessibility to O_2_ (Khorobrykh et al. 2020).

Although A_1_ lies within the PSI complex, structural analyses have revealed internal protein cavities near A_1A_ and A_1B_ that connect to the external aqueous phase, providing possible routes for O_2_ to reach these sites (Cherepanov et al. 2017). Access to A_1_ alone, however, does not determine the dominant site of O_2_ reduction, because the relative contributions of the PSI cofactors vary with light intensity. Kozuleva and colleagues isolated PSI from *Chlamydomonas* grown under approximately 70 µmol photons m⁻^2^ s⁻^1^ and measured O_2_ photoreduction under 50–2,200 µmol photons m⁻^2^ s⁻^1^. At irradiances of 300 µmol photons m⁻^2^ s⁻^1^ or higher, phylloquinone A_1_ was the major site of O_2_ reduction, whereas F_A_/F_B_ made a greater contribution at lower irradiances (Kozuleva 2022; Kozuleva and Ivanov 2023; Kozuleva et al. 2021). Because the reduced states of F_A_/F_B_ persist longer than those of A_1_ and F_X_, electron transfer from F_A_/F_B_ to O_2_ is favored at lower irradiances. As irradiance increases, repeated excitation supplies electrons to the PSI acceptor chain faster than they can be removed from the terminal clusters, allowing reduced F_X_ and eventually reduced A_1_ to accumulate. A_1_ may therefore make a greater contribution to O_2_ reduction at higher irradiances. In intact leaves, however, photosynthetic control normally restricts electron flow into PSI, limiting the sustained accumulation of reduced F_X_ and A_1_. O_2_ reduction at A_1_ is therefore expected to become appreciable only when photosynthetic control is impaired or overwhelmed and the PSI acceptor side becomes severely over-reduced (Miyake 2020).

Fd is also the site of O_2_^•^⁻ formation (Badger 1985). Although Fd could become a victim of O_2_^•^⁻, O_2_^•^⁻ formed at Fd in the stroma would not attack PSI Fe–S centers. Similarly, addition of methyl viologen (MV), which is an electron acceptor from PSI and produces O_2_^•^⁻ through autoxidation upon reduction, suppressed PSI photoinhibition induced by constant light in isolated thylakoid membranes (Sonoike 1996a) or by rSPs in isolated chloroplasts (Takagi et al. 2016).

### 6.3 ^1^O_2_ generation through charge recombination

Among the possible fates of electrons accumulated on the PSI acceptor side are electron transfer to O_2_ and charge recombination with P700⁺. The latter can return electrons safely to P700⁺, but some recombination pathways generate ^3^P700* and thereby promote the formation of ^1^O_2_, potentially damaging PSI. Whether charge recombination protects or damages PSI may depend on the pathway it follows. Formation of ^3^P700* from P700⁺A_1_⁻ is thought to involve re-formation of the primary P700⁺A_0_⁻ radical pair, followed by population of the triplet state of P700*. Alternatively, the P700⁺A_1_⁻ can return to the ground state through a recombination pathway without forming ^3^P700*. Rutherford et al. (2012) proposed that these contrasting outcomes reflect the different energetics of A_1A_ and A_1B_. In their model, a back-reaction from F_X_⁻ or another downstream acceptor may avoid ^3^P700* formation by proceeding through A_1A_, whereas recombination through the B-branch may be more likely to re-form the primary P700⁺A_0B_⁻ radical pair and produce ^3^P700* (Bindra et al. 2024; Cherepanov et al. 2017; Degen and Johnson 2024; Ishikita and Knapp 2003; Rutherford et al. 2012).

Evidence for a protective role of charge recombination comes from high-light experiments with intact leaves. Tiwari and colleagues showed that continuous high-light exposure caused 10–35% photodamage to F_A_/F_B_ in wild-type *A. thaliana*, while having little effect on F_X_ function or P700 oxidation capacity (Tiwari et al. 2024). They proposed that, following partial inactivation of F_A_/F_B_, electrons accumulating at F_X_ can recombine safely with P700⁺ if photosynthetic control maintains a sufficient population of P700⁺. In the *pgr5* mutant, by contrast, impaired photosynthetic control limited the accumulation of P700⁺. Reduction then spread to F_X_ and was accompanied by damage to this cluster, resulting in severe PSI damage (Degen et al. 2026; Tiwari et al. 2024). Thus, charge recombination between F_X_⁻ and P700⁺ may limit the spread of damage from F_A_/F_B_ to F_X_ and the PSI core.

A different pattern of photo-oxidative damage to PSI is observed in rSPs. Shimakawa et al. (2024) found that such pulses damaged the PSI core near P700 more severely than the Fe–S clusters. Under these conditions, P700⁺ and reduced A_0_/A_1_ may coexist more frequently, increasing the likelihood of charge recombination that produces ^3^P700* and, consequently, ^1^O_2_. The short lifetime and diffusion distance of ^1^O_2_ could then account for the preferential damage near the PSI reaction center rather than the Fe–S clusters. It remains unclear, however, why recombination pathways that form ^3^P700* would be favored over pathways that return the charge-separated state safely to the ground state during rSPs, or to what extent the B-branch is directly involved in this process.

A major defense against triplet-mediated ^1^O_2_ formation in photosynthetic organisms is triplet–triplet energy transfer from triplet chlorophyll to carotenoids, which quenches the chlorophyll triplet before it can sensitize O_2_ (Cazzaniga et al. 2012; Qin et al. 2025; Shi et al. 2026). A comparable process has been demonstrated in the reaction-center complex of green sulfur bacteria, where triplet energy transfer from core-antenna BChl *a* to carotenoids produces triplet carotenoids (^3^Car) (Azai et al. 2020; Inagaki et al. 2026). Direct experimental evidence for such quenching of ^3^P700* by a carotenoid in land-plant PSI is, however, still lacking.

### 6.4 Summary of ROS generation in PSI under different environmental conditions

The mechanisms of ROS generation in PSI are summarized in Fig. 5. Under continuous high light, photosynthetic control can maintain P700⁺. Thus, even when a fraction of the F_A_/F_B_ clusters is damaged, electrons accumulated at F_X_ may be safely removed by charge recombination with P700⁺ through the A-branch. Although this reaction does not restore forward electron transfer to Fd, it may prevent damage from spreading to F_X_ and the PSI core (Tiwari et al. 2024). When photosynthetic control does not function adequately, as in *pgr5*, P700⁺ cannot be maintained, and over-reduction and damage spread to F_X_ (Fig. 5a), leading to severe PSI photoinhibition (Mazor et al. 2017; Tiwari et al. 2016, 2024).

**Fig. 5.**
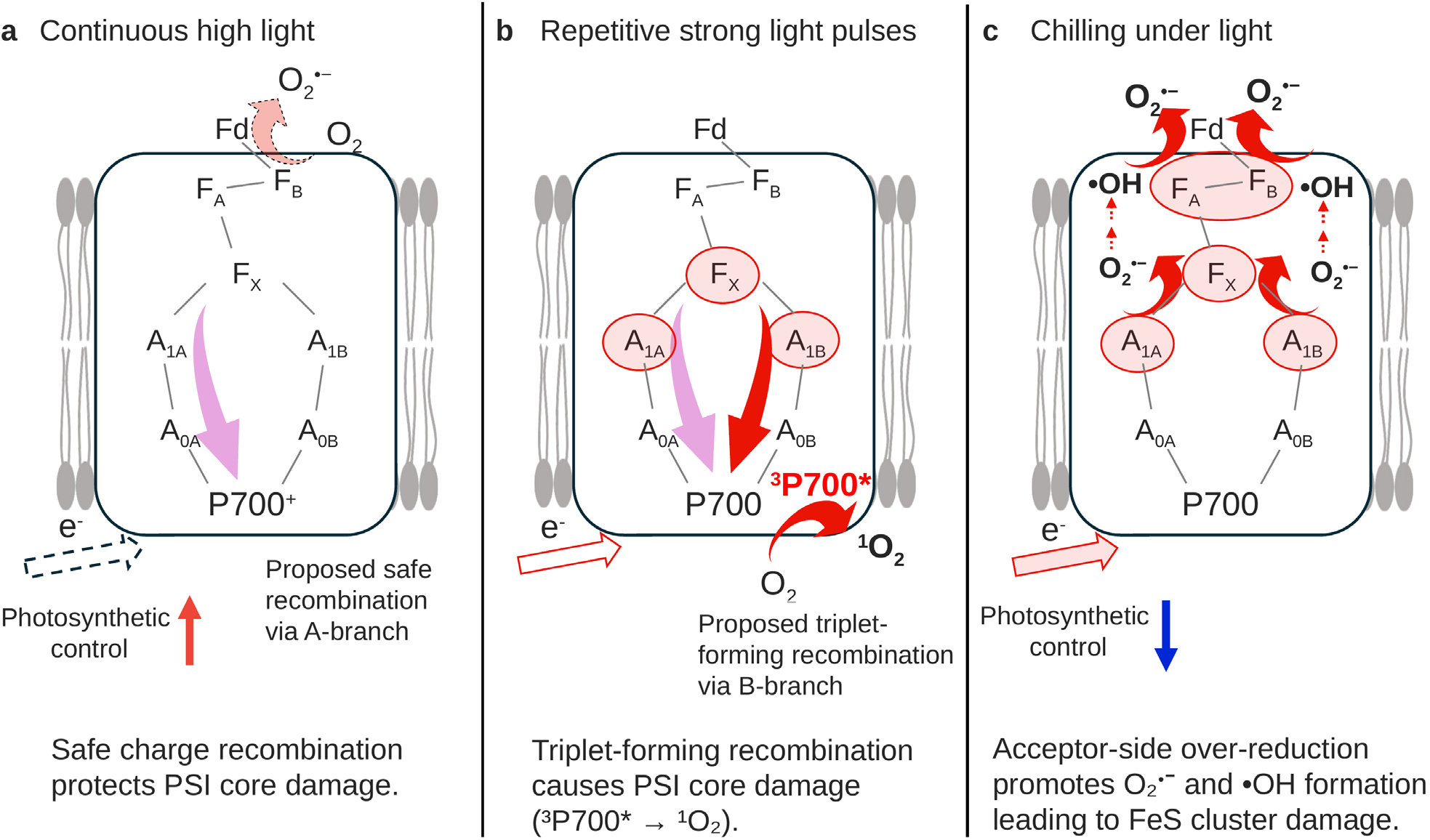
Proposed mechanisms of ROS production in PSI under different environmental conditions. (**a**) Under continuous high light, partial damage to F_A_/F_B_ may occur without leading to irreversible PSI core damage. Photosynthetic control maintains P700 in its oxidized state (P700⁺), allowing excess electrons to be safely removed through charge recombination from F_X_ to P700⁺ via the A-branch, even when F_A_/F_B_ is partially damaged. (**b**) Under repetitive saturating pulses, P700⁺ and reduced A_0_/A_1_ are more likely to coexist. Consequently, charge recombination via the B-branch may generate ^3^P700* more frequently and thereby increase ^1^O_2_ formation, potentially accounting for preferential damage near the PSI reaction center. (**c**) In cucumber, chilling-induced PSI photoinhibition progresses even under relatively weak light. At low temperature, the decline in NDH activity, together with uncoupling of the chloroplast H⁺-ATP synthase, impairs ΔpH formation and photosynthetic control. Consequently, over-reduction may extend beyond F_A_/F_B_ to F_X_ and A_1_. Because P700⁺ accumulation is also suppressed, safe removal of excess electrons through charge recombination from F_X_^−^ to P700⁺ becomes less effective. Electrons may therefore leak to O_2_ from multiple acceptors, including A_1_ and F_A_/F_B_, promoting O_2_^•^⁻ production. The resulting O_2_^•^⁻ may directly damage the Fe–S clusters or undergo nonenzymatic or SOD-catalyzed dismutation to form H_2_O_2_. The H_2_O_2_ thus produced can generate •OH through Fenton reactions involving the Fe–S clusters or free Fe^2+^, leading to further oxidative damage.

Under rSPs, and possibly, though to a much lesser extent, under FL, P700⁺ and reduced A_0_/A_1_ coexist more frequently. Consequently, charge recombination through the B-branch may more readily generate ^3^P700* and thereby increase ^1^O_2_ formation, potentially accounting for preferential damage to the region near the PSI reaction center. By contrast, the reduced states of the Fe–S clusters are thought to be relatively short-lived under rSPs, which would limit O_2_^•^⁻ production from these cofactors (Fig. 5b).

Finally, chilling-induced PSI photoinhibition in cucumber progresses even under relatively weak light of approximately 100–250 µmol photons m⁻² s⁻¹ (Fig. 5c). At low temperatures, NDH activity declines, while weakened association of CF_1_ with the thylakoid membrane impairs ΔpH formation, ATP synthesis, and photosynthetic control. Reduced ATP production, together with the temperature-dependent decreases in enzymatic reactions, strongly suppresses the CBB cycle and photorespiration, further decreasing electron consumption downstream of PSI. Consequently, even under weak light, electron flow into PSI may exceed the electron-consuming capacity of downstream reactions, allowing the reduced state to extend beyond F_A_/F_B_ to F_X_ and A_1_. Impaired photosynthetic control also suppresses the accumulation of P700⁺, making safe charge recombination from F_X_^−^ to P700⁺ less likely. Under chilling conditions, electrons may therefore leak to O_2_ from multiple acceptors, including A_1_ and F_A_/F_B_, promoting O_2_^•^⁻ production. The resulting O_2_^•^⁻ can damage the Fe–S clusters. In addition, dismutation of O_2_^•^⁻ produces H_2_O_2_, which can generate •OH through Fenton reactions involving the Fe–S clusters or free Fe^2+^, causing further oxidative damage (Hegedűs et al. 2008; Nakano et al. 2006; Šnyrychová et al. 2006). Consistent with a role of Fe in light-induced oxidative damage, cucumber grown under Fe-rich conditions is more susceptible to oxidative injury under light-chilling stress (Takeuchi et al. 2024).

Thus, the ROS species generated within PSI and the sites of production are likely to differ among stress conditions, depending on which acceptors become reduced, how long they remain reduced, and whether P700⁺ remains available for charge recombination.

Chilling injury caused by ROS generation involves the chain reaction of lipid peroxyl radicals that eventually causes membrane disruption and cell death. In chilling-tolerant plants, levels of antioxidants including ascorbic acid and tocopherols are high. In particular, α-tocopherol, a lipophilic antioxidant localized in the membrane, acts as scavenger of singlet oxygen and lipid peroxyl radicals. For reviews, see Asada (2006) and Mesa and Munné-Bosch (2023).

## 7. Roles of far-red light and allocation of excitation between PSI and PSII

To keep P700 in its oxidized form, excitation of PSI is prerequisite. In natural light environments in the terrestrial ecosystems, far-red light, which preferentially excites PSI, is abundant. When spillover of excitation from PSII to PSI occurs, PSI is further excited. Under fluctuating-light conditions, PSI may be most vulnerable when a leaf is suddenly exposed to high light after an extended period of low light. NPQ in such leaves is low and does not increase spontaneously. In contrast, the spillover of the excitation from PSII to PSI occurs almost instantaneously.

Concerning spillover, especially in shade-type chloroplasts, the occurrence of spillover itself is often doubted, because PSII is in the appressed membranes, whereas PSI is mostly in the non-appressed membranes (Andersson and Anderson 1980), and thus PSII and PSI are too far apart. A study of *Pinus sylvestris* L. (Scots pine) using time-resolved fluorescence analysis showed that the spillover increased in early spring with the decrease in the number of thylakoids per granum in the overwintered needles (Bag et al. 2020). These results support the notion that the spillover from PSII to PSI would not occur in chloroplasts with highly stacked grana.

We recently examined the degree of spillover in the leaf discs at 77 K using various plants, grown at high and low light. We measured the fluorescence induction at 760 nm following the model by Kitajima and Butler (1975). They assumed that there is no induction in intrinsic PSI fluorescence and that the fluorescence at 760 nm contains no PSII fluorescence. Then, the induction observed at 760 nm would be largely attributed to spillover from PSII. Although we confirmed that the intrinsic PSI fluorescence excited at 700 nm did not show any induction, the second assumption is not valid. Even at 760 nm, there was some vibronic PSII fluorescence. We estimated the leftover PSII fluorescence at 760 nm to be around 15% of the maximum fluorescence, *F_m_*, at 760 nm (Terashima et al. 2025). After illumination with weak far-red light and subsequent dark treatment for less than 2 min, the leaf discs in state 1 were frozen and their fluorescence induction was analyzed at 690 and 760 nm. Taking account of the PSII contamination, spillover of excitation was calculated to range from 16 to 28% of the excitation initially allocated to PSII. These values were not lower in low-light grown plants, indicating that spillover is rather substantial irrespective of the species and growth light conditions. The discrepancy between the results of studies using time-resolved fluorescence analysis and ours should be resolved. It is noteworthy that our electron microscopy analyses revealed that there is enough space for the PSII-PSI megacomplexes (Kim et al. 2023; Yokono et al. 2024) in chloroplasts of *A. odora* irrespective of the growth environments, which have vast inter-grana thylakoids. The margins of appressed thylakoids would be able to accommodate such megacomplexes (Terashima et al. 2025).

When the spillover from PSII to PSI is active, PSII may be protected because Chl* near the PSII reaction center decreases considerably. It is interesting to examine whether shade-tolerant plants grown in low light generally construct thylakoid membranes so as to be resistant to FL-induced photoinhibition. Given that shade-tolerant plants are frequently exposed to strong sunflecks in their natural habitats, such a thylakoid construction would be adaptive.

## 8. Technical problems in quantification of PSI parameters

### *In vitro* electron transport measurement with methyl viologen

When electron transport activities of thylakoid membranes are measured *in vitro*, methyl viologen (MV) is widely used as an electron acceptor from PSI (Izawa 1980). However, this reagent accepts electrons not only from F_A_/F_B_ at the membrane/protein surface, but also from F_X_ in the PSI complex in the membrane because of its hydrophobic/lipophilic nature (Fujii et al. 1990; Sonoike and Terashima 1994). In our earlier studies using cucumber, we isolated thylakoids after tr**e**ating the leaves with light at low temperatures and measured the PSI electron transport rate from ascorbate-reduced 3,6-diaminodurene (DAD) to MV at a concentration of 1 mM. Worse, only the electron transport rates at saturating light were measured (Terashima et al. 1989, 1991b). Thus, we failed to detect PSI photoinhibition at F_A_/F_B_. By contrast, in Terashima et al. (1994), we assessed the electron transport rate from ascorbate-reduced 3,6-diaminodurene to NADP^+^ in the presence of ferredoxin by measuring the absorbance change at 340 nm. Moreover, in this study, the initial slopes obtained at low irradiances were compared. In this way, PSI photoinhibition was found.

### Assessing the PSI redox state *in vivo*

Accurate assessment of the PSI redox state is essential for elucidating the mechanism of PSI photoinhibition. In conventional P700 analyses, Y(ND), Y(I), and Y(NA) are measured by monitoring concentration of P700^+^ with measuring pulses at two wavelengths, at 810 or 830 nm (sample) and 870 nm (reference) with the PAM 101, and at 830 (sample) and 870 nm (reference) with the Dual-PAM (both from Walz, Germany). At first, the sample is kept in the dark to reduce P700, and P_m_ level was measured with a rather intense far-red light and a short saturating flash (SP, for example for 30 ms). Next, actinic light is turned on, and, at the steady state, the P700^+^ level is measured (P). Then by adding the far-red light and a SP, P_m_’ is monitored. Using these data, Y(ND), Y(I) and Y(NA) are obtained as P/P_m_, (P_m_’ – P)/P_m_, and (P_m_ – P_m_’)/P_m_, respectively (Klughammer and Schreiber 1994). Two main concerns for this method have been raised (Fan et al. 2016; Furutani et al. 2022).

First, the components near the P700, plastocyanin (PC) and ferredoxin, show absorbance changes in near infra-red waveband as well. The absorption coefficient of P700^+^ was reported to be 12 mM^-1^ cm^-1^ at 815 nm for spinach chloroplasts (Inoue et al. 1973). Oxidation minus reduced differential absorption coefficient of P700^+^ at (830 nm - 870 nm) calculated from Inoue et al. (1973) is 8.6 mM^-1^ cm^-1^, while those of spinach plastocyanin and ferredoxin would be 0.31 and 0.07 mM^-1^ cm^-1^ (calculated from Schreiber 2017). Although the coefficient of Fd is small, that of PC is considerable. Therefore, the redox levels of PC and Fd influence the apparent redox status of P700^+^/P700 (see below). The Dual KLAS/NIR (Walz) deconvolutes redox levels of these three components, using four wavelength pairs of differential measurements (Schreiber 2017). Oja et al. (2003) measured the redox level of PC using “far-red light titration.”

Second, the method to determine Y(I) itself has been challenged (Furutani et al. 2022). Manipulating the redox level of PC before the intense far-red light and a SP application, they showed that Y(I) values under the similar conditions varied to a considerable extent. During the SP, not only P700 photooxidation but also electron donation to PSI and electron transfer from its acceptor side proceed rapidly. Values calculated using the SP therefore do not necessarily reflect the PSI redox state even under the steady illumination conditions (Furutani et al. 2022). Thus, Y(I) should be regarded as an operational index that reflects the balance between electron donation to PSI and electron transfer from its acceptor side during the SP rather than direct measure of the steady-state PSI quantum yield. The estimation of Y(NA) therefore suffers from the same problem.

Here, we raise another concern. It is now widely recognized that fluorescence signals from the leaf do not represent photosynthetic activity of the whole leaf, because only the fluorescence signal from the shallow part of the leaf is detected, especially when the measuring beam is blue or red (Oguchi et al. 2011; Terashima et al. 2009). Concerning the near infra-red light for PSI measurements, leaf absorptance in this waveband is only a few percent (Gates 1980), because there are few pigments absorbing light in near infra-red. Due to its diffusive nature of mesophyll structure, the near infra-red light is well scattered in the leaf sample, and it is believed that absorbance change in P700^+^ is rather evenly detected throughout the leaf (Fan et al. 2016).

Let us assume that P700^+^ is evenly detected. However, the actinic light for photosynthesis is absorbed unevenly. The chloroplasts near the illuminated side of the leaf absorb more light on a unit chlorophyll basis than those located deep in the leaf. Although the photosynthetic properties of chloroplasts change from sun-type to shade-type with the leaf depth, this gradient of sun-shade chloroplast properties is usually less steep than the light absorption gradient (Terashima et al. 2009). Then, we can expect that Y(ND) is greater and Y(I) is smaller in the chloroplasts near the illuminated surface than in those deep in the leaf. On the absorbed photon basis, therefore, Y(I) tends to be overestimated and Y(ND) tends to be underestimated. This would explain the tendency that the electron transport rate through PSI, ETR(I), is overestimated, as has been warned (Fan et al. 2016).

When PSI photoinhibition proceeds progressively, Y(I), and thereby ETR(I), tend to be further overestimated. Thus, the changes in P_m_ should be incorporated (Grebe et al. 2024; Kono et al. 2014; Takeuchi et al. 2025b).

The redox state of Fd obtained by deconvolution of near infra-red measurement data with systems like the Dual KLAS/NIR (Walz) provides a comprehensive view of PSI acceptor-side reduction (Klughammer and Schreiber 2016; Schreiber 2017). Fd redox state can be followed continuously under steady illumination without applying saturating flashes. After the chilling treatment, Fd reoxidation became markedly slower in the chilling-sensitive cucumber cultivar than in the chilling-tolerant cultivar (Takeuchi et al. 2025a). Fd reoxidation kinetics provide an integrated measure of electron efflux from reduced Fd (Furutani et al. 2023; Kadota et al. 2019). Fd oxidation reflects the combined contributions of the CBB cycle, photorespiration, FLV, CEF-PSI, and other alternative electron transport pathways. The very small oxidized minus reduced absorption coefficient of Fd in near infra-red currently prevents detailed analysis of Fd use. However, in principle, the oxidation kinetics may be expressed by addition of several exponential decay components.

The capacities for P700 and Fd oxidation are sensitive indicators of PSI redox regulation at low temperatures and can be used for early assessment of chilling sensitivity. Takeuchi et al. (2022) reported that the accumulation of P700⁺ after 3 h of chilling stress predicted subsequent chilling injury among cucumber cultivars. The predictive value of these measurements, however, depends on physiological context: in cucumber grown under different Fe supplies, oxidative injury was more pronounced under Fe-rich conditions despite comparable levels of P700 oxidation, consistent with amplification of ROS-mediated damage by Fe-dependent Fenton chemistry (Takeuchi et al. 2024).

It should be noted again that calculation of the electron flux through Fd suffers from the above-mentioned problem. Even when the average oxidation rate throughout the leaf can be measured rather correctly upon cessation of the actinic light, it is expected that the Fd reduction level and oxidation rate are lower and slower in the chloroplasts far from the illuminated surface. However, the electron flux through Fd is greater in the chloroplasts located near the illuminated surface. Thus, it is problematical to multiply the average Fd reduction level and reoxidation rate obtained for Fd with the photon flux density allocated to PSI.

## 9. Future directions in PSI photoinhibition research

### 9.1 Species differences and unresolved molecular mechanisms

#### PSI photoinhibition at low temperatures

Research on chilling-induced PSI photoinhibition has advanced substantially using cucumber as a model. An important question is whether the sequence of events leading to chilling-induced PSI photoinhibition described here also occurs in plant species other than cucumber. As has been discussed, PSI protection at low temperature depends on different combinations of processes that restrict electron flow to PSI and relieve reduction on its acceptor side. These include PSII photoinhibition, photosynthetic control, CEF-PSI, FLV, etc. The relative importance of these processes is likely to vary among plant species and growth environments. In cucumber, the chilling-induced weak association of CF_1_ with the thylakoid membrane uncouples ATP synthesis and impairs ΔpH formation and thereby photosynthetic control. The resulting loss of donor-side regulation makes PSI protection increasingly dependent on acceptor-side processes, particularly NDH (Takeuchi et al. 2025a, 2026). By contrast, in plants that rapidly downregulate PSII activity, electron flow into PSI is suppressed, potentially reducing their dependence on acceptor-side processes such as NDH. Future comparative studies should therefore examine how different species combine donor-side regulation with acceptor-side electron dissipation, rather than evaluating each protective process in isolation.

Further studies are needed to determine how changes in NDH activity at low temperature affect chloroplast function. NDH is embedded in a broader chloroplast redox network: its activity is modulated by several regulatory factors (Ermakova et al. 2024b; He et al. 2026; Kambakam et al. 2016; Kohzuma et al. 2026, preprint; Lascano et al. 2003; Nikkanen et al. 2018, 2020; Wang et al. 2006; Zhang Q et al. 2024; Zheng et al. 2025), and changes in its activity can, in turn, affect chloroplast metabolism, including NADPH homeostasis (Nashilevitz et al. 2010; Zhang Q et al. 2024; Zhang Y et al. 2023). Understanding the consequences of altered NDH activity during chilling will therefore require examining both its regulation and its metabolic effects within this network.

Studies of PSI photoinhibition under stresses other than chilling have identified diverse redox-regulatory and electron-consuming processes that contribute to PSI protection, some of which operate beyond the linear photosynthetic electron transport chain. These include FTR, nitrogen and sulfur assimilation, the malate valve, mitochondrial respiration, and plastid terminal oxidase (PTOX) (Backhausen et al. 1994; Hani et al. 2025; Hanke et al. 2008; Höhner et al. 2021; Konishi and Noguchi 2025; Krämer et al. 2024; Messant et al. 2024; Yamada et al. 2024). It remains to be determined to what extent these pathways protect PSI at low temperature, whether they relieve acceptor-side reduction directly or indirectly, and how they interact with the protective mechanisms described above. Another important question is which component of PSI is damaged first during chilling-induced photoinhibition.

Recent evidence indicates that light-harvesting complex I (LHCI) may be one of the early targets of PSI photoinhibition (Takagi et al. 2026). Oxidative damage to LHCI has been reported during chilling-induced PSI photoinhibition (Alboresi et al. 2009; Rajagopal et al. 2005). Oxidatively modified amino acid residues have also been identified in LHCI proteins, including residues near bound chlorophylls and on the protein surface (Kale et al. 2020, 2022). Disruption of excitation-energy transfer from LHCI to the PSI core could prolong chlorophyll excitation within LHCI, increasing the probability of ^3^Chl* formation and subsequent ^1^O_2_ generation (Carbonera et al. 2005; Kale et al. 2020; Rajagopal et al. 2005). Collectively, these observations suggest that LHCI damage may be an early event during chilling-induced PSI photoinhibition (Sonoike 1999; Takeuchi et al. 2026). This possibility may also help explain the early destabilization of the PSI–NDH supercomplex. Lhca5 and Lhca6 mediate the association between PSI and NDH, and damage to these subunits during chilling stress has been suggested to contribute to destabilization of the PSI–NDH supercomplex (Takeuchi et al. 2026).

#### FL and rSPs induced PSI photoinhibition: Points of tension concerning the division of labor between the two CEF-PSI pathways

Several unresolved issues are summarized in Table S2 and discussed in Text S2. Three points deserve mentioning here. The first concerns gating itself (see § 5.3). Two observations weigh against the idea that the pmf holds NDH back in a leaf: a light-induced pmf did not inhibit NDH in the way that added ATP did (Fisher et al. 2019), and NDH-dependent CEF-PSI operates in the H_2_O_2_-accumulating line GO5 despite a pmf about twice that of the wild type (Strand et al. 2015). By contrast, collapsing the pmf with uncouplers increased the NDH-dependent post-illumination fluorescence rise in thylakoids from spinach, *A. hybridus* and *A. thaliana*; the increase was smaller in *A. thaliana*, but this reflects the larger share of the antimycin A-sensitive pathway in its fluorescence rise (Strand et al. 2017). Neither of the two observations is decisive: gating defined by the net driving force rather than by Δp alone accommodates GO5, and Fisher et al. (2019) themselves expect NDH to become limited at a higher pmf (Table S2, row 1). We therefore adopt gating as a working hypothesis. It adds to, rather than replaces, the established reasons why NDH contributes little under red high light (low abundance, slow turnover and competition with PSII for oxidized PQ; Ermakova et al. 2024a), and it explains why the balance should shift with light quality as well as with irradiance. Two mutant studies point the same way: in *hope2*, the NDH pathway contributed most under low light, where back-pressure from the pmf is small (Degen et al. 2023), and in *crr2-2* under red FL, NDH relieved the acceptor side of PSI in the low-light phases but not in the high-light phases, possibly because a large ΔpH and a reduced PQ pool favor its reverse reaction (Zhou et al. 2023). In a recent review, Shikanai et al. (2025) also proposed that back-pressure from the pmf, possibly through a reverse reaction, limits NDH, so that NDH contributes to pmf formation mainly under low light. Two qualifications remain: an oxidized chain favors NDH only while enough reduced donors are left (Joët et al. 2002; Table S2, row 2), and ATP feedback could produce a similar division of roles (Fisher et al. 2019; Table S2, row 3). The sign of the net driving force has never been determined *in vivo*, and a direct test is overdue (Table S3, predictions 1–3).

The second point concerns the pulse-type regime. Under repeated 1-s pulses of approximately 3,000 µmol photons m⁻² s⁻¹, *pgr5* and *ndho* lose P_m_ to a similar extent as the wild type (Rantala et al. 2020), as in our FR-free controls, in which P_m_ fell to approximately 70% in all genotypes (Kono et al. 2017): the loss of either pathway does not increase PSI photoinhibition under this kind of burst. In pulse regimes, genotype dependence appears only when FR is added, and it then runs through NDH, whereas in the 2-min regime it runs through PGR5. This is what both the masking hypothesis and the gating hypothesis predict, although the mechanism of photoinhibition may differ between the two types of regimes (Sonoike 2025; Table S2, row 6). The third point concerns the molecular role of PGR5, which remains uncertain (Table S2, row 8). A cycle at steady state returns to PSI exactly as many electrons as it removes and therefore cannot keep P700 oxidized by itself; its protective effect must come from the protons it moves, either as extra ATP that relieves the acceptor side or as lumen acidification that engages photosynthetic control (Vilyanen and Kozuleva 2025). Our account in § 5.2 uses both routes and requires only that the PGR5-dependent pathway builds the pmf quickly from a large electron flux without a proton pump of its own. If PGR5 proves to be a regulator rather than a carrier, the "capacity" entries of Table 1 describe the pathway rather than the protein, and the rescue of *pgr5* by FR is read as FR removing the need for that regulation; the conclusion is unchanged.

#### Testable predictions

The hypotheses make predictions that can be tested with existing methods (Table S3). The central one is that FR shifts the thermodynamic balance of NDH toward the forward reaction: the pmf from the electrochromic shift and the redox states of P700, plastocyanin and the PQ pool, measured together in leaves, should show that the net driving force is more positive under FR than in the high-light phase of red FL. The Δp dependence of NDH itself can be measured in coupled thylakoids under FR when the pmf is set without ATP, for example by an acid–base transition (Jagendorf and Uribe 1966); on the genetic side, FR should fail to rescue the *pgr5 crr* double mutant. If predictions (1) to (5) fail to show a wavelength-dependent constraint on NDH, the gating hypothesis should be abandoned, and the division of roles should be explained by capacity, ATP feedback and activation alone. The masking hypothesis, however, would survive: FR oxidizes P700 by the excitation balance, this removes the specific need for PGR5, and NDH provides the low-flux recycling that remains necessary.

### 9.2 Need of ecophysiological studies

#### Ecological implications

Two lines of evidence bear on which pathway matters in nature. The first concerns light intensity. Under FL that reproduced the irradiance, but not the spectrum, measured inside a maize canopy at 100-ms resolution, the NDH-deficient mutants *ndho* and *ndhm* lost on average approximately 63% of their shoot dry weight and the Burren *ndhG* variant approximately 10%, whereas neither was affected under constant light or slow square-wave fluctuations (Theeuwen et al. 2022, preprint). When the same lines were grown under a similar canopy-like regime, the NDH knockouts showed lower linear electron flow, pmf and biomass than under constant light, whereas Fv/Fm was unaffected (L.A. Vorbach, unpublished report). PSI photoinhibition was not measured in either study. The second concerns light quality. Canopy and understory light is enriched in FR, the red:far-red photon ratio falling from about 1.2 in daylight to about 0.1 under a dense canopy (Casal 2013; Franklin and Quail 2010; Smith 1982), and in field-grown *Erigeron annuus* (L.) Pers. and *Commelina communis* L., adding FR to red FL largely prevented PSI photoinhibition (Kono et al. 2017). The two lines support two different statements, that NDH matters for growth under canopy-like fluctuations and that NDH-dependent protection is engaged by FR; their combination is a hypothesis: in the FR-rich, rapidly fluctuating light of canopies, the NDH-dependent pathway may be the ecologically relevant route of PSI protection, and the red LED regimes of most laboratory studies may have overestimated the role of PGR5 and underestimated that of NDH; testing this requires PSI photoinhibition to be measured under natural spectra (Table S3, prediction 11).

To understand PSI photoinhibition and its suppression mechanisms in extant plants, we should not forget that plants have evolved under natural environmental conditions. Studies of PSI photoinhibition under FL have become increasingly active, and PSI is now considered to be vulnerable to FL. However, we have found that PSI photoinhibition is almost completely suppressed by far-red light at a level comparable to that present in natural sunlight. Despite this clear result, the extent of PSI photoinhibition under fluctuating natural light has scarcely been examined. Careful field studies are needed to demonstrate the suppressive effect of far-red light on PSI photoinhibition under natural conditions. Another important problem is that we do not know the detailed roles of far-red light in photosynthesis. We may call this situation “Far-red drop.” In addition to the problems discussed above, far-red light is not included in the actinic light of most conventional photosynthetic instruments. The latest LI-6800 (LI-COR) allows far-red light to be added, but this is not the default setting. With such excitation light, insufficient excitation of PSI can cause over-reduction of the PQ pool, resulting in a decrease in Y(II) (Kono et al. 2022). Improvement of the spectral composition of excitation light is therefore essential.

Based on the effects of far-red light supplementation, it has been proposed that photosynthetically active radiation (PAR), conventionally defined as 400–700 nm, should be extended to 750 nm (Zhen and Bugbee 2020). The conventional definition of PAR as 400–700 nm was partly based on the fact that green leaves strongly absorb light in this wavelength range. The absorptance of green leaves at 700 nm is approximately comparable to that at 550 nm, and decreases sharply at longer wavelengths and becomes a few percent at 750 nm. Considering the first law of photochemistry, “only absorbed photons can do photochemical work,” the low absorptance of green leaves between 700 and 750 nm suggests that a simple extension of PAR to 750 nm is not appropriate (Levine et al. 2026).

In this review, we have examined the mechanisms of PSI photoinhibition in green leaves and the mechanisms that suppress it. In plant factories and other controlled environments, we can now freely use light sources with a wide variety of spectral compositions. In a “controlled” light environment, any wavelength composition can be selected as long as the light is absorbed by plants. However, whenever plant functions are considered, it is important to keep in mind that extant plants have evolved under natural environmental conditions.

## Supplementary Information

Text S1. Affinity of CEF-PSI pathways to Fd

Text S2. Points of tension and open questions

Text S3. Experiments to test the masking and redox–Δp gating hypotheses

## Acknowledgement

We thank Professor Chikahiro MIYAKE for closely reading some parts of the text and providing useful suggestions. We dedicate this review paper to the late Professor Shizuo YOSHIDA (who passed away on January 9, 2024), a pioneer in plant membrane biology at low temperatures, particularly known for his studies on the role of plasma membrane H^+^-ATPase, and to the late Professor Sakae KATOH (who passed away on May 21, 2025), a pioneer in photosynthesis research, particularly known for purification of plastocyanin and determination of its role. One of the authors (I.T.) was enlightened by Prof. Yoshida’s ecophysiological approach. Prof. Katoh taught him how to approach thylakoid reactions, with incredible patience.

## Declarations

### Author contributions

IT and MK conceived the review. RO performed the literature search and data analysis for the meta-analysis (§ 3, Table S1 and Fig. 3). KT, MK, IT and RO searched the literature for their sections and drafted the manuscript: KT drafted §§ 1, 4 and 6 and the paragraphs on chilling in § 9.1; MK drafted §§ 2 and 5, the rest of § 9.1, the paragraph “Ecological implications” in § 9.2, Supplementary Tables S2 and S3, and Supplementary Texts S1–S3; IT drafted the Introduction, §§ 7 and 8, and the rest of § 9.2. KS and IT compiled the drafts. All authors critically revised the manuscript and read and approved the final version.

### Data availability

Data in this article are available from the corresponding authors upon request.

## Abbreviations

A_0_: primary electron acceptor of PSI (Chl *a*)
A_1_: secondary electron acceptor of PSI (phylloquinone)
APX: ascorbate peroxidase
CBB cycle: Calvin–Benson–Bassham cycle
CEF-PSI: cyclic electron flow around PSI
CF_1_: catalytic portion (coupling factor 1) of the chloroplast ATP synthase
Chl: chlorophyll
Cyt *b*_6_*f*: cytochrome *b*_6_*f* complex
DAD: 3,6-diaminodurene
DCMU: 3-(3,4-dichlorophenyl)-1,1-dimethylurea
ECS: electrochromic shift
ETR(I): electron transport rate through PSI
F_A_, F_B_: terminal iron–sulfur clusters of PSI, bound to PsaC
Fd: ferredoxin
Fe–S: iron–sulfur
FL: fluctuating light
FLV: flavodiiron protein
FNR: ferredoxin–NADP^+^ reductase
FR: far-red light
FTR: ferredoxin–thioredoxin reductase
F_v_/F_m_: maximum quantum yield of PSII photochemistry
F_X_: iron–sulfur cluster of PSI, bound to PsaA and PsaB
*g*H^+^: proton conductivity of the thylakoid ATP synthase
HL: high light
*K*_pi_: rate constant of PSI photoinhibition
*K*_pi_/*I*: rate constant of PSI photoinhibition divided by irradiance
LED: light-emitting diode
LHCI: light-harvesting complex I
LL: low light
MV: methyl viologen
NDH: chloroplast NADH dehydrogenase-like complex
NPQ: non-photochemical quenching of chlorophyll fluorescence
NTRC: NADPH-dependent thioredoxin reductase C
^1^O_2_: singlet oxygen
O_2_^•^⁻: superoxide radical anion
P700: reaction center chlorophyll of PSI
PAR: photosynthetically active radiation
PC: plastocyanin
PGR5: PROTON GRADIENT REGULATION 5
PGRL1: PGR5-LIKE PHOTOSYNTHETIC PHENOTYPE 1
PIFR: post-illumination fluorescence rise
P_m_: maximum amount of photooxidizable P700
pmf: proton motive force
PQ: plastoquinone
PQH_2_: plastoquinol
PSI: photosystem I
PSII: photosystem II
PTOX: plastid terminal oxidase
RISE: reduction-induced suppression of electron flow
ROS: reactive oxygen species
rSPs: repetitive saturating pulses
SOD: superoxide dismutase
SP: saturating pulse
Trx: thioredoxin
WT: wild type
Y(I): quantum yield of PSI photochemistry
Y(II): effective quantum yield of PSII photochemistry
Y(NA): quantum yield of non-photochemical energy dissipation due to acceptor-side limitation of PSI
Y(ND): quantum yield of non-photochemical energy dissipation due to donor-side limitation of PSI
Y(NO): quantum yield of non-regulated energy dissipation in PSII
Y(NPQ): quantum yield of regulated non-photochemical energy dissipation in PSII
Δ*E*_h_: difference in redox potential between the electron acceptor (PQ) and donor (Fd) of NDH
Δ*p*: proton motive force expressed in mV
ΔpH: pH difference across the thylakoid membrane
ΔΨ: membrane potential across the thylakoid membrane

## Supplementary Materials

### 1. Supplementary Table S1 and Explanatory Notes

We summarize the results of PSI photoinhibition studies. Experimental conditions and results are shown as follows.

- Samples: Intact leaves, detached leaves, leaf discs, or isolated thylakoids are categorized.
- Species: Species names (common names) are listed. Experiments using the *pgr5* mutant of *Arabidopsis thaliana* are shown in bold.
- Photoinhibition temperatures: *T* (°C).
- Photoinhibition light regimes: Light treatments used are categorized into continuous (C), fluctuating (F) and intermittent (I) lighting regimes.
- Photoinhibition irradiances: The irradiance level (photosynthetic photon flux density) in μmol m^-2^ s^-1^ is shown. For fluctuating light treatments, all irradiance levels employed are shown like A/B/C.
- The photoinhibition photon-exposure, or cumulative irradiance, in mol m^-2^ is listed.
- The residual PSI activity after the treatment is shown as %PSI.
- The rate constant of PSI photoinhibition divided by irradiance, *K*_pi_/*I* ( (mol photons)^-1^ m^2^) is shown. For *K*_pi_ calculation, see main text.

**Supplementary Table S1.** Summary of PSI photoinhibition studies.

| **Paper** | **Samples** | **Species** | ***T*** | **Light** | **Irradiance** | **Exposure** | **PSI %** | ***K*<sub>pi</sub>/*I*** |
|---|---|---|---|---|---|---|---|---|
| Satoh 1970a | thylakoids | *Spinacia oleracea* L. (Spinach) | 25 | C | 240 | 0.86 | 42 | 1.004 |
| Satoh 1970a | thylakoids | Spinach | 4 | C | 240 | 0.86 | 88 | 0.148 |
| Sonoike 1995 | thylakoids | *Cucumis sativus* L. (Cucumber) | 5 | C | 300 | 5.4 | 50 | 0.128 |
| Sonoike 1995 | thylakoids | Cucumber | 5 | C | 100 | 1.8 | 63 | 0.257 |
| Sonoike 1995 | thylakoids | Cucumber | 5 | C | 200 | 3.6 | 40 | 0.255 |
| Sonoike 1995 | thylakoids | Cucumber | 25 | C | 200 | 1.08 | 70 | 0.330 |
| Sonoike 1995 | thylakoids | Spinach | 25 | C | 300 | 1.62 | 28 | 0.797 |
| Sonoike 1995 | thylakoids | Spinach | 5 | C | 100 | 1.8 | 28 | 0.717 |
| Sonoike 1995 | thylakoids | Spinach | 25 | C | 100 | 0.54 | 33 | 2.053 |
| Sonoike 1999 | thylakoids | Cucumber | 0 | C | 110 | 1.98 | 38 | 0.489 |
| Sonoike 1999 | thylakoids | Cucumber | 4 | C | 110 | 1.98 | 42 | 0.438 |
| Sonoike 1999 | thylakoids | Cucumber | 0 | C | 60 | 1.08 | 45 | 0.739 |
| Sonoike 1999 | thylakoids | Cucumber | 4 | C | 60 | 1.08 | 48 | 0.680 |
| Terashima et al. 1994 | Leaf | Cucumber | 4 | C | 220 | 5.94 | 25 | 0.233 |
| Havaux and Davaud 1994 | Leaf disc | *Solanum tuberosum* L. (Potato) | 3 | C | 3500 | 4.2 | 75 | 0.068 |
| Havaux and Davaud 1994 | Leaf disc | Potato | 2 | C | 900 | 8.1 | 57 | 0.069 |
| Havaux and Davaud 1994 | Leaf disc | Potato | 2 | C | 3700 | 6.66 | 64 | 0.067 |
| Terashima et al. 1998 | Leaf intact | Cucumber | 5 | C | 200 | 2.16 | 60 | 0.236 |
| Terashima et al. 1998 | Leaf intact | Cucumber | 5 | C | 200 | 4.32 | 40 | 0.212 |
| Tjus et al. 1998 | Detached leaf | *Hordeum vulgare* L. (Barley) | 4 | C | 100 | 2.88 | 80 | 0.077 |
| Ivanov et al. 1998 | Detached leaf | *Secale cereale* L. (Winter rye) | 5 | C | 1600 | 23.0 | 66 | 0.018 |
| Ivanov et al. 1998 | Detached leaf | Winter rye | 5 | C | 1600 | 23.0 | 69 | 0.016 |
| Ivanov et al. 1998 | Detached leaf | Winter rye | 5 | C | 1600 | 23.0 | 92 | 0.004 |
| Ivanov et al. 1998 | Detached leaf | Winter rye | 5 | C | 1600 | 23.0 | 59 | 0.023 |
| Ivanov et al. 1998 | Detached leaf | Winter rye | 5 | C | 1600 | 23.0 | 108 | -0.003 |
| Teicher et al. 2000 | Leaf | Barley | Chilling | C | 150 | 30.2 | 88 | 0.004 |
| Teicher et al. 2000 | Leaf | Barley | Chilling | C | 150 | 30.2 | 90 | 0.004 |
| Kornyeyev et al. 2003 | Leaf disc | *Gossypium hirsutum* L. (Cotton) | 10 | C | 500 | 7.2 | 50 | 0.096 |
| Kornyeyev et al. 2003 | Leaf disc | Cotton | 10 | C | 500 | 7.2 | 67 | 0.056 |
| Zhang and Scheller 2004 | Leaf | *Arabidopsis thaliana* (L.) Heynh. WT.<br>(*A. thaliana* WT) | 4 | C | 150 | 2.7 | 71 | 0.126 |
| Ivanov et al. 2012 | Leaf | *A. thaliana* WT | 5 | **C** | 600 | 4.32 | 33 | 0.260 |
| Ivanov et al. 2012 | Leaf | *A. thaliana* WT | 5 | C | 600 | 4.32 | 63 | 0.107 |
| Cazzaniga et al. 2012 | Leaf intact | *A. thaliana* WT | 8 | C | 550 | 5.94 | 65 | 0.073 |
| Zhang et al. 2014 | Leaf disc | Cucumber | 6 | C | 400 | 13.0 | 51 | 0.052 |
| Zhang et al. 2014 | Leaf disc | Cucumber | 6 | C | 400 | 13.0 | 36 | 0.079 |
| Zhang et al. 2016 | Leaf intact | Cucumber | 6 | C | 200 | 4.32 | 33 | 0.257 |
| Zhang et al. 2016 | Leaf intact | Cucumber | 6 | C | 200 | 4.32 | 63 | 0.107 |
| Havaux and Davaud 1994 | Leaf disc | Potato | 23 | C | 3500 | 4.2 | 96 | 0.010 |
| Havaux and Davaud 1994 | Leaf disc | Potato | 25 | C | 3700 | 6.66 | 82 | 0.030 |
| Sonoike 1999 | Detached leaf | Cucumber | 20 | C | 200 | 3.6 | 100 | 0.000 |
| Tikkanen et al. 2014 | Leaf | *A. thaliana* WT | RT | C | 800 | 5.76 | 100 | 0.000 |
| **Tikkanen et al. 2014** | **Leaf** | ***A. thaliana pgr5*** | **RT** | **C** | **800** | **5.76** | **80** | **0.039** |
| Tikkanen et al. 2014 | Leaf | *A. thaliana* WT | RT | C | 800 | 5.76 | 100 | 0.000 |
| **Tikkanen et al. 2014** | **Leaf** | ***A. thaliana pgr5*** | **RT** | **C** | **800** | **5.76** | **50** | **0.120** |
| **Lima-Melo et al. 2019** | **Leaf intact** | ***A. thaliana pgr5*** | **23** | **C** | **1000** | **0.9** | **60** | **0.568** |
| Huang et al. 2016 | Leaf intact | *Psychotria henryi* H. Lév. | RT | C | 2258 | 13.5 | 75 | 0.021 |
| Oguchi et al. 2021 | Leaf intact | *Capsicum annuum* L. | RT | C | 950 | 4.56 | 96 | 0.009 |
| Oguchi et al. 2021 | Leaf intact | *C. annuum* | RT | C | 950 | 4.56 | 92 | 0.018 |
| **Tiwari et al. 2024** | **Detached leaf** | ***A. thaliana pgr5*** | **RT** | **C** | **850** | **6.12** | **34** | **0.177** |
| Tiwari et al. 2024 | Detached leaf | *A. thaliana* WT | RT | C | 850 | 6.12 | 100 | 0.000 |
| Tsuyama and Kobayashi 2009 | Leaf | *A. thaliana* WT | RT | F | 8000/60 | 12.3 | 72 | 0.027 |
| Tsuyama and Kobayashi 2009 | Leaf | *A. thaliana crr2-2* | RT | F | 8000/60 | 12.3 | 33 | 0.090 |
| Kono et al. 2014 | Leaf | *A. thaliana* WT | RT | F | 240/30 | 0.340 | 89 | 0.343 |
| **Kono et al. 2014** | **Leaf** | ***A. thaliana pgr5*** | **RT** | **F** | **240/30** | **0.340** | **62** | **1.405** |
| Huang et al. 2016 | Leaf intact | *P. henryi* | RT | F | 1294/100 | 2.51 | 95 | 0.019 |
| Huang et al. 2016 | Leaf intact | *P. henryi* | RT | F | 2258/94 | 7.08 | 93 | 0.010 |
| Kono et al. 2017 | Leaf | *A. thaliana* WT | RT | F | 1200/30 | 0.84 | 72 | 0.399 |
| Kono et al. 2017 | Leaf | *A. thaliana* WT | RT | F | 1200/135 | 1.54 | 85 | 0.106 |
| Kono et al. 2017 | Leaf | *A. thaliana* WT | RT | F | 1200+FR(25.6 W m<sup>-2</sup>) | 0.64 | 97 | 0.048 |
| Kono et al. 2017 | Leaf | *A. thaliana* WT | RT | F | 1200+FR(12.1 W m<sup>-2</sup>) | 0.64 | 95 | 0.080 |
| Kono et al. 2017 | Leaf | *A. thaliana* WT | RT | F | 1200+FR(3.93 W m<sup>-2</sup>) | 0.64 | 86 | 0.236 |
| Tikkanen and Grebe 2018 | Leaf | *A. thaliana* WT | RT | F | 5000/100/80 | 3.73 | 30 | 0.323 |
| Yang et al. 2019 | Leaf | *Prunus cerasoides* Buch.-Ham. ex D. Don | RT | F | 1809/59 | 1.12 | 95 | 0.046 |
| Yang et al. 2019 | Leaf | *P. cerasoides* | RT | F | 1809/59 | 1.12 | 84 | 0.156 |
| Lempiäinen et al. 2022 | Leaf intact | *A. thaliana* WT | 25 | F | 3200/100/35 | 2.17 | 39 | 0.434 |
| Lempiäinen et al. 2022 | Leaf intact | *A. thaliana* WT | 25 | F | 3200/100/35 | 4.34 | 15 | 0.437 |
| Kono et al. 2017 | Leaf | *A. thaliana* WT | RT | I | 1200 | 0.64 | 63 | 0.722 |
| Takagi et al. 2017b | Leaf | *Helianthus annuus* L. (Sunflower) | 25 | I | 5000 | 1.80 | 35 | 0.583 |
| Takagi et al. 2017b | Leaf | *Zea mays* L. | 25 | I | 5000 | 1.80 | 38 | 0.538 |
| Takagi et al. 2017b | Leaf | *Rhipsalis baccifera* (J.S.Muell.) Stearn | 25 | I | 5000 | 1.80 | 57 | 0.312 |
| Takagi et al. 2017b | Leaf | *Eucalyptus globulus* Labill. | 25 | I | 5000 | 1.80 | 44 | 0.456 |
| Takagi et al. 2017b | Leaf | *Olea europaea* L. | 25 | I | 5000 | 1.80 | 46 | 0.431 |
| Takagi et al. 2017b | Leaf | *Quercus glauca* Thunb. | 25 | I | 5000 | 1.80 | 37 | 0.552 |
| Takagi et al. 2017b | Leaf | *Camellia japonica* L. | 25 | I | 5000 | 1.80 | 75 | 0.160 |
| Takagi et al. 2017b | Leaf | *Osmanthus fragrans* (Thunb.) Lour. | 25 | I | 5000 | 1.80 | 41 | 0.495 |
| Takagi et al. 2017b | Leaf | *Acer palmatum* Thunb. | 25 | I | 5000 | 1.80 | 60 | 0.284 |
| Takagi et al. 2017b | Leaf | *Populus nigra L.* | 25 | I | 5000 | 1.80 | 59 | 0.293 |
| Takagi et al. 2017b | Leaf | *Prunus* × *yedoensis* Matsum. ‘Somei-yoshino’ | 25 | I | 5000 | 1.80 | 51 | 0.374 |
| Takagi et al. 2017b | Leaf | *Sequoia sempervirens* (Lamb.) Endl. | 25 | I | 5000 | 1.80 | 89 | 0.065 |
| Takagi et al. 2017b | Leaf | *Pinus thunbergii* Parl. | 25 | I | 5000 | 1.80 | 96 | 0.023 |
| Takagi et al. 2017b | Leaf | *Cedrus deodara* (Roxb. ex D.Don) G.Don | 25 | I | 5000 | 1.80 | 92 | 0.046 |
| Takagi et al. 2017b | Leaf | *Juniperus chinensis* L. | 25 | I | 5000 | 1.80 | 94 | 0.034 |
| Takagi et al. 2017b | Leaf | *Podocarpus macrophyllus* (Thunb.) Sweet | 25 | I | 5000 | 1.80 | 90 | 0.059 |
| Takagi et al. 2017b | Leaf | *Cycas revoluta* Thunb. | 25 | I | 5000 | 1.80 | 95 | 0.028 |
| Takagi et al. 2017b | Leaf | *Ginkgo biloba* L. | 25 | I | 5000 | 1.80 | 79 | 0.131 |
| Takagi et al. 2017b | Leaf | *Matteuccia struthiopteris* (L.) Tod. | 25 | I | 5000 | 1.80 | 73 | 0.179 |
| Takagi et al. 2017b | Leaf | *Selaginella moellendorffii* Hieron. | 25 | I | 5000 | 1.80 | 70 | 0.198 |
| Takagi et al. 2017b | Leaf | *Phlebodium aureum* (L.) J.Sm. | 25 | I | 5000 | 1.80 | 79 | 0.131 |
| Takagi et al. 2017b | Leaf | *Nephrolepis cordifolia* (L.) C.Presl | 25 | I | 5000 | 1.80 | 80 | 0.124 |
| Takagi et al. 2017b | Leaf | *Equisetum arvense* L. | 25 | I | 5000 | 1.80 | 87 | 0.077 |
| Takagi et al. 2017b | Leaf | *Pyrrosia lingua* (Thunb.) Farw. | 25 | I | 5000 | 1.80 | 76 | 0.152 |
| Takagi et al. 2017b | Leaf | *Dryopteris erythrosora* (D.C.Eaton) Kuntze | 25 | I | 5000 | 1.80 | 78 | 0.138 |
| Takagi et al. 2017b | Leaf | *Botrychium ternatum* (Thunb.) Sw. | 25 | I | 5000 | 1.80 | 90 | 0.059 |
| Takagi et al. 2017b | Leaf | *Marchantia polymorpha L.* | 25 | I | 5000 | 1.80 | 70 | 0.198 |
| Takagi et al. 2017b | Leaf | *Conocephalum conicum* (L.) Dum. | 25 | I | 5000 | 1.80 | 48 | 0.414 |
| Takagi et al. 2017b | Leaf | *Polytrichum juniperinum* Willd. ex Hedw. | 25 | I | 5000 | 1.80 | 96 | 0.023 |
| Takagi et al. 2017b | Leaf | *Hypopterygium fauriei* Besch. | 25 | I | 5000 | 1.80 | 62 | 0.266 |
| Sejima et al. 2014 | Leaf | Sunflower | RT | F | 21400/1400 | 28.80 | 85 | 0.006 |
| Sejima et al. 2014 | Leaf | Sunflower | RT | F | 20580/580 | 16.99 | 65 | 0.025 |
| Sejima et al. 2014 | Leaf | Sunflower | RT | F | 20200/200 | 11.52 | 21 | 0.135 |
| Sejima et al. 2014 | Leaf | Sunflower | RT | I | 20000 | 8.64 | 10 | 0.267 |
| Zivcak et al. 2015 | Leaf | *Triticum aestivum* L. (Wheat) | RT | I | 15000 | 0.41 | 45 | 1.972 |
| Takagi et al. 2016 | Isolated chloroplasts | Spinach | 25 | I | 20000 | 2.16 | 15 | 0.878 |
| Takagi et al. 2019 | Leaf | Wheat | 25 | I | 20000 | 2.16 | 35 | 0.486 |
| Takagi et al. 2019 | Leaf | Wheat | 25 | I | 20000 | 2.16 | 58 | 0.256 |
| Furutani et al. 2023 | Leaf intact | *A. thaliana* WT | RT | I | 15000 | 0.81 | 28 | 1.554 |
| **Furutani et al. 2023** | **Leaf intact** | ***A. thaliana pgr5*** | **RT** | **I** | **15000** | **0.81** | **28** | **1.556** |

Satoh (1970a) irradiated the thylakoid samples with light from tungsten lamps at 12000 lux, which may be converted to PPFD at 240 µmol m<sup>-2</sup> s<sup>-1</sup>.

## 2. Supplementary Text S1

### Affinity of CEF-PSI pathways to Fd

This text expands the paragraph "Interaction with Fd" in § 5.1. It gives the sources and conditions of the values quoted there, compares the two CEF-PSI pathways with the other Fd-dependent enzymes of the stroma, summarizes what is known about the amount and redox state of Fd *in vivo*, and lists the open questions, which are taken up in Table S3, prediction 5.

### How the two pathways bind Fd

For NDH, the Fd-binding subunit is NdhS (CRR31), a protein with an SH3-like fold (Yamamoto et al. 2011). In ruptured *Arabidopsis* chloroplasts, NDH-dependent PQ reduction is almost saturated at 3 µM Fd in the wild type, whereas in *ndhs* it remains low even at 20 µM Fd; NdhS is thus required for high-affinity binding, although it is not essential for NDH activity (Yamamoto et al. 2011). The binding is electrostatic, as it is for all known Fd complexes (Knaff and Hirasawa 1991): of nine conserved charged residues in the SH3-like domain, only Arg-193 is required, and replacing it by a neutral, acidic or hydrophobic residue lowers the efficiency of Fd-dependent PQ reduction (Yamamoto and Shikanai 2013). The structure of cyanobacterial NDH-1 shows two candidate Fd-binding sites (Laughlin et al. 2019). For the PGR5-dependent pathway, the picture is less complete. In yeast two-hybrid assays, PGRL1 interacts with Fd, PGR5, the PSI subunit PsaD, Cyt *b*₆ and both FNR isoforms, and its interaction with Fd depends on a C-terminal loop (DalCorso et al. 2008). Recombinant PGRL1 binds iron through six conserved cysteines, accepts electrons from Fd reduced by FNR and NADPH, and reduces quinones in a reaction that antimycin A inhibits; on this basis, Hertle et al. (2013) proposed that PGRL1 is the Fd–PQ reductase (FQR). They also quantified the proteins: about 0.70 mmol mol⁻¹ Chl for PGRL1 and about 0.09 for both PGR5 and the NDH subunit NdhH. An involvement of FNR in the antimycin A-sensitive pathway was suggested earlier, because inhibitors of FNR (heparin, dimaleimide) and Fd modified so that it cannot donate electrons to FNR both lowered the FQR activity of pea thylakoids (Cleland and Bendall 1992). Two findings argue against a direct catalytic role of PGR5: it has no cofactor-binding motif (Shikanai 2007), and in leaves of *pgr5*, Fd is more reduced and the Fd-dependent CEF-PSI measured from the oxidation kinetics of Fd is higher than in the wild type (Maekawa et al. 2024). The same *in vivo* measurement shows that this Fd-dependent activity does not require NDH, because it is present in *crr4* (Ohnishi et al. 2023). The identity of the enzyme that oxidizes Fd in the antimycin A-sensitive pathway therefore remains open (Table S2, row 8).

### What the affinity values mean

The only study that measured both pathways in the same system is that of Fisher et al. (2019), who followed the post-illumination fluorescence rise in uncoupled thylakoids supplied with Fd and NADPH. In *A. hybridus* thylakoids with antimycin A, which isolates NDH, the apparent *K*_m_ for Fd was 2.1 ± 0.6 µM; the rate was still rising at the highest concentration tested, 5.5 µM, which was about 70% of the fitted maximum, so this value is a lower estimate. In spinach thylakoids without antimycin A, in which most of the activity is antimycin A-sensitive, the apparent *K*_m_ was 1.3 ± 0.4 µM and the rate saturated near 2 µM. The two values thus differ by less than their uncertainty, and they were obtained in different species. Earlier studies of the antimycin A-sensitive pathway reported larger Fd requirements, but under different conditions. Tagawa et al. (1963) found that cyclic photophosphorylation by broken spinach chloroplasts under anaerobic conditions needed up to ten times more Fd than non-cyclic photophosphorylation. Arnon and Chain (1975) measured cyclic photophosphorylation under 715-nm light in air and found an optimum at about 10 µM Fd, with about ten times more needed under anaerobic conditions; they also showed that the cycle required a small supply of electrons from PSII, because DCMU abolished it. Cleland and Bendall (1992) measured the oxidation of Fd by FQR directly, from the absorbance of Fd at 463 nm, in uncoupled pea thylakoids under anaerobic conditions with DCMU. The rate increased with the concentration of reduced Fd and saturated at about 75 µM. These assays differ from that of Fisher et al. (2019) in three ways. First, they measured ATP synthesis or the oxidation of Fd rather than the reduction of PQ. Second, the redox poise of the system was set differently. In the assays in air, O₂ also accepts electrons from reduced Fd; Arnon and Chain (1975) attributed the lower activity at Fd concentrations above the optimum to this re-oxidation of reduced Fd by O₂ rather than by the cycle. In the anaerobic assay of Cleland and Bendall (1992), Fd was reduced photochemically with deazaflavin before the thylakoids were added. Third, none of them measured NDH. The older values therefore describe the dependence of the antimycin A-sensitive pathway on reduced Fd rather than its affinity for Fd, and they cannot be compared directly with the *K*_m_ values of Fisher et al. (2019). What the old and new studies agree on is that the antimycin A-sensitive pathway is not saturated at 1 µM Fd, the concentration used in many assays (e.g., Fisher et al. 2019).

### Comparison with FNR and the assimilatory enzymes

The affinities of the stromal Fd-dependent enzymes for Fd were reviewed by Knaff and Hirasawa (1991). For FNR, the dissociation constant of the Fd–FNR complex at low ionic strength is below 50 nM in the absence of NADP⁺ and rises to about 0.5 µM when the NADP⁺ site is occupied, a negative cooperativity that helps the oxidized Fd to leave the enzyme after each turnover (Batie and Kamin 1984a, b); at physiological ionic strength, the dissociation constant is 1–10 µM (Knaff and Hirasawa 1991). The dissociation constants for Fd are 0.63–1.0 µM for nitrite reductase, rising to 11 µM when nitrite is bound; 9–14.5 µM for Fd-dependent glutamate synthase, whose *K*_m_ for Fd is about 2 µM; and below 100 nM for Fd–thioredoxin reductase (Knaff and Hirasawa 1991). All these complexes are electrostatic, and FNR, nitrite reductase and glutamate synthase may share a common binding surface on Fd, whereas Fd–thioredoxin reductase may use a different one (Knaff and Hirasawa 1991). Two conclusions follow for CEF-PSI. First, the *K*_m_ values of the two pathways (1–2 µM) are of the same order as the affinities of FNR and the assimilatory enzymes, so neither pathway has a special advantage in the competition for reduced Fd. Second, FNR is the most abundant competitor: in spinach chloroplasts, the molar ratio of Fd to FNR is about 5 : 3 (Böhme 1978) and about 2 : 1 (Terashima and Inoue 1985). As long as NADP⁺ is available, FNR with its high affinity and high turnover will take most of the reduced Fd, and the cyclic pathways receive electrons mainly when NADP⁺ runs short. This is exactly what Tagawa et al. (1963) observed in broken chloroplasts, where cyclic photophosphorylation started at the moment when NADP⁺ was exhausted, and what is seen in leaves, where the Fd-dependent CEF-PSI appears when linear electron flow is saturated (Ohnishi et al. 2023; see below).

### Fd content, its redox state and the supply to CEF-PSI

The amount of Fd in spinach chloroplasts has been measured directly. By quantitative immunoelectrophoresis, Böhme (1978) found Fd : FNR : plastocyanin ≈ 5 : 3 : 4 per P700, with about 80% of the Fd soluble. In chloroplasts isolated from palisade and spongy tissue, Terashima and Inoue (1985) found 6.1 and 2.9 mmol Fd mol⁻¹ Chl against 2.55 and 2.43 mmol P700 mol⁻¹ Chl, that is, about 2.4 and 1.2 Fd per P700, with FNR at about half the amount of Fd; both Fd and FNR were thus lower in the spongy chloroplasts, which receive the weaker and FR-enriched light inside the leaf. Values of two to six Fd per PSI have been compiled for higher plants (Antal et al. 2013). Combined with a chlorophyll concentration of 30–60 mM in the chloroplast (Antal et al. 2013), the Fd contents of 3–6 mmol mol⁻¹ Chl correspond to about 0.1–0.4 mM. Antal et al. (2013) list a somewhat higher value, about 0.6 mM Fd together with 0.3 mM FNR and 0.1 mM PSI, whereas Hanke and Hase (2008) quote about 50 µM; the spread reflects the reference volume and the source of each estimate, but even the lowest value is more than ten times the *K*_m_ values of the two pathways. Experiments that changed the amount of Fd point to the same conclusion. In transplastomic tobacco with about four times the wild-type Fd content, CEF-PSI and NPQ were higher, whereas CO₂ assimilation was not (Yamamoto et al. 2006). In antisense potato with 40–80% of the wild-type Fd, CEF-PSI was also higher (the ratio of the quantum yields of PSI and PSII rose from about 1.3 to above 3), Q_A_ and P700 were more reduced, and plants with less than 40% Fd were not viable (Holtgrefe et al. 2003). That both an increase and a decrease in Fd stimulate CEF-PSI and over-reduce the PQ pool (Hanke and Hase 2008) shows that the amount of Fd acts on the redox poise of the chain rather than as a simple substrate. The redox state of Fd can now be followed in leaves with near-infrared absorbance. At steady state, part of the Fd pool remains reduced, and the reduced fraction rises with light intensity (Satoh et al. 2025). The oxidation rate of Fd that is not accounted for by linear electron flow, taken as the rate of Fd-dependent CEF-PSI, appears when linear electron flow is saturated and the PQ pool is oxidized (Ohnishi et al. 2023), increases with the reduced fraction of Fd and falls when the PQ pool becomes reduced (Satoh et al. 2025), and is higher in *pgr5*, in which Fd is more reduced (Maekawa et al. 2024). These results fit a rate proportional to the product of reduced Fd and oxidized PQ, and they identify the reduced fraction of Fd, not its total amount, as the quantity that limits the supply to CEF-PSI *in vivo*. The isoforms of Fd may add a further level of control. In *Arabidopsis*, Fd2 accounts for 90–99% of the leaf Fd; lowering Fd2 slows linear electron flow, whereas lowering the minor isoform Fd1 increases it and oxidizes the PQ pool, which suggests that Fd1 tends to keep electrons within the chain (Hanke and Hase 2008). In maize, the bundle-sheath isoform FdII reacts more slowly with FNR than the mesophyll isoform FdI and donates more electrons to the PQ pool (Kimata-Ariga et al. 2000). Which isoform feeds each CEF-PSI pathway in C₃ leaves is not known.

### Open questions (Table S3, prediction 5)

(a) The apparent *K*_m_ values of the two pathways have not been measured in the same species with the same Fd preparation; the comparison of Fisher et al. (2019) is between *A. hybridus* and spinach. (b) All *K*_m_ values come from uncoupled thylakoids. Whether the affinity for Fd changes with Δp has not been tested, and Cleland and Bendall (1992) noted that a regulatory effect of adenylates might appear only in coupled thylakoids. (c) The reduced fraction of Fd under FR, and the share of it taken by FNR, have not been measured; this is the quantity that decides how much substrate the cyclic pathways receive when PSII supplies few electrons. (d) Fisher et al. (2019) found inhibition of both pathways by ATP, whereas Cleland and Bendall (1992) found no effect of ATP or ADP up to 1 mM on the antimycin A-sensitive pathway; the reason for the difference is unknown (Table S2, row 3). (e) Whether PGRL1 is the FQR, and what the role of PGR5 is, remain unsettled (Table S2, row 8). (f) The Fd isoform used by each pathway is not known, and the absolute stromal concentration of Fd is known only to within an order of magnitude.

## 3. Supplementary Table S2 and Supplementary Text S2

**Supplementary Table S2.**
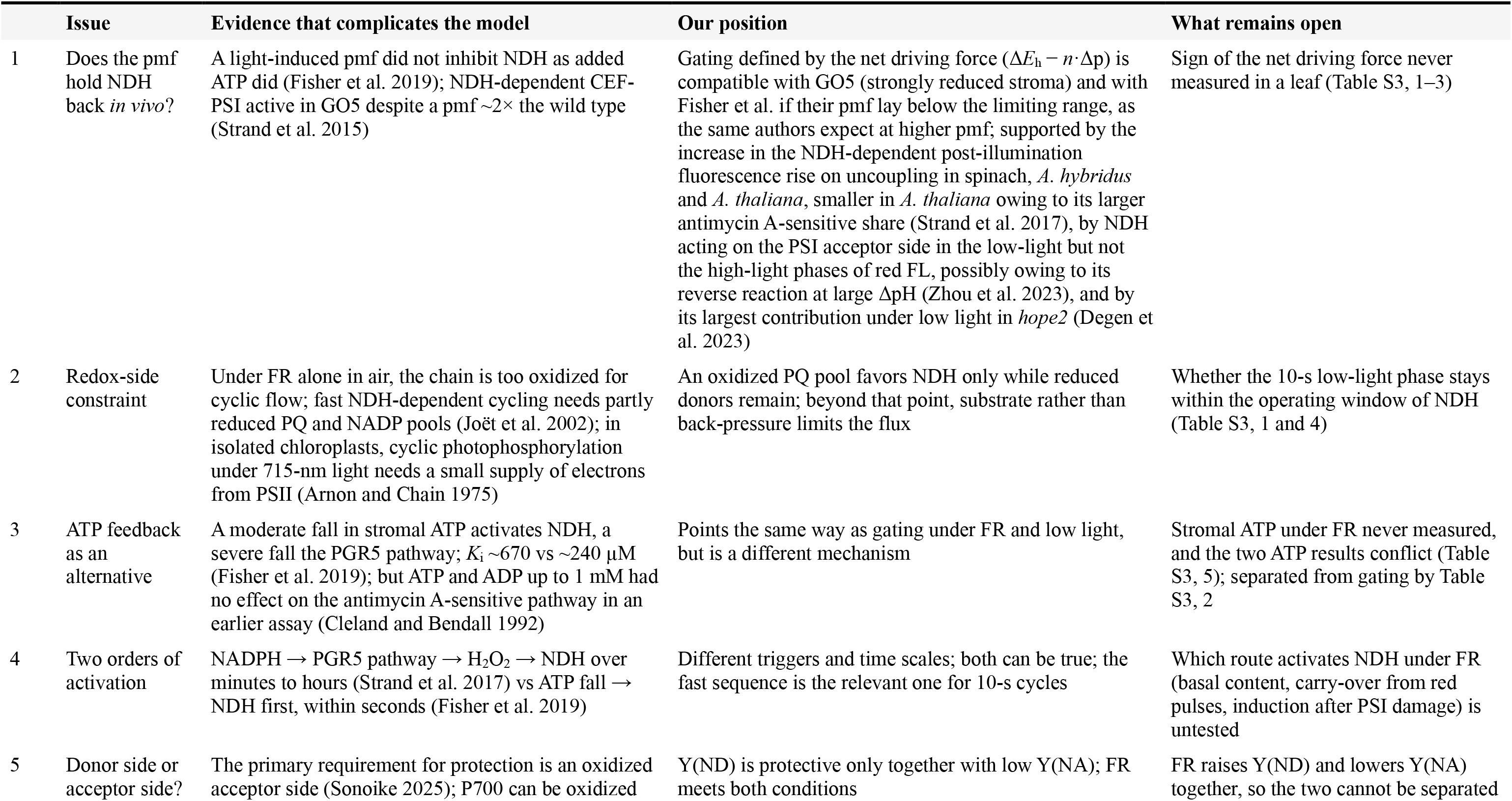
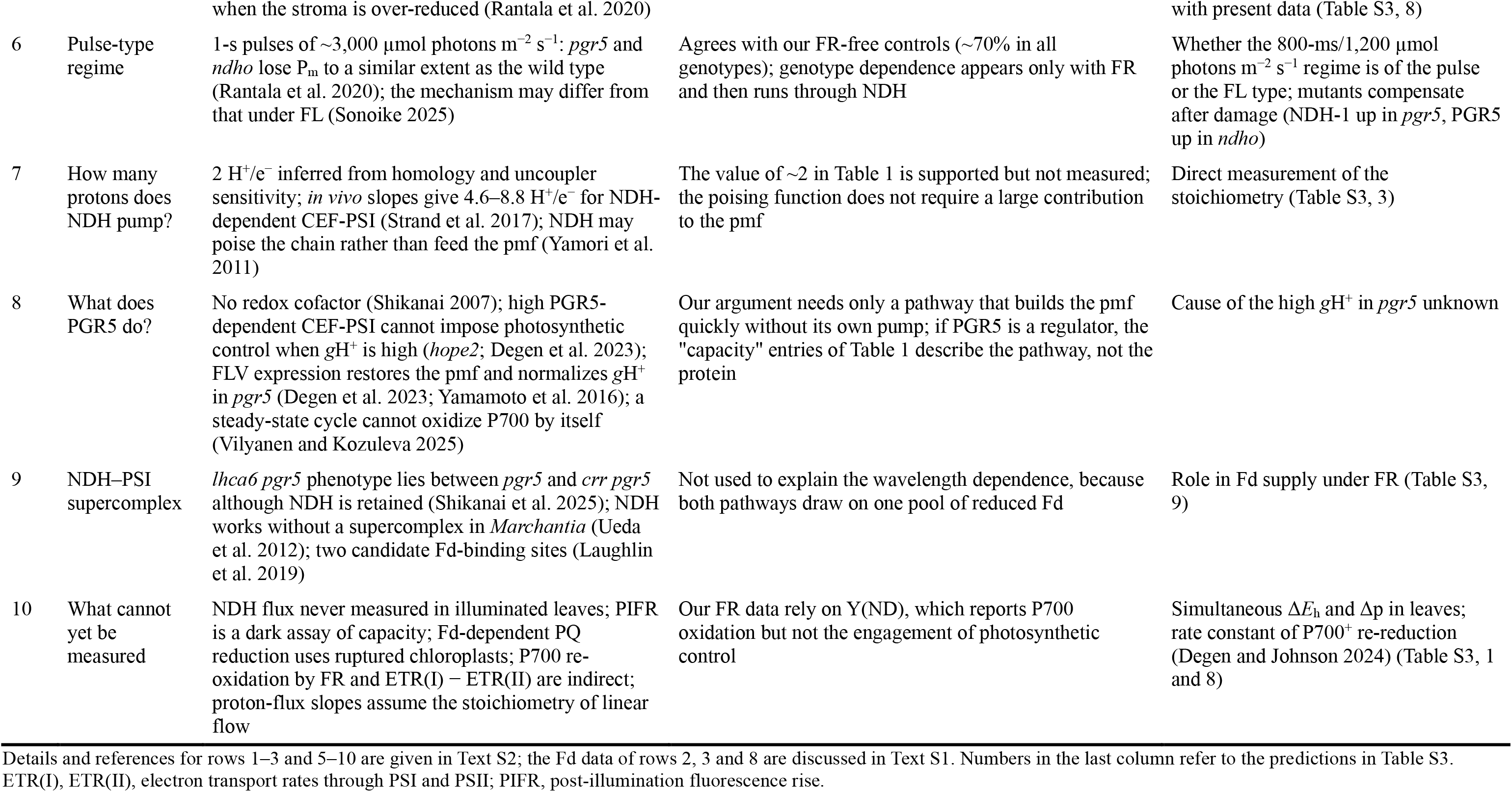
Points of tension in the proposed division of roles between the CEF-PSI pathways.

|  | Issue | Evidence that complicates the model | Our position | What remains open |
|---|---|---|---|---|
| 1 | Does the pmf hold NDH back *in vivo*? | A light-induced pmf did not inhibit NDH as added ATP did (Fisher et al. 2019); NDH-dependent CEF-PSI active in GO5 despite a pmf ~2× the wild type (Strand et al. 2015) | Gating defined by the net driving force (Δ*E*<sub>h</sub> − *n*·Δp) is compatible with GO5 (strongly reduced stroma) and with Fisher et al. if their pmf lay below the limiting range, as the same authors expect at higher pmf; supported by the increase in the NDH-dependent post-illumination fluorescence rise on uncoupling in spinach, *A. hybridus* and *A. thaliana*, smaller in *A. thaliana* owing to its larger antimycin A-sensitive share (Strand et al. 2017), by NDH acting on the PSI acceptor side in the low-light but not the high-light phases of red FL, possibly owing to its reverse reaction at large ΔpH (Zhou et al. 2023), and by its largest contribution under low light in *hope2* (Degen et al. 2023) | Sign of the net driving force never measured in a leaf (Table S3, 1–3) |
| 2 | Redox-side constraint | Under FR alone in air, the chain is too oxidized for cyclic flow; fast NDH-dependent cycling needs partly reduced PQ and NADP pools (Joët et al. 2002); in isolated chloroplasts, cyclic photophosphorylation under 715-nm light needs a small supply of electrons from PSII (Arnon and Chain 1975) | An oxidized PQ pool favors NDH only while reduced donors remain; beyond that point, substrate rather than back-pressure limits the flux | Whether the 10-s low-light phase stays within the operating window of NDH (Table S3, 1 and 4) |
| 3 | ATP feedback as an alternative | A moderate fall in stromal ATP activates NDH, a severe fall the PGR5 pathway; *K*<sub>i</sub> ~670 vs ~240 µM (Fisher et al. 2019); but ATP and ADP up to 1 mM had no effect on the antimycin A-sensitive pathway in an earlier assay (Cleland and Bendall 1992) | Points the same way as gating under FR and low light, but is a different mechanism | Stromal ATP under FR never measured, and the two ATP results conflict (Table S3, 5); separated from gating by Table S3, 2 |
| 4 | Two orders of activation | NADPH → PGR5 pathway → H<sub>2</sub>O<sub>2</sub> → NDH over minutes to hours (Strand et al. 2017) vs ATP fall → NDH first, within seconds (Fisher et al. 2019) | Different triggers and time scales; both can be true; the fast sequence is the relevant one for 10-s cycles | Which route activates NDH under FR (basal content, carry-over from red pulses, induction after PSI damage) is untested |
| 5 | Donor side or acceptor side? | The primary requirement for protection is an oxidized acceptor side (Sonoike 2025); P700 can be oxidized when the stroma is over-reduced (Rantala et al. 2020) | Y(ND) is protective only together with low Y(NA); FR meets both conditions | FR raises Y(ND) and lowers Y(NA) together, so the two cannot be separated with present data (Table S3, 8) |
| 6 | Pulse-type regime | 1-s pulses of ~3,000 µmol photons m<sup>−2</sup> s<sup>−1</sup>: *pgr5* and *ndho* lose P<sub>m</sub> to a similar extent as the wild type (Rantala et al. 2020); the mechanism may differ from that under FL (Sonoike 2025) | Agrees with our FR-free controls (~70% in all genotypes); genotype dependence appears only with FR and then runs through NDH | Whether the 800-ms/1,200 µmol photons m<sup>−2</sup> s<sup>−1</sup> regime is of the pulse or the FL type; mutants compensate after damage (NDH-1 up in *pgr5*, PGR5 up in *ndho*) |
| 7 | How many protons does NDH pump? | 2 H<sup>+</sup>/e<sup>−</sup> inferred from homology and uncoupler sensitivity; *in vivo* slopes give 4.6–8.8 H<sup>+</sup>/e<sup>−</sup> for NDH-dependent CEF-PSI (Strand et al. 2017); NDH may poise the chain rather than feed the pmf (Yamori et al. 2011) | The value of ~2 in Table 1 is supported but not measured; the poising function does not require a large contribution to the pmf | Direct measurement of the stoichiometry (Table S3, 3) |
| 8 | What does PGR5 do? | No redox cofactor (Shikanai 2007); high PGR5-dependent CEF-PSI cannot impose photosynthetic control when *g*H<sup>+</sup> is high (*hope2*; Degen et al. 2023); FLV expression restores the pmf and normalizes *g*H<sup>+</sup> in *pgr5* (Degen et al. 2023; Yamamoto et al. 2016); a steady-state cycle cannot oxidize P700 by itself (Vilyanen and Kozuleva 2025) | Our argument needs only a pathway that builds the pmf quickly without its own pump; if PGR5 is a regulator, the "capacity" entries of Table 1 describe the pathway, not the protein | Cause of the high *g*H<sup>+</sup> in *pgr5* unknown |
| 9 | NDH–PSI supercomplex | *lhca6 pgr5* phenotype lies between *pgr5* and *crr pgr5* although NDH is retained (Shikanai et al. 2025); NDH works without a supercomplex in *Marchantia* (Ueda et al. 2012); two candidate Fd-binding sites (Laughlin et al. 2019) | Not used to explain the wavelength dependence, because both pathways draw on one pool of reduced Fd | Role in Fd supply under FR (Table S3, 9) |
| 10 | What cannot yet be measured | NDH flux never measured in illuminated leaves; PIFR is a dark assay of capacity; Fd-dependent PQ reduction uses ruptured chloroplasts; P700 re-oxidation by FR and ETR(I) − ETR(II) are indirect; proton-flux slopes assume the stoichiometry of linear flow | Our FR data rely on Y(ND), which reports P700 oxidation but not the engagement of photosynthetic control | Simultaneous Δ*E*<sub>h</sub> and Δp in leaves; rate constant of P700<sup>+</sup> re-reduction (Degen and Johnson 2024) (Table S3, 1 and 8) |

Details and references for rows 1–3 and 5–10 are given in Text S2; the Fd data of rows 2, 3 and 8 are discussed in Text S1. Numbers in the last column refer to the predictions in Table S3.
ETR(I), ETR(II), electron transport rates through PSI and PSII; PIFR, post-illumination fluorescence rise.

### Points of tension and open questions

This text expands Table S2. The rows of Table S2 discussed under each heading are given in parentheses; row 4 (two orders of activation) is summarized in the table only. The question of whether P700⁺ is protective in itself, raised in § 5.3, is discussed under row 5.

### Is NDH really held back by pmf *in vivo*? (Table S2, rows 1–3)

The redox–Δp gating hypothesis rests on the thermodynamics of a proton pump, and the evidence that this thermodynamic process operates in a leaf is still indirect. Two observations weigh against it. First, in a later study from the same laboratory, the authors tried to impose a pmf on thylakoids by adding ATP, so that the ATP synthase ran in reverse and pumped protons into the lumen. ATP strongly inhibits NDH, but a light-induced pmf of a similar size does not (Fisher et al. 2019). The authors concluded that ATP directly inhibits NDH. For our purpose, the important point is the other half of the result: a pmf of this size, generated by light, did not hold back NDH. The same authors nevertheless suggested that NDH may become thermodynamically limited at higher pmf (Fisher et al. 2019); therefore, the two observations are not in conflict if the pmf reached in their light experiment was below the range in which back-pressure becomes limiting. Second, NDH-dependent CEF-PSI is clearly active in the H₂O₂-accumulating line GO5, in which the pmf is approximately twice that of the wild type (Strand et al. 2015); we have argued in § 5.3 that this is compatible with gating defined by the net driving force (Δ*E*_h_ − *n*·Δp), but it is not evidence for it. The uncoupler experiments of Strand et al. (2017), by contrast, support gating rather than weaken it. Collapsing the pmf with uncouplers increased the NDH-dependent post-illumination fluorescence rise in thylakoids from spinach, *Amaranthus hybridus* and *Arabidopsis thaliana*. The increase was about twofold in spinach and *A. hybridus* and smaller in *A. thaliana*, which the authors attributed to the larger share of the antimycin A-sensitive pathway, which is not limited by back-pressure, in the fluorescence rise of this species. A further constraint arises from the redox side. Under FR alone, the electron transport chain of tobacco leaves in air is too oxidized for significant cyclic flow, including flow through NDH, and rapid cycling appears only when the PQ and NADP pools are partly reduced (Joët et al. 2002); in isolated chloroplasts, cyclic photophosphorylation under 715-nm light likewise requires a small supply of electrons from PSII, and DCMU abolishes it (Arnon and Chain 1975). Therefore, an oxidized PQ pool thermodynamically favors NDH only as long as enough reduced donors remain; beyond that point, the lack of substrate, not back-pressure, limits the flux. Thus, we present gating as a hypothesis that adds to, and does not replace, the established reasons why NDH contributes little under red high light: its low abundance of approximately 0.02 per PSI, its slow turnover, and its competition with PSII for oxidized PQ (Ermakova et al. 2024a; Table 1). Gating adds a reason why the balance should shift with light quality as well as with irradiance. Two mutant studies point the same way. The first is the *hope2* mutant, in which a defect in the regulation of the ATP synthase is compensated by a high rate of PGR5-dependent CEF: the NDH pathway made its largest contribution under low light, where it is less limited by back-pressure from the pmf, and the authors proposed that competition for electrons at the acceptor side together with thermodynamic constraints on turnover decides which pathway is favored (Degen et al. 2023). The second is the *crr2-2* mutant under red FL (1616/40 | 1 min/5 min): Y(NA) was elevated in the low-light phases but not in the high-light phases, and the authors suggested that a large ΔpH and a reduced PQ pool may favor the reverse reaction of NDH in high light (Zhou et al. 2023). However, no study has simultaneously determined the pmf and redox states of PQ and Fd in a leaf under red and FR; therefore, the sign of the net driving force *in vivo* is unknown. It should also be said that ATP feedback could produce a similar division of roles. In the scheme of Fisher et al. (2019), a moderate fall in stromal ATP activates NDH and a severe fall activates the PGR5-dependent pathway; under FR and low light, the demand for ATP persists while its supply is low. Thermodynamic gating and ATP feedback point in the same direction, but they are different mechanisms that can be separated experimentally.

### Donor side or acceptor side? (Table S2, row 5)

Two recent papers question the direct protective role of P700⁺. Sonoike (2025) argues that the primary requirement for PSI protection is an oxidized acceptor side, that the dissipation of excitation by P700⁺ should have a smaller effect on photoinhibition than the reduction state of the Fe–S clusters, and that FR would aggravate rather than prevent photoinhibition if it merely moved electrons from the donor side to the acceptor side. In his view, the protective agent is the CEF-PSI that FR induces, not FR itself. Rantala et al. (2020) argued that P700 can also become oxidized when the stromal acceptors are over-reduced, so that a rapid oxidation of P700 does not prove that PSI is protected. Taking account of these claims, we propose the roles of P700⁺ in the following way. Y(ND) is an indicator of donor-side limitation, and it is protective only when it increases together with an oxidation of the acceptor side. When FR is added, these two conditions are met: because the electron flux through PSI is small, the sinks downstream of PSI can accept the electrons released by P700 oxidation, so that P700⁺ accumulates while Fd and NADP⁺ remain oxidized. This is exactly the combination of high Y(ND) and low Y(NA) seen under FR (§ 5.3). The protective functions of P700⁺ in this state are twofold. First, excitation that reaches an oxidized reaction center is dissipated without transferring an electron downstream. Photoacoustic measurements on intact leaves show that the extent of P700 oxidation induced by FR is strictly proportional to a positive variable component of heat emission (Bukhov and Carpentier 2003, their Fig. 3c), and that the fraction of reduced P700 matches the fraction of energy stored photochemically (their Fig. 4c); open reaction centers are strong photochemical quenchers, and heat emission rises in proportion to the oxidized P700. Second, charge recombination between P700⁺ and reduced Fe–S clusters can re-oxidize the clusters that would otherwise react with H₂O₂ (Shimakawa and Miyake 2018). Both functions depend on P700 being kept oxidized. When the electron sink downstream of PSI becomes very small, P700 can no longer be kept oxidized even though the pmf is high, because the oxidation of excited P700 becomes rate-limiting (Miyake 2020). The protective effect of FR is therefore conditional: it holds as long as FR keeps the whole chain, and not only P700, in an oxidized state.

The difficulty, which Sonoike (2025) also acknowledges, is that our FR data cannot separate the two. Under red FL, Kono and Terashima (2016) argued that the relief of the acceptor side must come before the rise in Y(ND) (§ 5.2); this is a logical requirement rather than a measured order, and in our FR experiments both parameters were read at the same time points, and at each of them FR had already raised Y(ND) and lowered Y(NA) (Kono et al. 2017); thus, the protection could be attributed to either change. Separating them will require an experiment, for example a series of FR intensities in which Y(ND) rises before Y(NA) falls, or a condition in which P700 is kept oxidized while the acceptor side is deliberately reduced.

### The pulse-type regime (Table S2, row 6)

Rantala et al. (2020) exposed *Arabidopsis* leaves to repeated 1-s pulses of approximately 3,000 µmol photons m⁻² s⁻¹ on a weak background for 2 or 4 h and measured P_m_ after 24 h of recovery. P_m_ fell to approximately 60%, 50%, and 50% after 2 h and to approximately 30%, 15%, and 25% after 4 h in the wild type, *pgr5*, and *ndho*, respectively. Because the relative loss was similar, the authors concluded that neither pathway protects PSI against this type of burst. This conclusion agrees with our own data: in the 800-ms pulse regime of Kono et al. (2017), P_m_ fell to approximately 70% in all genotypes when FR was absent, with no significant difference between them. In pulse regimes, genotype dependence appears only when FR is added, and it then runs through NDH; in the 2-min regime, it runs through PGR5. Two points follow. First, the statement that CEF-PSI does not protect PSI is specific to the pulse regime, and Rantala et al. (2020) accept that PGR5 is required under FL and NDH-1 under rapidly fluctuating light. Sonoike (2025) has argued that photoinhibition by repetitive saturating pulses proceeds by a mechanism different from that under slower FL, with the Fe–S clusters left intact. Second, the fact that FR protects PSI in a pulse regime in which CEF-PSI alone does not is what the masking hypothesis and redox–Δp gating together predict: FR removes the need for a fast, pmf-driven oxidation of P700, and NDH supplies the low-flux recycling that remains necessary. A further complication is that the mutants compensate: after the pulse treatment, NDH-1 increased in *pgr5* and PGR5 in *ndho* (Rantala et al. 2020), so that a mutant measured after photoinhibition is not the same plant as before it.

### How many protons does NDH pump? (Table S2, row 7)

The value of 2 H⁺ pumped by NDH per electron used throughout § 5 is well supported but not measured. This is based on the homology of the membrane arm of NDH with the proton-pumping subunits of respiratory complex I (Laughlin et al. 2019), on the sensitivity of the post-illumination fluorescence rise to uncouplers, which requires a stoichiometry of at least 2 (Strand et al. 2017), and on the slopes of proton flux against PSI electron flux in leaves of the wild-type and the high-CEF mutant *hcef1*. These slopes were 1.8 and 2.75 when fitted freely and 1.6 and 4.7 when forced through the origin (Strand et al. 2017); with 3 H⁺ per electron for linear flow, they correspond to about 4.6 to 8.8 H⁺ per electron for NDH-dependent CEF, that is, to between about 2.6 and 6.8 protons pumped by NDH itself. The *in vivo* data thus show that NDH pumps protons but do not fix the number. Yamori et al. (2011) proposed the alternative view that the flux through NDH is too small to contribute to the pmf and that NDH acts instead by poising the redox state of the intersystem carriers. In our framework, the poising function (§ 5.3) does not depend on a large contribution to the pmf.

### What does PGR5 do? (Table S2, row 8)

The molecular role of PGR5 remains uncertain. PGR5 carries no motif for a redox cofactor and is unlikely to transfer electrons (Shikanai 2007), Fd-dependent PQ reduction is reduced but not abolished in *pgr5* (Munekage et al. 2002), and the mutant’s failure to hold the pmf at a light increase has been interpreted as a loss of control over linear electron transport and over the ATP synthase (Suorsa et al. 2012), an interpretation supported by the elevated proton conductivity of *pgr5* under high light (Rantala et al. 2020; Wang et al. 2015). Two recent studies have sharpened this picture. In *hope2*, a mutant of the ATP synthase γ-subunit with a constitutively high proton conductivity, the pmf and qE are kept at wild-type levels by an increased PGR5-dependent CEF, yet steady-state P700 oxidation is virtually absent and PSI remains susceptible to photoinhibition: a high rate of PGR5-dependent CEF cannot impose photosynthetic control while the proton conductivity remains high (Degen and Johnson 2024; Degen et al. 2023). On the other hand, the expression of the flavodiiron proteins of *Physcomitrium patens* (Hedw.) Mitt. (syn. *Physcomitrella patens* (Hedw.) Bruch & Schimp.) in *pgr5* restores the pmf and P700 oxidation, relieves the acceptor-side limitation (Yamamoto et al. 2016), and normalizes the elevated proton conductivity (Degen et al. 2023), which argues against a direct regulation of the ATP synthase by PGR5; the cause of the high conductivity in *pgr5* is still unknown (Degen et al. 2023). Vilyanen and Kozuleva (2025) add a general point: a cycle at steady state returns exactly as many electrons to PSI as it removes and therefore cannot keep P700 oxidized by itself; its protective effect must come from the protons it moves, either through extra ATP that speeds the Calvin–Benson cycle and relieves the acceptor side, or through lumen acidification and photosynthetic control. Our account in § 5.2 uses both routes and does not require a decision on the molecular role of PGR5. What it requires is only that the PGR5-dependent pathway builds the pmf quickly from a large electron flux without a proton pump of its own; if PGR5 turns out to act mainly as a regulator, the "capacity" entries of Table 1 describe the pathway rather than the protein, and the rescue of *pgr5* by FR is read as FR removing the need for that regulation—the conclusion is unchanged.

### The NDH–PSI supercomplex (Table S2, row 9)

We have not used the supercomplex to explain the wavelength dependence (§ 5.1), but its functional significance remains unresolved. In *lhca6*, which cannot form the supercomplex, the NDH subunits are largely retained, yet the *lhca6 pgr5* double mutant shows a phenotype between *pgr5* and *crr pgr5*, a discrepancy that Shikanai et al. (2025) could not explain; in *Marchantia*, NDH functions without any supercomplex (Ueda et al. 2012). In chilled cucumber leaves, the partial disassembly of the supercomplex accompanies the loss of NDH activity and PSI photoinhibition (Takeuchi et al. 2026; § 1.3). Cyanobacterial NDH-1 has two candidate Fd-binding sites (Laughlin et al. 2019). Whether the supercomplex changes the supply of reduced Fd to NDH or its local redox environment under FR is a legitimate question (Table S3, prediction 9).

### What cannot yet be measured (Table S2, row 10)

Finally, the flux through NDH has never been directly measured in illuminated leaves. Every available method carries an assumption: the post-illumination fluorescence rise is measured in the dark, when Δp has collapsed and NDH is least constrained, so that it reports capacity rather than the flux in the light (Strand et al. 2017); Fd-dependent PQ reduction is measured in ruptured chloroplasts; the re-oxidation of P700 by FR (Shikanai et al. 2025) and the difference between the electron transport rates of PSI and PSII are indirect; and the proton-flux slopes of Strand et al. (2017) assume fixed stoichiometries for linear flow. The two variables of the gating hypothesis, Δ*E*_h_ and Δp, have not been measured together under red and FR light, and the ATP level under FR has not been measured at all. Our FR data rely on Y(ND), which reports the extent of P700 oxidation but not the engagement of photosynthetic control; the rate constant of P700⁺ re-reduction is a more direct indicator (Degen and Johnson 2024). These gaps are addressed in Text S3.

## 4. Supplementary Table S3 and Supplementary Text S3

**Supplementary Table S3.** Testable predictions of the masking and redox–Δp gating hypotheses.

|  | Prediction | Experiment | Result that supports | Result that refutes |
|---|---|---|---|---|
| *Testing redox–Δp gating directly* |  |  |  |  |
| 1 | The sign of the net driving force (Δ*E*<sub>h</sub> − *n*·Δp) differs between red and FR light | Leaves: pmf from ECS and redox states of P700, PC and PQ under red vs red + FR; driving force calculated as in Strand et al. (2017) | Net driving force more positive under FR, mainly through PQ oxidation; the low-light phase stays within the operating window of NDH | Net driving force equally positive in both regimes |
| 2 | The NDH-dependent rate falls as the imposed Δp rises; the antimycin A-sensitive rate does not | Coupled thylakoids under FR only; Δp set without ATP (acid–base transition, K<sup>+</sup> diffusion potential, ATP synthase inhibition); PIFR or P700<sup>+</sup> re-reduction, ± antimycin A or *pgr5*; PQ redox state set by the FR:red ratio | Monotonic decline with Δp; increase as PQ is oxidized | No Δp dependence |
| 3 | NDH carries a larger share of cyclic flux under FR in leaves | Proton flux (ECS) vs PSI electron flux (Strand et al. 2017) under red vs FR, ± antimycin A, and in *crr*; KEA3 effects on DIRK taken into account | H<sup>+</sup>/e<sup>−</sup> ratio rises with FR, survives antimycin A, and is absent in *crr* | No change, or a change lost with antimycin A |
| 4 | NDH runs forward only within an operating window: upper limit set by the pmf, lower limit by the supply of reduced Fd and NADPH | NDH share (as in 3) across irradiances under FR-supplemented light in plants of known growth irradiance | Share rises while the pmf builds and falls once it saturates (~140 µmol photons m<sup>−2</sup> s<sup>−1</sup> for plants grown at 50; Wang et al. 2015); share falls again when the red component is lowered until the PQ pool is almost fully oxidized | Monotonic increase with irradiance; no decline at the oxidized end |
| 5 | Fd supply, Fd affinity and the ATP level favor NDH under FR | Apparent *K*<sub>m</sub> for Fd (uncoupled, different species: ~2 µM NDH, ~1 µM antimycin A-sensitive; Fisher et al. 2019) in the same species and in coupled thylakoids ± antimycin A, *crr*, *pgr5*; reduced fraction of Fd and Fd-dependent CEF-PSI from Fd oxidation kinetics in leaves under FR (Ohnishi et al. 2023; Satoh et al. 2025); stromal ATP under FR-supplemented low light | *K*<sub>m</sub> changes with Δp; reduced Fd small but not zero under FR, and Fd-dependent CEF-PSI rises with FR in the wild type and *pgr5* but not in *crr*; ATP in the range where NDH is active but the PGR5 pathway is inhibited | *K*<sub>m</sub> independent of Δp; Fd-dependent CEF-PSI rises with FR in *crr*; ATP outside that range |
| *Testing the masking hypothesis and the role of P700<sup>+</sup>* |  |  |  |  |
| 6 | The division of roles is set by the light regime | Full matrix: 2-min regime (Kono and Terashima 2016) and pulse regime (Kono et al. 2017) × *pgr5*, *crr* × ± FR | 2-min regime: FR rescues *pgr5* only partially; pulse regime: removing FR abolishes the genotype dependence | FR rescues *pgr5* fully in both regimes |
| 7 | FR protection bypasses PGR5 and requires NDH | *pgr5 crr4-3* double mutant under FR-supplemented FL (young leaves, low growth light) | FR rescues *pgr5* but not the double mutant | FR rescues the double mutant |
| 8 | P700 is oxidized by different means in the two regimes | Series of FR intensities or FR:red ratios; P<sub>m</sub> loss vs residual Y(NA) and Y(ND); rate constant of P700<sup>+</sup> re-reduction measured alongside Y(ND) (Degen and Johnson 2024) | P<sub>m</sub> loss tracks Y(NA) (or both); FR raises Y(ND) without slowing re-reduction, red high light raises it by slowing re-reduction | FR slows P700<sup>+</sup> re-reduction as red high light does |
| *Structure* |  |  |  |  |
| 9 | The supercomplex matters only if it supplies Fd to NDH | *lhca5 lhca6* double mutant and Lhca6 domain-swap lines (Otani et al. 2017) under FR-supplemented FL | Wild-type behavior (if the NDH content is adequate) | *crr*-like behavior |
| *Ecological and applied questions* |  |  |  |  |
| 10 | FR and a low cytochrome *b*<sub>6</sub>*f* content protect PSI by the same route | Intermediate-Rieske tobacco (Kono et al. 2025) ± FR-supplemented FL; growth under canopy-like light | Combined treatment protects PSI with a smaller loss of capacity than either alone; intermediate lines show intermediate protection | No additivity |
| 11 | The NDH growth defect under canopy-like FL depends on FR | *crr*, *pgr5* and wild type under canopy-spectrum rapid FL with FR present or filtered out; PSI photoinhibition and biomass measured together | Defect larger with FR and then accompanied by P<sub>m</sub> loss; removing FR shifts the dependence toward PGR5 | Defect independent of FR |

Predictions are grouped by the question they address; each is described in full in Text S3. DIRK, dark-interval relaxation kinetics; ECS, electrochromic shift; PC, plastocyanin; PIFR, post-illumination fluorescence rise.

### Experiments to test the masking and redox–Δp gating hypotheses

The hypotheses presented in § 5.3 make predictions that can be tested using existing methods. We list them here, grouped by the question they address, and state for each what result would support or refute the hypothesis. The numbers correspond to those in Table S3.

### Testing redox–Δp gating directly (Table S3, predictions 1–5)

(1) *The sign of the net driving force (ΔE_h_ − n·Δp) under red and FR light.* The central prediction is that FR shifts the thermodynamic balance of NDH toward the forward reaction. This can be tested in leaves by measuring the pmf from the electrochromic shift and the redox states of P700, plastocyanin, and the PQ pool simultaneously under red light and under red light supplemented with FR, and by calculating the driving force from the measured redox states, as Strand et al. (2017) did for isolated thylakoids. We predict that the driving force for the reduction of PQ by reduced Fd exceeds *n*·Δp by a larger margin under FR than under the high-light phase of red FL, and that the difference arises mainly from the oxidation of the PQ pool rather than from a change in total pmf (Kono et al. 2020). The same measurements should show whether, in the 10-s low-light phase, the PQ pool stays within the operating window of NDH (§ 5.3): oxidized enough to keep Δ*E*_h_ large, but not so oxidized that the supply of reduced donors limits cycling (Joët et al. 2002). If the net driving force is found to be equally positive in both regimes, gating cannot explain the wavelength dependence, and the explanation must be sought in capacity, ATP feedback, or activation.
(2) *The Δp dependence of NDH in coupled thylakoids without competition from linear flow.* The uncoupler experiments of Strand et al. (2017) showed that NDH is sensitive to Δp, but uncouplers abolish the pmf rather than set it to graded values. In coupled thylakoids illuminated with FR only, so that linear flow is absent, the pmf can be set to defined values without ATP. This can be done by an acid–base transition in which a permeant organic acid carries protons into the lumen (Jagendorf and Uribe 1966; Uribe and Jagendorf 1967a, b), by a K⁺ diffusion potential, or by inhibition of the ATP synthase. The NDH-dependent reduction of PQ can then be followed by the post-illumination fluorescence rise or by the re-reduction of P700⁺, with the PGR5-dependent component removed by antimycin A or using *pgr5*. We predict that the NDH-dependent rate falls monotonically as the imposed Δp increases, whereas the antimycin A-sensitive rate does not. Imposing the pmf without ATP is essential because ATP inhibits both pathways directly (Fisher et al. 2019). In the same system, the PQ redox state can be set by the ratio of FR to red light, and the NDH-dependent rate should increase as the pool is oxidized.
(3) *The effective flux in leaves under competition from linear flow.* The proton-flux method of Strand et al. (2017), which compares the proton flux from the electrochromic shift with the electron flux through PSI, can be applied to leaves under red and FR-supplemented light, with and without antimycin A, and in the *crr* mutants. If NDH carries a larger share of the cyclic flux under FR, the ratio of proton to electron flux should increase with FR in the wild type, this increase should survive antimycin A, and it should be absent in *crr* mutants. The dark-interval relaxation kinetics on which this method rests must be interpreted with care: the thylakoid K⁺/H⁺ antiporter KEA3, which exports protons from the lumen in exchange for K⁺ and is activated when the light decreases and the stromal energy state falls (Armbruster et al. 2014, 2016; Uflewski et al. 2024), can alter the relaxation of the electrochromic shift during the dark interval and thus the proton flux and the ΔpH/ΔΨ partitioning derived from it. The difference between the maximal NDH activity measured in experiment (2) and the effective flux measured in experiment (3) would quantify, for the first time, how much of the capacity of NDH is actually used in a leaf under a given light quality.
(4) *The operating window of NDH.* § 5.3 proposed that NDH runs forward only within an operating window, whose upper limit is set by the pmf and whose lower limit is set by the supply of reduced Fd and NADPH. The upper limit can be tested by measuring the NDH-dependent share of cyclic flux, as in experiment (3), across a series of irradiances under FR-supplemented light in plants with known growth irradiance. We predict that the NDH share increases with irradiance while the pmf is still building and falls once the pmf saturates, which in plants grown at 50 µmol photons m⁻² s⁻¹ occurs at approximately 140 µmol photons m⁻² s⁻¹ (Wang et al. 2015). A monotonic increase would argue against an upper limit set by the pmf. The lower limit can be tested in the same series by lowering the red component of the light until the PQ pool is almost fully oxidized: the NDH share should then fall again, as cycling did in tobacco leaves under FR alone (Joët et al. 2002).
(5) *Fd supply, Fd affinity and ATP under FR.* The apparent *K*_m_ values for Fd of the two pathways (about 2 µM for NDH and about 1 µM for the antimycin A-sensitive pathway) were obtained in uncoupled thylakoids and in different species (Fisher et al. 2019; Text S1). Measuring them in the same species and in coupled thylakoids, with and without antimycin A and in *crr* and *pgr5*, would show whether the affinity for Fd itself changes with Δp, which would be a second route by which the redox poise could favor one pathway. Because both pathways compete with FNR and the other Fd-dependent enzymes for reduced Fd on roughly equal terms (Text S1), the quantity that limits the supply to CEF-PSI *in vivo* is the reduced fraction of Fd rather than its total amount. This fraction can now be measured in leaves from the near-infrared absorbance of Fd (Ohnishi et al. 2023; Satoh et al. 2025). Ideally, the decay of reduced Fd would be resolved into the rate constants of its individual sinks, so that the share of each pathway could be read directly; at the present signal-to-noise ratio of the Fd signal this is not feasible, and the comparison has to rely on mutants and inhibitors. We predict that under FR-supplemented low light it is small but not zero, and that the Fd-dependent CEF-PSI measured from the oxidation kinetics of Fd rises with FR in the wild type and in *pgr5* but not in *crr* mutants. In parallel, the stromal ATP level under FR-supplemented low light has never been measured; if it falls into the range in which NDH is active but the PGR5-dependent pathway is inhibited (Fisher et al. 2019), the ATP feedback and the thermodynamic gating would be acting together, and their contributions would have to be separated by experiment (2). The discrepancy between the inhibition by ATP found by Fisher et al. (2019) and its absence in the assay of Cleland and Bendall (1992) should be resolved in the same system (Table S2, row 3).

### Testing the masking hypothesis and the role of P700⁺ (Table S3, predictions 6–8)

(6) *Regime × FR.* Our two FL regimes have never been compared under the same conditions. We predict that in the 2-min regime of Kono and Terashima (2016), in which the high-light phase is long and the PQ pool is reduced throughout it, FR will rescue *pgr5* only partially, because the pmf-driven oxidation of P700 during the high-light phase still depends on the PGR5-dependent pathway; and that in the pulse regime of Kono et al. (2017), removing FR abolishes the genotype dependence, as observed. The full matrix of two regimes, two mutants, and FR present or absent would test whether the division of roles is set by the light regime, as the model claims.
(7) *The pgr5 crr double mutant under FR.* If FR protection runs through NDH and bypasses PGR5, FR should fail to rescue the *pgr5 crr4-3* double mutant, whereas it should rescue *pgr5*. The double mutant grows poorly (Munekage et al. 2004); therefore, the experiment must use young leaves under low growth light, but the prediction is unambiguous.
(8) *Separating Y(ND) from Y(NA).* Under FR, the two parameters change together (Text S2). A series of FR intensities or FR-to-red ratios in which Y(ND) rises before Y(NA) falls would separate them; the loss of P_m_ should then correlate with the residual Y(NA) rather than with Y(ND) if the acceptor side is decisive, and with both if P700⁺ contributes in its own right. In the same experiments, the rate constant of P700⁺ re-reduction should be measured alongside Y(ND) (Degen and Johnson 2024). We predict that FR raises Y(ND) in the low-light phase without slowing P700⁺ re-reduction, that is, without engaging photosynthetic control, whereas red high light raises Y(ND) by slowing it. This would provide the first direct evidence that P700 is oxidized by different means in the two regimes (Table 2).

### Structure (Table S3, prediction 9)

(9) *NDH–PSI supercomplex.* Two kinds of plants lack the supercomplex but retain NDH: the *lhca5 lhca6* double mutant and the domain-swap lines of Otani et al. (2017), in which the stromal loop of Lhca6 is replaced. These should be exposed to FR-supplemented FL. If the supercomplex is needed only for the stability of NDH, these lines should behave like the wild type, as long as their NDH content is adequate. If the supercomplex is needed for the supply of reduced Fd to NDH, they should behave like *crr* mutants under FR.

### Ecological and applied questions (Table S3, predictions 10 and 11)

(10) *Trading capacity for protection.* Tobacco with a low Cyt *b*₆*f* content maintains PSI integrity under extreme pulses, but at the cost of photosynthetic capacity (Kono et al. 2025). If FR and a low Cyt *b*₆*f* content protect PSI by the same route (§ 5.4), a moderate reduction in Rieske protein combined with FR-supplemented FL should protect PSI with a smaller loss of capacity than either treatment alone, and plants with intermediate Rieske content, which have not yet been tested for photoinhibition, should show intermediate protection. The growth of these lines under a canopy-like light regime would show whether the trade-off can be shifted in practice or not.
(11) *Light quality as an independent variable in the field.* The growth defect of NDH mutants under FL that reproduced the irradiance, but not the spectrum, of a maize canopy (Theeuwen et al. 2022, preprint) and the protection of PSI by FR in the laboratory (Kono et al. 2017) have not been connected. Why NDH matters under such rapid fluctuations even without FR is itself open, and our framework suggests a hypothesis. A canopy regime spends most of its time at low irradiance, and its bright spots are too short for the pmf to reach the level that stops NDH; the role of NDH under this regime would then be the one described in § 5.3 and in Table S3, prediction 4, namely to keep the intersystem chain poised and to supply ATP during the low-light periods, rather than to protect PSI directly. This agrees with the observation that the Burren *ndhG* variant affects PSII efficiency only below about 350 µmol photons m⁻² s⁻¹ (Theeuwen et al. 2022, preprint), and it predicts that the growth defect of the NDH mutants under such a regime is accompanied by a more reduced PQ pool and a lower pmf in the low-light periods, but not necessarily by a loss of P_m_. The experiment that connects them is to grow *crr*, *pgr5*, and wild-type plants under a rapidly fluctuating regime that reproduces a canopy spectrum, with FR either present or removed by a filter, and to measure PSI photoinhibition and biomass together. We predict that the growth defect of the NDH mutants is larger with FR than without, that it is accompanied by a loss of P_m_ when FR is present, and that removing FR shifts the dependence from NDH toward PGR5. This experiment would also test the suggestion of § 9.2 that red LED regimes have overestimated the role of PGR5 in nature.

If experiments (1) to (5) fail to show a wavelength-dependent thermodynamic constraint on NDH, the redox–Δp gating hypothesis should be abandoned, and the division of roles would have to be explained by capacity, ATP feedback, and activation alone. The masking hypothesis would survive that outcome, and so would the observations that rest on it: that FR oxidizes P700 by the excitation balance, that this removes the specific need for PGR5, and that NDH provides the low-flux recycling under FR. They do not depend on how NDH is gated, but only on the fact that NDH is the pathway left when a fast, pmf-driven oxidation of P700 is no longer required.

